# Effects of nicotine compared to placebo gum on sensitivity to pain and mediating effects of peak alpha frequency

**DOI:** 10.1101/2023.08.11.552723

**Authors:** Samantha K. Millard, Alan K.I. Chiang, Peter Humburg, Nahian Chowdhury, Raafay Rehan, Andrew J. Furman, Ali Mazaheri, Siobhan M. Schabrun, David A. Seminowicz

## Abstract

Recent research has linked individual peak alpha frequency (PAF) to pain sensitivity, but whether PAF alterations can influence pain remains unclear. Our study investigated the effects of nicotine on pain sensitivity and whether pain changes are mediated by PAF changes. In a randomised, double-blind, placebo-controlled experiment, 62 healthy adults (18–44 years) received either 4 mg nicotine (n=29) or placebo gum (n=33). Resting state EEG and pain ratings during prolonged heat and pressure models were collected before and after nicotine intake. The nicotine group showed a small decrease in heat-pain ratings compared to placebo group when controlling confounders, and a small increase in PAF across the scalp from pre- to post-gum, both with and without confounder adjustment. These effects were most pronounced in the central-parietal and right-frontal electrodes. However, mediation analysis did not support the notion that PAF changes mediate nicotine’s effects on pain sensitivity. While a growing body of literature supports a link between PAF and both acute and chronic pain, further work is needed to understand the mechanisms of this link.

## 1 Introduction

Chronic pain is a highly prevalent and multifaceted health issue, which produces significant physical, emotional, and economic burden to those in pain, their families and friends, health care systems, and society [1–3]. Identifying modifiable factors that influence pain sensitivity could be a key step in reducing the presence and burden of chronic pain [4–6]. Individual peak alpha frequency (PAF) is an electro-physiological brain measure that shows promise as a biomarker of pain sensitivity [7–13]. Although the number of studies investigating PAF as a predictor of pain is increasing [14–18], no experiments have manipulated PAF while also measuring pain, which has limited our understanding of whether, and how, PAF is mechanistically linked to pain sensitivity.

PAF refers to the dominant oscillatory frequency in the 8–12 Hz alpha range. As alpha oscillations are believed to represent periodic inhibitory processes that direct temporal aspects of sensory information processing [19,20], it is thought that PAF reflects the rate at which sensory information is sampled from the external environment [21–23]. In the context of pain, slower PAF speed has been associated with the presence and duration of chronic pain [17,24–28], while in healthy, pain-free individuals, slower PAF has been associated with higher pain sensitivity in experimental pain models [8–10] and acute post-surgical settings [11].

While previous studies suggest that PAF could be used as either a biomarker for disease progression or as a predictive biomarker for pain sensitivity, critical gaps remain in our understanding of how PAF is mechanistically linked to pain. Specifically, no study has examined whether the manipulation of PAF alters pain sensitivity. Addressing this question has significant implications, as it would determine i) whether PAF is causally implicated in pain processing, and ii) whether manipulation of PAF has relevance as a method of clinical pain modulation. Several methods have been used to alter individual PAF, including brain stimulation [29–31], exercise [32–34], visual stimulation [35,36], cognitive tasks [37], and nicotine [38–40]. Because evidence suggests that nicotine can modulate PAF, where both nicotine and smoking increase PAF speed [38,41–48], we chose nicotine to assess our aim of whether changes in PAF mediate changes in pain in a ‘mediation by design’ approach [49]. In addition, given evidence that nicotine may increase experimental pain thresholds and tolerance [50–54], nicotine could also influence pain ratings during tonic pain. Thus, the primary questions addressed in this study were whether nicotine can: i) increase peak alpha frequency (PAF), ii) reduce pain sensitivity, and iii) determine whether changes in pain sensitivity are mediated by changes in PAF speed.

A secondary question pertains to the nature of the PAF–pain relationship. A substantial proportion of the current literature suggests that slower PAF speed is associated with higher pain sensitivity [8–11,17,24–27]. However, conflicting findings exist, with some studies reporting no relationship between PAF and chronic pain [55–58], or an opposite relationship where faster PAF is associated with higher pain sensitivity in experimental pain settings [7,15]. Divergent results could be due to variations in the relationship between PAF and different types of pain, or the dependence of this relationship on individual factors. For example, it was recently suggested that PAF–pain relationships may only be observed in neuropathic pain or expressed differently in males and females [17]. One way to address this issue is to investigate PAF across multiple pain models [12]; currently the PAF–pain relationship has been studied using thermal [7,8,10], musculoskeletal [9,15], and surgical [11] pain, but never with multiple pain modalities included in the same study. Thus, a secondary question was whether baseline PAF, measured in a pain-free resting state, correlated with pain sensitivity during a model of heat pain (i.e. PHP), which has been previously assessed and is negatively correlated with PAF [10], and a model of pressure pain (i.e. cuff pressure algometry; CPA [59–63]) that has not been assessed in relation to PAF. Applying two experimental pain models also allows assessment of the generalisability of nicotine’s effects on pain and the potential mediating effects of PAF, including whether these effects are consistent across different types of pain.

Using a randomised, placebo-controlled, double-blind, parallel design, a sample of healthy, pain-free, non-smoking participants had their resting state PAF measured before and after two models of prolonged pain (i.e. CPA and PHP; pre-gum). Participants then chewed either nicotine or placebo gum, after which their resting state PAF was re-assessed before and after the two prolonged pain models (i.e. CPA and PHP; post-gum). We predicted that: 1. nicotine gum would increase PAF speed in comparison to placebo gum; 2. nicotine gum would decrease pain ratings in comparison to placebo gum; 3. decreases in pain ratings following nicotine gum would be mediated by increases in the speed of PAF; and 4. faster baseline PAF would be associated with lower baseline pain ratings. Based on a pre-registered analysis plan (https://osf.io/pc4rq/), predictions 1–3 were tested using two-wave latent change score (2W-LCS) mediation models [64–66], while prediction 4 was tested using correlation analysis.

## 2 Results

The statistical analysis plan for this study was pre-registered on Open Science Framework (OSF: https://osf.io/pc4rq/) in July 2022.

### 2.1 Sample Characteristics

The participant sample included 62 healthy adults (26.45 ± 7.28; range: 18–44 years), without neurological or psychological disorders. Fourteen participants self-identified as North-West European (22.58%), 12 as South-East Asian, (19.36%), and eight reported mixed ancestries (12.90%; see Supplementary Material 1 for a full list of self-reported ancestry). There were 32 females (24.53 ± 6.52; range: 18–43 years) and 30 males (28.5 ± 7.58; range: 19–44 years); no participants reported their sex at birth as inter-sex. Thirty-one individuals identified as women, 30 as men, and 1 as non-binary.

Regarding pain in the past two weeks, 20 participants reported “None” (nicotine: n = 10; placebo: n = 10), 16 reported “Very Mild” (nicotine: n = 14; placebo: n = 12), 15 reported “Mild” (nicotine: n = 5; placebo: n = 10), one reported “Moderate” (nicotine: n = 1, placebo: n = 0), and no participants reported “Severe” pain in the past two weeks. Participants were successfully blinded to group allocation (Supplementary Material 2) and baseline demographics are displayed in Table 1. Depressive symptoms ranged from 0–4 (out of a maximum of 6), anxiety from 0–6 (out of a maximum of 6), perceived stress from 2–18 (out of a maximum of 40), and sleep quality from 2–10 (out of a maximum of 10; i.e. “excellent sleep quality”). Following CONSORT recommendations [67,68], no baseline statistics are reported.

**Table 1:** Mean ± standard deviations (SD) for baseline participant characteristics split by gum group.

| Group | N = 62 | Nicotine (n = 29) | Placebo (n = 33) |
| --- | --- | --- | --- |
| Age | 26.45 ($\pm 7.28$) | 26.18 ($\pm 8.06$) | 26.39 ($\pm 6.65$) |
| Weight (kg) | 68.16 ($\pm 13.42$) | 68.9 ($\pm 13.00$) | 67.5 ($\pm 13.90$) |
| Height (cm) | 172.29 ($\pm 10.08$) | 172.40 ($\pm 10.00$) | 172.20 ($\pm 10.30$) |
| Depressive | 0.53 ($\pm 0.97$) | 0.62 ($\pm 1.10$) | 0.46 ($\pm 0.75$) |
| Anxiety | 0.69 ($\pm 1.06$) | 0.76 ($\pm 1.30$) | 0.64 ($\pm 0.82$) |
| Stress | 9.82 ($\pm 3.81$) | 8.93 ($\pm 4.04$) | 10.61 ($\pm 3.48$) |
| Sleep quality | 6.97 ($\pm 1.77$) | 6.79 ($\pm 1.86$) | 7.12 ($\pm 1.71$) |
| Heat pain threshold | 41.27 ($\pm 2.59$) | 40.97 ($\pm 2.79$) | 41.54 ($\pm 2.41$) |
| Pressure threshold | 293.92 ($\pm 121.21$) | 318.70 ($\pm 111.25$) | 272.14 ($\pm 127.00$) |
| Alpha power (8–12 Hz) | 26.24 ($\pm 7.02$) | 25.89 ($\pm 7.64$) | 26.55 ($\pm 6.54$) |

*Note.* PAF = peak alpha frequency; PHP = phasic heat pain; CPA = cuff pressure algometry.

### 2.2 Effects of nicotine gum on alpha oscillations and prolonged pain ratings

Mediation analysis was conducted to assess the effects of nicotine on PAF and pain ratings, and whether change in PAF mediated change in pain.

ANCOVA-equivalent two-wave latent change score (2W-LCS) models (i.e. with autoregressive and coupling parameters) and difference score 2W-LCS models (i.e. without auto-regressive and coupling parameters) were constructed for mean PHP and CPA pain outcomes separately (see Methods for further details). As no significant difference was found between model fits (see Supplementary Materials 3.1 and 3.2), the difference score 2W-LCS models are reported here; whilst the full ANCOVA-equivalent 2W-LCS models are presented in Supplementary Material 3.1.

#### Nicotine gum modulated global PAF (8–12 Hz) and PHP ratings, but change in PHP ratings were not mediated by change in PAF

The PHP model consisted of five repetitions of a 40 second heat stimulation to the inside left forearm at a fixed 46 °C (see Methods), which was most frequently described as moderate/severe hot, sharp, and shooting pain (Figure 1, plot A). Raw time courses of pain ratings during PHP are displayed in Figure 1 (plot C), from which the mean pain during max heat stimulation was calculated (Table 2, Figure 1, plot E).

**Figure 1:**
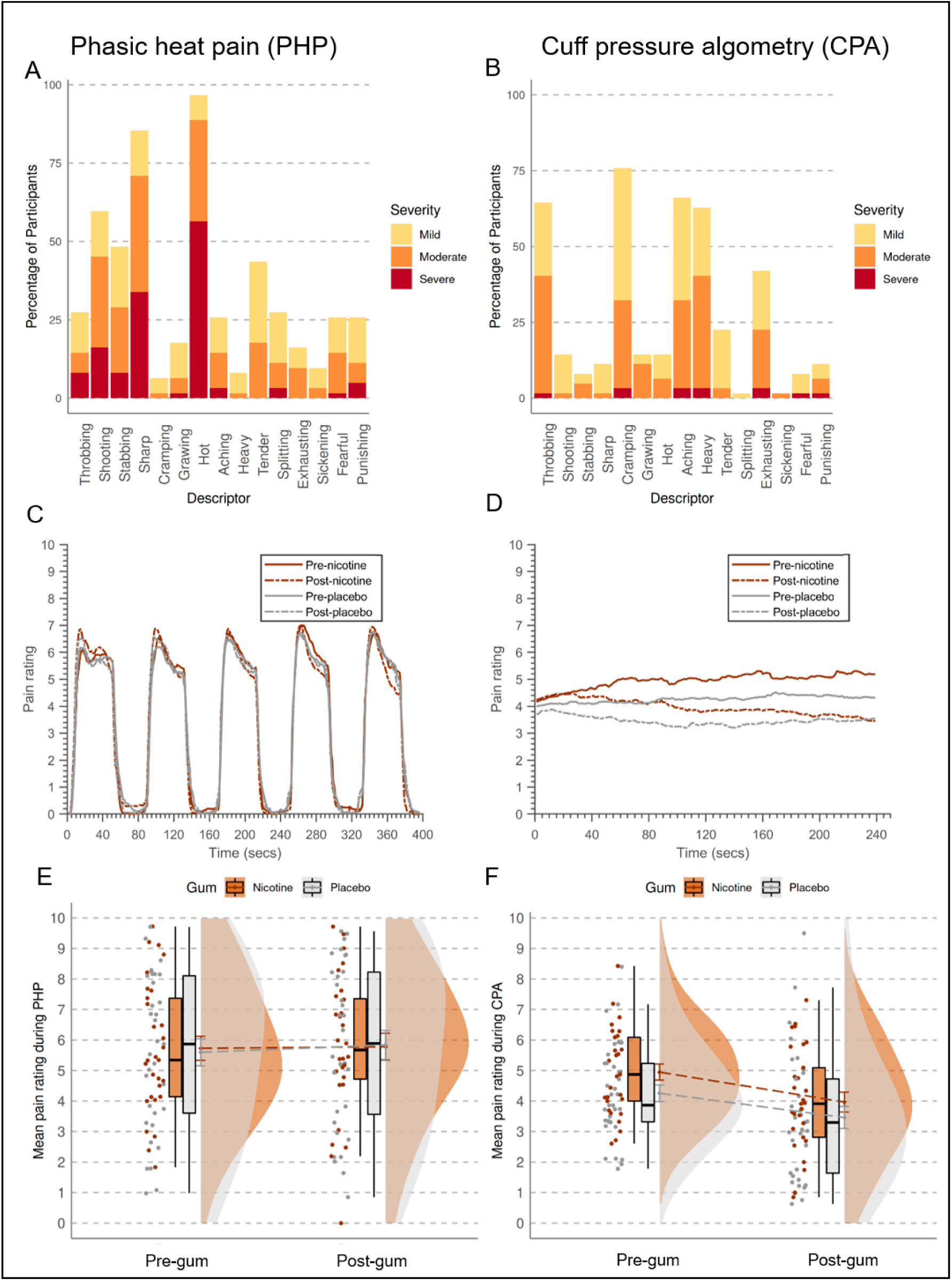
Percentage of participants reporting each category descriptor of pain on the McGill pain questionnaire for A) phasic heat pain (PHP) and B) cuff pressure algometry (CPA) at baseline. Average pain time courses split into nicotine (orange, n = 29) and placebo (grey, n = 33) gum groups for C) PHP and D) CPA models of prolonged pain. Raincloud plots of mean pain ratings during E) PHP and F) CPA models of prolonged pain.

**Table 2:** Mean ± standard deviations (SD) for pain ratings during phasic heat pain (PHP) and cuff pressure algometry (CPA), for pre- and post-gum, for the whole sample and separated by nicotine and placebo groups. Note that post-gum PHP ratings could not be collected for two participants. Therefore, the full sample of 62 participants were used for baseline correlation analyses of PHP and CPA, as well as for CPA mediation models, but list-wise deletion to ensure full pre-post samples meant that only 60 participants were used for PHP mediation models.

| Gum<br>group | n | PHP |  | n | CPA |  |
| --- | --- | --- | --- | --- | --- | --- |
|  |  | Pre-gum | Post-gum |  | Pre-gum | Post-gum |
| Total | 62 | 5.66 ($\pm 2.34$) | | 62 | 4.58 ($\pm 1.53$) | 3.70 ($\pm 1.93$) |
| | 60 | 5.66 ($\pm 2.32$) | 5.81 ($\pm 2.50$) | | | |
| Nicotine | 29 | 5.73 ($\pm 2.13$) | | 29 | 4.95 ($\pm 1.43$) | 3.97 ($\pm 1.76$) |
| | 28 | 5.83 ($\pm 2.10$) | 5.78 ($\pm 2.33$) | | | |
| Placebo | 33 | 5.84 ($\pm 2.67$) | | 33 | 4.26 ($\pm 1.57$) | 3.46 ($\pm 2.07$) |
| | 32 | 5.51 ($\pm 2.53$) | 5.84 ($\pm 2.67$) | | | |

Five participants could not tolerate the fixed 46 °C stimulus during familiarisation, therefore a lower temperature of 45 °C was used for both pre- and post-gum PHP assessments for these participants, as reported in the protocol (see Methods). Two participants were excluded from mediation analysis assessing the effects of nicotine on PHP ratings: one from the placebo group due to an equipment issue during the second pain assessment, and one from the nicotine group that could not tolerate the baseline temperature for the second pain assessment (nicotine: n = 28; placebo: n = 32).

The difference score model for PHP (Figure 3; model fit: 𝜒^2^(24) = 25.874, 𝑝 = 0.36, CFI = 0.99, RMSEA = 0.036, SRMR = 0.078) suggests a significant total (𝑏 = −0.62, 𝑝 = .023, bootstrapped 95% CI: [−1.21, −0.15]) and direct (𝑏 = −0.68, 𝑝 = .021, bootstrapped 95% CI: [−1.28, −0.16]) effect of nicotine on change in PHP ratings (Figure 1). Change in pain was reduced by a factor of −0.68 for the nicotine group compared to the placebo group when controlling for confounding variables (Figure 3). See Supplementary Material 3.2, Table 5, for full model output.

Frequency spectra (plots A and B) and topographical plots (plot E) of PAF pre-post nicotine, are displayed by group in Figure 2, with PAF values displayed in Table 3. The difference score model suggests a significant effect of nicotine gum on change in global PAF (𝑏 = 0.085, p = .018, bootstrapped 95% CI: [0.017, 0.15]), suggesting change in PAF was increased by a factor of 0.085 for the nicotine group compared to placebo when controlling for confounding variables (Figure 3). Therefore, nicotine changed PAF and PHP ratings in the predicted directions compared to what would be expected without nicotine (i.e. placebo). However, as the estimated effect of change in PAF speed on change in PHP ratings was non-significant (𝑏 = 0.23, 𝑝 = .78, bootstrapped 95% CI: [-1.28, 1.9]), the change in PAF did not mediate the change in PHP ratings (i.e. indirect effect; 𝑏 = 0.019, 𝑝 = .81, bootstrapped 95% CI: [-0.093, 0.23]).

**Figure 2:**
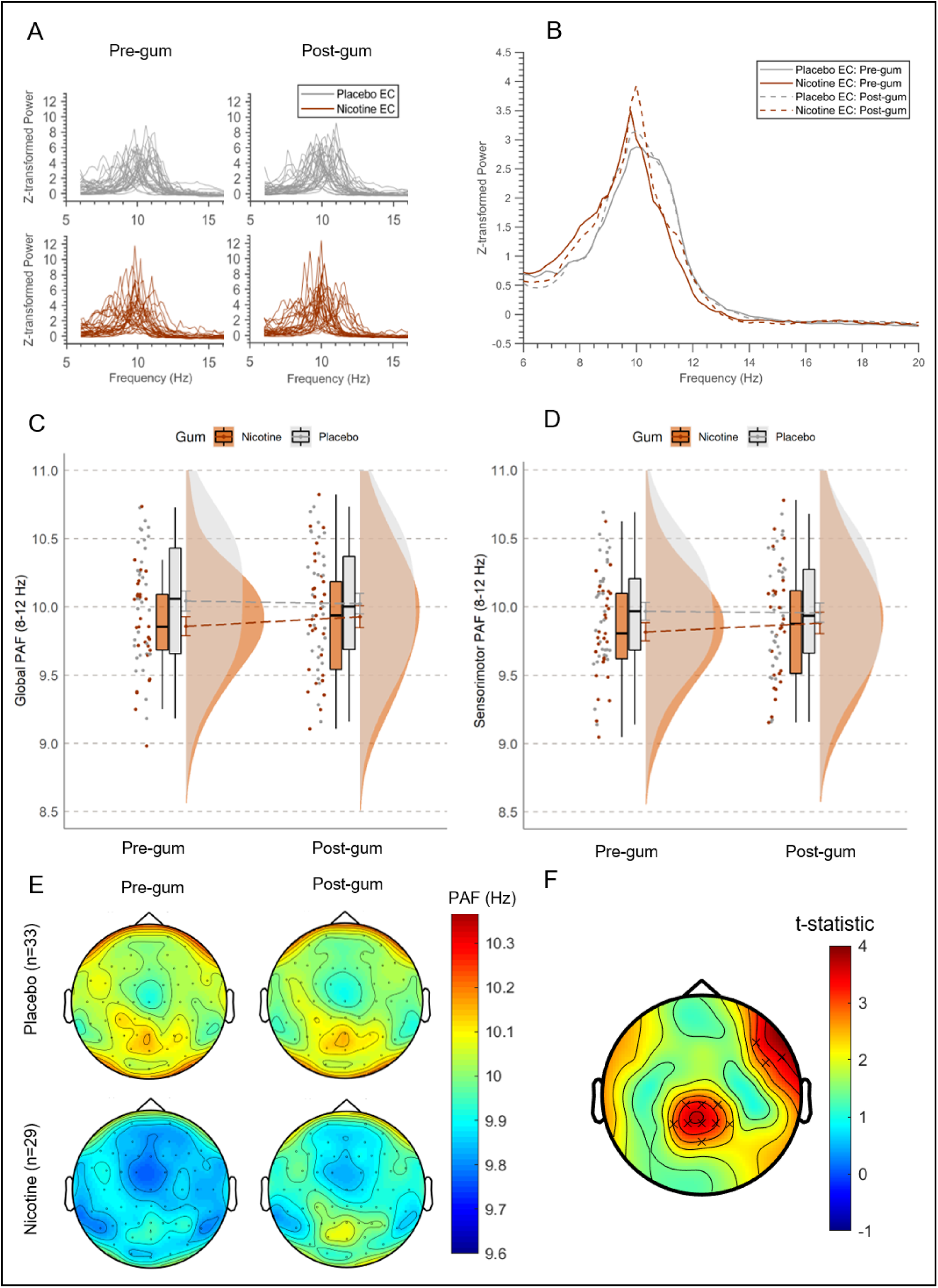
A) Individual spectral plots of eyes closed (EC) resting state EEG data showing dominant peaks within the alpha range of 8–12 Hz. Note that many participants’ peaks fall outside the narrow 9–11 Hz window. Top panel shows individual spectra for participants in the placebo group (in grey, n = 33), bottom panel shows individual spectra for participants in the nicotine group (in orange, n = 29). B) Spectral plots of eyes closed (EC) resting state EEG shows averaged spectra by group and time point. C) Raincloud plots of mean global and D) sensorimotor PAF values, calculated with a wide window (8–12 Hz) and the centre of gravity method, before and after chewing either nicotine (in orange) or placebo (in grey) gum. E) Topographical plots of PAF calculated for each electrode with the centre of gravity method and 8–12 Hz window are shown for the pre-gum baseline (rest 1) and post-gum (rest 3). F) Topographical plot of t-test statistics produced by cluster-based permutation analysis of nicotine group, displaying significant difference in PAF after chewing nicotine gum in two separate clusters marked by crosses where p<.01 for adjacent electrodes (x). These effects were most pronounced in the central-parietal and right-frontal electrodes.

**Figure 3:**
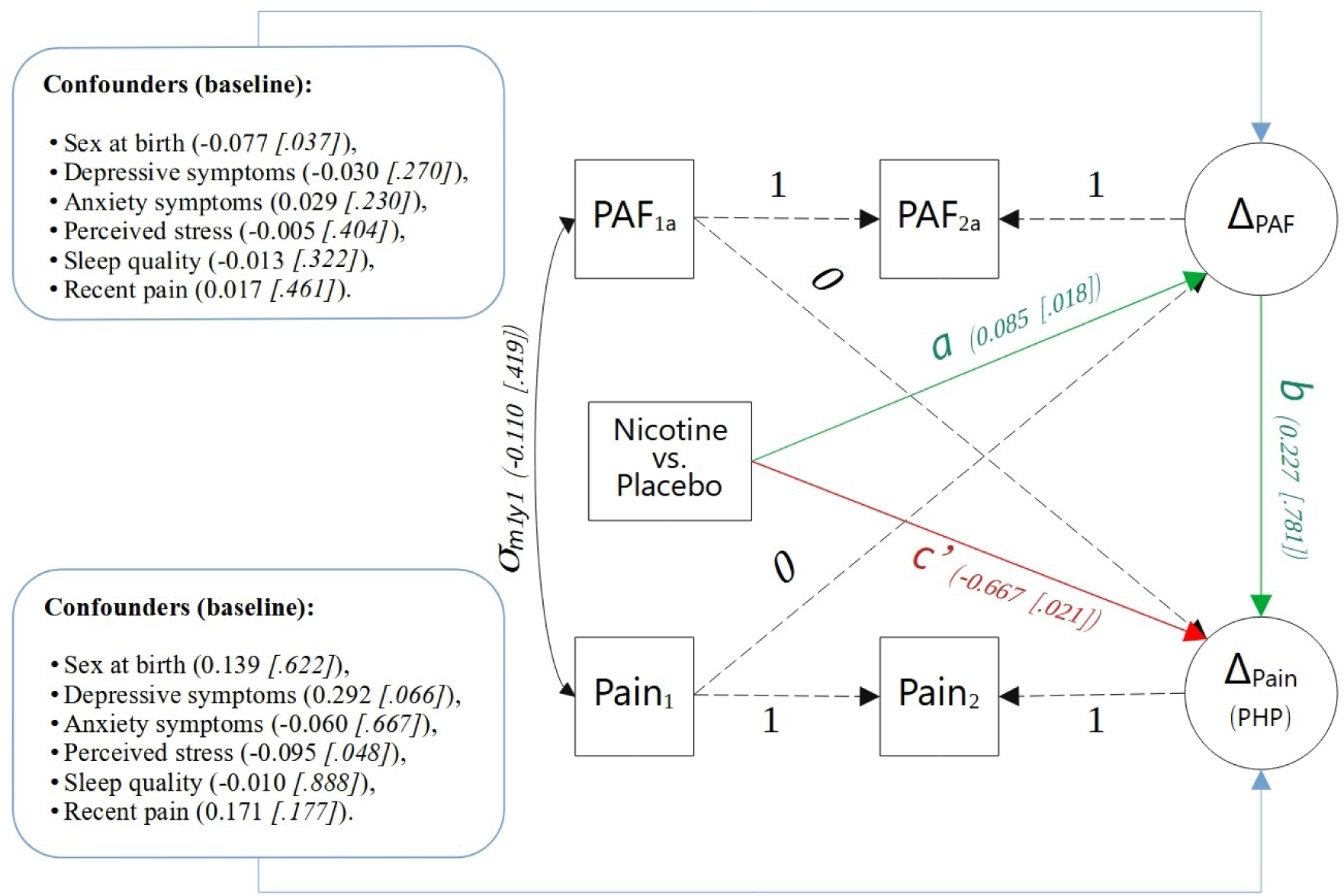
Path model displaying unstandardised estimates and p-values for the difference score two-wave latent change score (2W-LCS) model assessing the effect of nicotine against placebo (i.e. gum group) on change in global 8–12 Hz peak alpha frequency (i.e. Δ_𝑃𝐴𝜞_) and change in mean phasic heat pain (PHP) ratings (i.e. Δ_𝑃𝑎𝑖𝑛_). Green arrows highlight the indirect effects (i.e. a and b), the red arrow highlights the direct effect (i.e. c’), the blue lines highlight pre-determined confounding variables, and dashed lines denote fixed effects. This model suggests that chewing nicotine gum, compared to placebo, increases the speed of PAF, and decreases PHP ratings, however the change in PAF does not influence the change in PHP ratings, therefore changes in PHP ratings after nicotine gum are not mediated by changes in PAF.

**Table 3:** Mean ± standard deviations (SD) for sensorimotor and global peak alpha frequency (PAF) pre- and post-gum chewing, for the whole sample and by nicotine and placebo groups. PAF calculated for wide frequency window (8–12 Hz) with the centre of gravity method. Note that the 62 participants is the full sample that was used for mediation analysis of cuff pressure algometry (CPA) pain ratings and baseline correlations of PAF with CPA and phasic heat pain (PHP), whilst the 60 participants is the sample used for mediation analysis of PHP, as two participants were deleted list wise due to missing post-gum PHP assessments, one from each gum group.

| Gum group | Sensorimotor |  |  |  |  |  |
| --- | --- | --- | --- | --- | --- | --- |
|  | PAF |  |  | Global PAF |  |  |
|  | n | Pre-gum | Post-gum | n | Pre-gum | Post-gum |
| Total | 62 | 9.90 ($\pm 0.37$) | 9.92 ($\pm 0.41$) | 62 | 9.96 ($\pm 0.41$) | 9.98 ($\pm 0.43$) |
| | 60 | 9.90 ($\pm 0.38$) | 9.93 ($\pm 0.41$) | 60 | 9.96 ($\pm 0.41$) | 9.98 ($\pm 0.44$) |
| Nicotine | 29 | 9.82 ($\pm 0.36$) | 9.88 ($\pm 0.42$) | 29 | 9.86 ($\pm 0.37$) | 9.93 ($\pm 0.44$) |
| | 28 | 9.81 ($\pm 0.36$) | 9.87 ($\pm 0.43$) | 28 | 9.85 ($\pm 0.38$) | 9.92 ($\pm 0.44$) |
| Placebo | 33 | 9.97 ($\pm 0.38$) | 9.96 ($\pm 0.41$) | 33 | 10.04 ($\pm 0.42$) | 10.02 ($\pm 0.43$) |
| | 32 | 9.98 ($\pm 0.38$) | 9.97 ($\pm 0.40$) | 32 | 10.05 ($\pm 0.42$) | 10.04 ($\pm 0.43$) |

There was also a significant effect of perceived stress at baseline on change in PHP ratings when controlling for group allocation and other confounding variables (𝑏 = −0.096, 𝑝 = .048, bootstrapped 95% CI: [−0.19, −0.000047]), where higher perceived stress resulted in larger decreases in PHP ratings (see Supplementary Material 3.3 for post-hoc analysis of stress). There was also a significant effect of sex on change in PAF (𝑏 = −0.077, p = .037, bootstrapped 95% CI: [−0.16, −0.011]); change in PAF was reduced by a factor of −0.077 for males compared to females when controlling for gum group and confounders, therefore the increase in PAF speed was greater for females than males (see Supplementary Material 3.4 for post-hoc analysis of sex differences).

A complete table of PAF and power values for all four time points is displayed in Supplementary Material 4.1. Exploratory analysis of interactions between time point (i.e. pre-/post-gum) and group (i.e. nicotine/placebo) on PHP ratings, PAF speed, PHP ratings, and alpha power, using repeated measures analysis of variance (RM-ANOVA) that do not control for confounding, are displayed in Supplementary Materials 4 and 5. Notably, exploratory RM-ANOVAs still indicated interactions between gum group and time point on PAF (8–12 Hz) indicating that PAF only increased for the nicotine group after chewing (Supplementary Material 4), but there was no interaction between gum group and time point on PHP ratings (Supplementary Material 5). This underscores the necessity of controlling for confounding variables, as nicotine may only produce pain reduction for those with higher stress, and calls for further investigation into the robustness of the effect of nicotine on heat pain ratings.

#### Nicotine gum modulated global PAF (8–12 Hz), with no total, direct, or indirect effects on CPA ratings

The CPA model involved four minutes of continuous cuff pressure to the left calf muscle that was initially calibrated to each individual’s 4/10 pain (see Methods), and was most frequently described as mild/moderate cramping, aching, throbbing, and heavy pain (Figure 1, plot B). Raw time courses of pain ratings during CPA are displayed in Figure 1 (plot D), from which mean pain during the 4 minutes of stimulation was calculated (Table 2, Figure 1, plot F). No participants were excluded from the analyses of pressure pain (nicotine: n = 29; placebo: n = 33).

Estimates from the difference score 2W-LCS model for CPA pain ratings (Figure 4) for total (𝑏 = −0.28, 𝑝 = .50, bootstrapped 95% CI: [-1.13, 0.49]), direct (𝑏 = −0.17, 𝑝 = .69, bootstrapped 95% CI: [-1.07, 0.59]), and indirect (𝑏 = −0.11, 𝑝 = .53, boot-strapped 95% CI: [-0.61, 0.18]) effects of nicotine on CPA ratings were non-significant (model fit: 𝜒^2^(24) = 29.55, 𝑝 = 0.20, CFI = 0.97, RMSEA = 0.061, SRMR = 0.085). Therefore, there was no evidence to suggest that nicotine gum directly or indirectly influenced change in CPA ratings.

**Figure 4:**
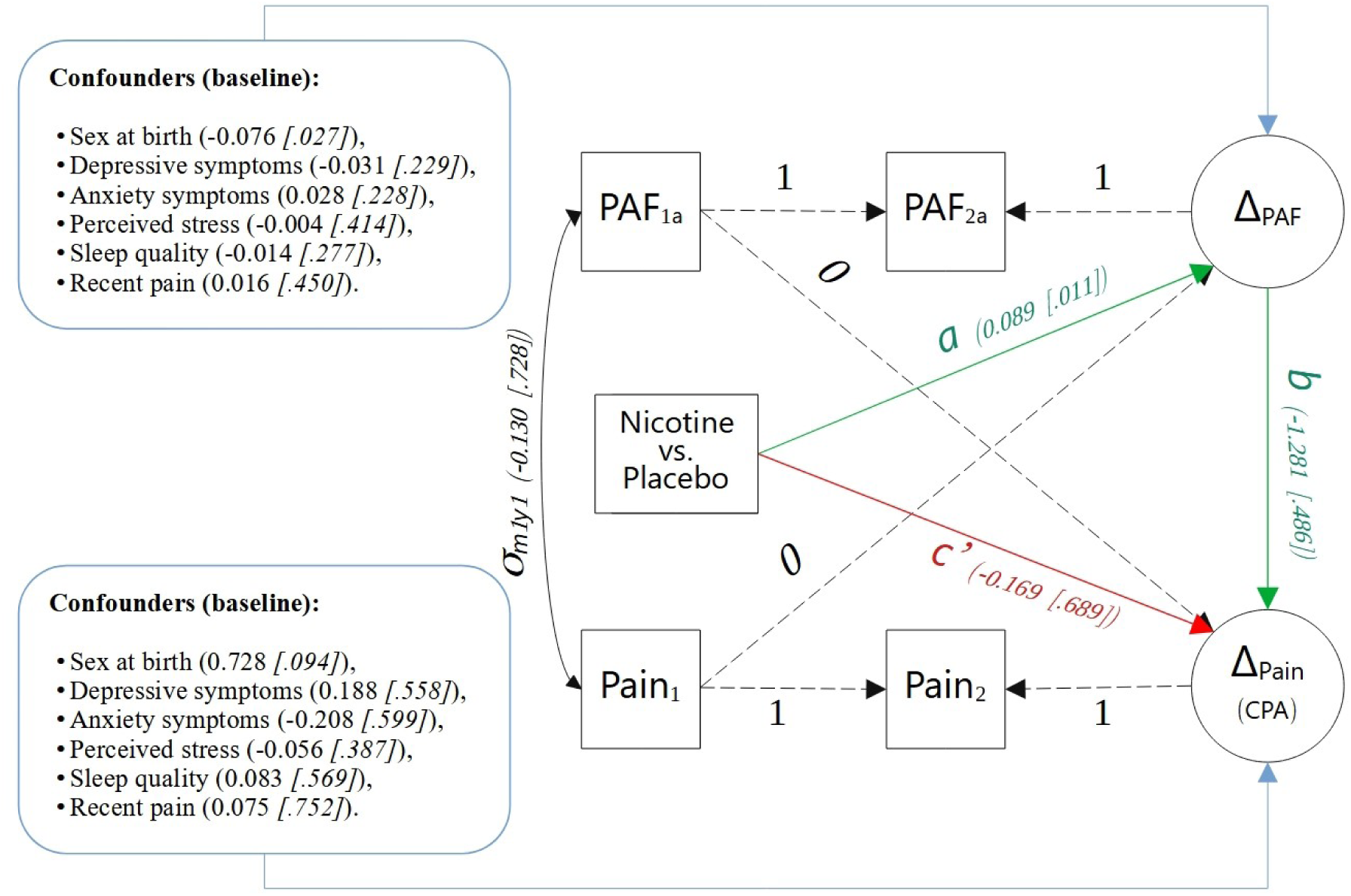
Path model displaying unstandardised estimates and p-values for the difference score two-wave latent change score (2W-LCS) model assessing the effect of nicotine against placebo (i.e. gum group) on change in global 8–12 Hz peak alpha frequency (i.e. Δ_𝑃𝐴𝜞_) and change in mean cuff pressure algometry (CPA) ratings (i.e. Δ_𝑃𝑎𝑖𝑛_). Green arrows highlight the indirect effects (i.e. a and b), the red arrow highlights the direct effect (i.e. c’), the blue lines highlight pre-determined confounding variables, and dashed lines denote fixed effects. This model suggests that chewing nicotine gum, compared to placebo, increases the speed of PAF and does not influence CPA ratings, directly or indirectly.

Consistent with the PHP model, there was a significant effect of nicotine gum on change in global PAF (8–12 Hz) speed (𝑏 = 0.089, p = .011, bootstrapped 95% CI: [0.018, 0.16]), where change in PAF speed increased by 0.089 for the nicotine group compared to the placebo group. Additionally, there was again an effect of sex on change in PAF (𝑏 = −0.076, p = .027, bootstrapped 95% CI: [−0.15, −0.01]), where the increase in PAF speed was greater for females than males. And the estimated effect of change in PAF speed on change in pain ratings was non-significant for CPA (𝑏 = −1.28, 𝑝 = .49, bootstrapped 95% CI: [-5.21, 2.11]).

Exploratory analysis of interactions between time point (i.e. pre-/post-gum) and group (i.e. nicotine/placebo) on CPA ratings using RM-ANOVA (Supplementary Material 5) suggested that CPA pain ratings decreased after chewing gum irrespective of group (Figure 1, plots D and F).

#### Exploratory data-driven analysis of alpha

The effect of nicotine on PAF speed was only significant for global PAF with a wide frequency window (8–12 Hz) and not for sensorimotor PAF (i.e. Cz, C1, C2, C3, C4) or narrow frequency windows (9–11 Hz; Supplementary Materials 3.4–3.6). This suggests that nicotine produces change in PAF somewhere in sensor-space, but not in the pre-defined sensorimotor region comprising central electrodes. An exploratory cluster-based permutation analysis (see Methods) showed significant increases in PAF speed after chewing nicotine gum, with this effect being maximal over two spatially separated clusters of electrodes: one cluster over the central-parietal region, and another over the right-frontal region (Figure 2, plot F). None of the electrodes exceeded the threshold of .05 to be clustered in the placebo group.

The potential mediating effect of this change in PAF on change in PHP and CPA was explored (not pre-registered) by averaging within each cluster (central-parietal: CP1, CP2, Cpz, P1, P2, P3, P4, Pz, POz; right-frontal: F8, FT8, FT10) and across both clusters. This averaging across electrodes produced three new variables, each assessed in relation to mediating effects on PHP and CPA ratings. The resulting in six exploratory mediation analysis (difference score 2W-LCS) models demonstrated minimal differences from the main analysis of global PAF (8-12 Hz), except for the expected stronger effect of nicotine on change in PAF (𝑏𝑠 = 0.11–0.14, ps < .003; Supplementary Materials 3.8–3.10).

Lastly, use of in-house code developed to automatically extract a sensorimotor alpha component using independent component analysis (ICA) showed similar results to the sensorimotor ROI mediation analysis (Supplementary Material 3.11). This supports the robustness of the PAF—pain effect regardless of the method used to isolate sensorimotor alpha.

#### Summary of mediation analyses

In summary, nicotine demonstrated significant total and direct effects on pain intensity ratings (i.e., reduction compared to placebo) during PHP, but not during CPA, where pain ratings decreased irrespective of gum group. There was also a significant effect of nicotine on change in global PAF speed (i.e., increased compared to placebo). However, there was no evidence of an indirect effect, suggesting that the change in PAF speed did not mediate the change in pain intensity ratings during PHP.

### 2.3 Relationship between PAF and pain at baseline

Evidence for negative correlations between pre-gum PAF (8–12 Hz) and pre-gum pain ratings during PHP (𝜏 = −0.14, BF_10_ = 0.52) and during CPA (𝜏 = −0.086, BF_10_ = 0.33; Figure 5) is inconclusive. Therefore, there was no evidence for or against slower PAF being associated with higher pain ratings. However, in an exploratory analysis (not pre-registered) of separate correlations in males and females (Figure 5, plot C) akin to those conduced in previous research on this topic [10], there was moderate evidence for a negative relationship between global PAF (8–12 Hz) and PHP ratings in males (𝜏 = −0.39, BF_10_ = 3.22). In contrast, the evidence for a relationship between global PAF and PHP ratings in females was equivocal (𝜏 = −0.032, BF_10_

**Figure 5:**
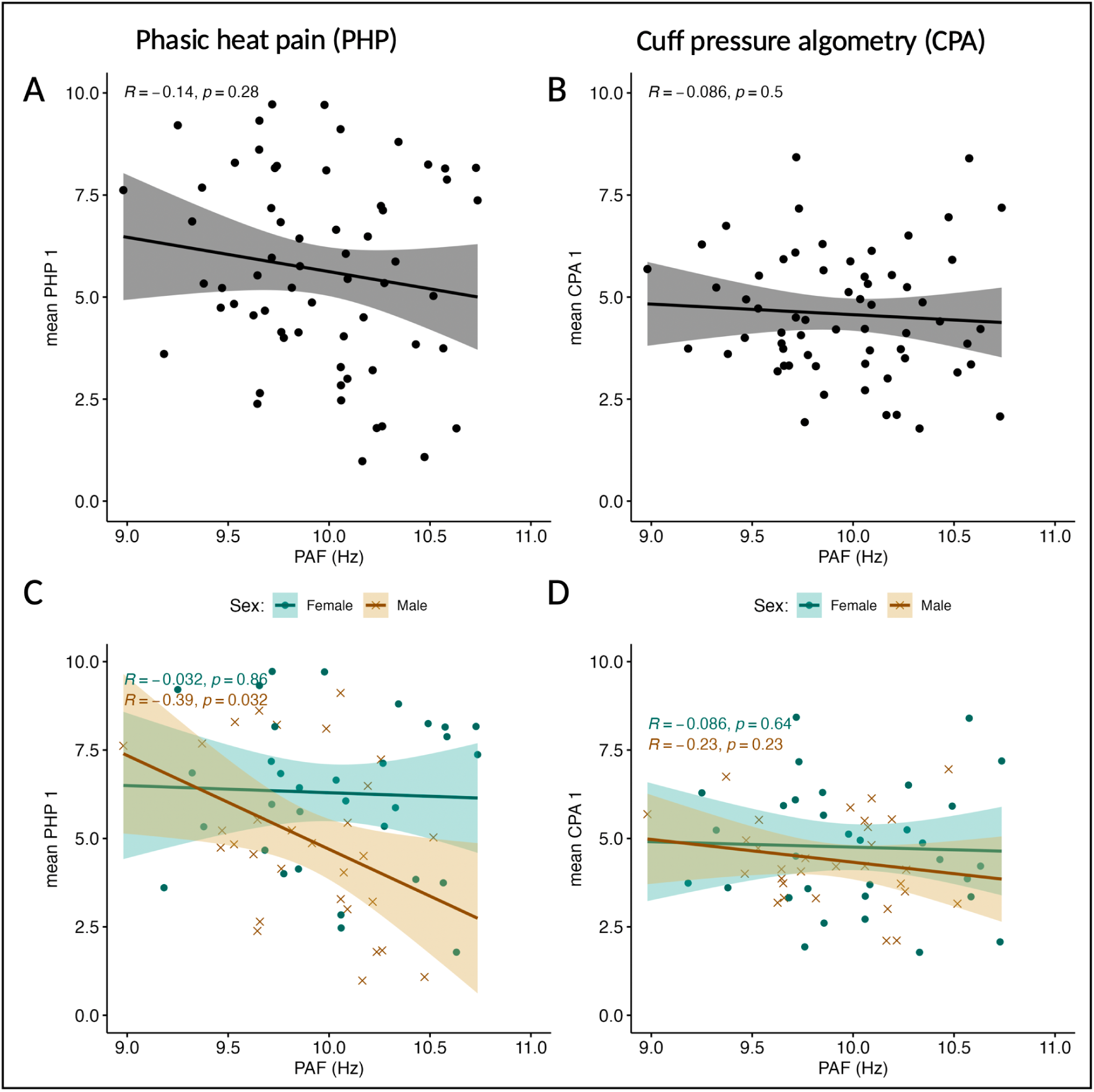
Wide band (8–12 Hz) global peak alpha frequency (PAF) does not correlate with A) mean pain ratings during phasic heat pain (PHP) or B) cuff pressure algometry (CPA). Regression lines and shaded 95% confidence intervals. Correlations separated by sex for females in teal (n = 32) and males in brown (n = 30) for C) PHP and D) CPA, suggest a strengthening of a negative relationship between sensorimotor PAF and PHP ratings for males (BF_10_ = 3.22).

= 0.40). Separating by sex for the correlations between PAF and CPA ratings still produced inconclusive evidence for a relationship (males: 𝜏 = −0.23, BF_10_ = 0.61; females: 𝜏 = −0.086, BF_10_ = 0.40).

The relationship between PAF and pain was also assessed using median splits on PAF, showing similar patterns of slower PAF being associated with higher PHP ratings in males only (Supplementary Material 6.1). Correlations between global PAF (8–12 Hz) and max pain during PHP also suggested slower PAF was only associated with higher max PHP ratings in male participants (Supplementary Material 6.2). Five participants could not tolerate the 46 °C stimulus during familiarisation, therefore a lower temperature of 45 °C was used for these participants. Repeat analyses excluding these five participants did not impact the observed relationships (Supplementary Material 6.3).

In summary, the evidence for a relationship between PAF and pain is inconclusive, with the exception of the relationship between PAF and PHP ratings by sex, where there was moderate evidence of a negative correlation in male participants.

## 3 Discussion

In this pre-registered study, we probed the proposed PAF–pain relationship [8–12] by administration of nicotine – a substance suggested to acutely alter PAF [46]. Compared to placebo, our data demonstrate three key findings: 1) nicotine increased PAF speed; 2) nicotine reduced prolonged heat pain intensity but only when controlling for confounding and not for prolonged pressure pain model; and 3) decreases in prolonged heat pain intensity were not mediated by changes in PAF. This suggests that changes in heat pain due to nicotine occur through mechanisms unrelated to change in PAF and require replication as the effects required control for confounding and the reduction was small in magnitude. In addition, when assessing the whole sample, we found inconclusive evidence for 4) slower PAF speed at baseline being associated with higher pain ratings for both the heat and pressure pain models. However, during exploratory analysis we found moderate evidence that slower PAF speed at baseline is associated with higher heat pain ratings at baseline in males but not females. In contrast PAF was not associated with pressure pain ratings for males or females.

### 3.1 The effect of nicotine on PAF

We aimed to determine whether nicotine gum could increase PAF speed. In line with previous literature in non-smokers [38,47,48], we contribute additional evidence that chewing 4 mg of nicotine gum produces a statistically significant increase in PAF speed in non-smoking nicotine gum naïve, pain-free, adult participants. An exploratory cluster-based permutation analysis indicated that the change in wide window PAF (8–12 Hz) produced by nicotine, was most pronounced in a cluster of electrodes in the central-parietal region and a separate cluster of electrodes in the right-frontal region, rather than in the pre-registered sensorimotor ROI (i.e., Cz, C1, C2, C3, C4).

Research on nicotine-induced changes in PAF in healthy non-smokers has largely relied on small samples (<20) using injections or patches [47,48], with Bowers and colleagues [38] providing the only larger study (n = 62 males) using 6 mg nicotine gum. Their results showed modest PAF increases (∼0.34–0.51 Hz; 3.5–5.8%) compared to the larger effects (1–2 Hz; ∼10–20%) reported in earlier small-sample studies, with magnitude and scalp distribution influenced by COMT genotype: Met allele carriers exhibited nicotine-induced increases (Val/Met: frontal, central, occipital; Met/Met: parietal), while Val/Val homozygotes showed no change. The present study −— now the largest to include both male and female non-smokers —-found smaller effects, likely due to the lower nicotine dose (4 mg), consistent with dose-dependent PAF modulation reported in abstinent smokers [69]. Broader EEG coverage (64 vs. 8 electrodes), sex inclusion, and unmeasured genetic variation may also account for differences between the present study and Bowers and colleagues [38]. Interestingly, beyond nicotine’s main effect on PAF, our 2W-LCS mediation model indicated a significant influence of sex, with females showing greater PAF increases than males. Given that female participants were significantly lighter than males (Supplementary Material 3.4) and the model did not test the interaction between sex at birth and gum group, these findings should be interpreted cautiously and warrant further investigation.

### 3.2 The effect of nicotine on pain

The choice of nicotine was driven by its potential to change PAF, but also, short-term nicotine use is thought to have acute analgesic properties in experimental settings, with a review reporting that nicotine increased pain thresholds and pain tolerance [51]. In addition, research in a rat model suggests analgesic effects on mechanical thresholds after short-term nicotine use [70]. However, previous research has not assessed the effects of nicotine on prolonged experimental pain models. The present study found that 4 mg of nicotine reduced heat pain ratings during prolonged heat pain compared to placebo for our human participants, but that prolonged pressure pain decreased irrespective of which gum was chewed. Our findings are thus partly consistent with the idea that nicotine may have acute analgesic properties [51], although further research is required to explore factors that may influence nicotine’s potential impact on a variety of prolonged pain models. We further advance the literature by reporting this effect in a model of prolonged heat pain, which better approximates the experience of clinical pain than short lasting models used to assess thresholds and tolerance [51].

However, we note that the observed effect of nicotine on pain was small in magnitude, and most prominent in comparison to the effect of placebo, where pain ratings increased after chewing, which brings into question whether this reduction in pain is meaningful in practice. While acknowledging the modest effect size, it’s essential to consider the context of our study’s focus. Assessing the clinical relevance of pain reduction is pertinent in applications involving the use of any intervention for pain management [71]. However, from a mechanistic standpoint, particularly in understanding the implications of and relation to PAF, the specific magnitude of the pain effect becomes less pivotal, as small effects can still have mechanistic or biological meaning without establishing clinical meaning [72]. Nevertheless, future research should examine whether effects on pain increase in magnitude with different nicotine administration regimens (i.e. dose and frequency).

We did not see an analgesic effect for prolonged pressure pain, as ratings during pressure pain were lower at the second assessment irrespective of gum allocation. The way we calibrated the pressure pain model may have impacted our results. Previous literature has used cuff inflation where the cuff pressure increases until the participant reports 4/10 pain, at which point it remains stable for the desired number of minutes [59–61,73]. In contrast, a separate calibration was conducted at the beginning of this experiment to calculate the pressure used for the remainder of the study, based on the average of three trials of increasing pressure to 4/10 pain (see Methods). This method may result in lower pain ratings at the second pain induction with the same pressure in a same-session repeated measures investigation.

The mechanisms by which nicotine may influence pain likely depend on the type of pain and whether nicotine use is acute or chronic, along with several other contributing factors [51]. For example, in addition to the effect of nicotine on prolonged heat pain ratings, our results suggest an effect of stress on changes in heat pain ratings, with those self-reporting higher stress at baseline having greater reductions in pain. Our post-hoc analysis suggested that this relationship between higher stress and larger decrease in PHP ratings was only present for the nicotine group (Supplementary Material 3.3). As stress is linked to nicotine use [74,75] and pain [76–78], these interactions should be explored in future.

### 3.3 The relationship between PAF and pain

Prior research suggests that slower PAF is associated with higher pain sensitivity [8–11], and while the validation of PAF as a predictive biomarker continues [13,14], the question of whether PAF is mechanistically linked to pain remains unanswered. Modulating PAF speed and pain sensitivity with nicotine is necessary but not sufficient to demonstrate a causal pathway from nicotine to PAF to pain (i.e. intervention → mediator → outcome) [79]. We did not find evidence for a causal relationship between PAF and pain sensitivity, as our 2W-LCS model did not show a significant mediating (i.e. indirect) effect of change in PAF on change in pain due to nicotine. The implication of this is that pain sensitivity may not be dependent on state changes in PAF, thus changing PAF may not be a useful target for pain modulation. However, as this is the first study attempting to directly manipulate PAF, observing only small effects, we cannot confidently state that a causal relationship between PAF and pain sensitivity does not exist, and future research should explore these relationships with larger sample sizes, larger doses of nicotine, and alternative PAF modulators.

As an alternative explanation, an indirect effect may exist, but could not be seen in the present study due to the single, low dose of nicotine or for other methodological reasons. For example, the relationship between change in PAF and change in pain may be non-linear, and therefore difficult to see using a 2W-LCS model that is based on linear regression analyses. Additionally, the use of global PAF may have introduced mediation measurement error into our mediation analysis. The spatial precision used in the current study was based on previous literature on PAF as a biomarker of pain sensitivity, which have used global and/or sensorimotor ROIs [8,10]. Identification and use of the exploratory electrode clusters found in this study could build upon the current work (e.g., [80]). However, exploratory analysis of the clusters found in the present analysis demonstrated no influence on mediation analysis results (Supplementary Materials 3.8–3.10). Alternatively, ICA could be used to identify separate sources of alpha oscillations [81], as used in other experimental PAF–pain studies [8,18], which could aid to disentangle the potential relevance of different alpha sources in the PAF–pain relationship. Although this comes with the need to develop more reproducible and automated methods for identifying such components.

In-house code was developed to compare a sensorimotor component to the results presented in this manuscript (Supplementary Material 3.11), showing similar results to the sensorimotor ROI mediation analysis presented here. However, examination of which alpha – be it sensor or source space – are related to pain, how they can be robustly represented, and how they can be manipulated are ripe avenues for future study.

Furthermore, there may have been other mediators that suppress the mediating effects of PAF within our model [66,82,83]. This is plausible, as nicotine is thought to act on pain through multiple pathways [52,54,84], and more direct PAF modulation could more definitively evaluate whether PAF is mechanistically linked to pain sensitivity. Future interventions to modulate PAF could use non-invasive brain stimulation (NIBS), such as transcranial alternating current stimulation (tACS) or repetitive transcranial magnetic stimulation (rTMS), to influence neuronal firing and thereby directly target the brain mechanisms theorised to generate PAF [85–87]. This approach has since been investigated following the initial preprint of the present manuscript [88,89]. In addition, other approaches to modulating approaches could be considered, such as exercise [32–34], visual stimulation [35,36], cannabis [90,91], and neurofeedback [92,93], while considering their viability in terms of blinding, tolerability, and scalability.

Regarding the relationship between PAF and pain at baseline, the negative correlation between PAF and pain seen in previous work [7–11,16] was only observed here for male participants during the PHP model for global PAF in an exploratory analysis. Effect sizes for the relationship between PAF and pain ratings from previous work were in the range of r = 0.4–0.6 [8–11]. In the present study we saw a weaker relationship for PHP pain (N = 62, r = −0.14). However, sex-stratified analysis demonstrated that males exhibit comparable effect sizes to previous studies (n = 30, r = −0.39), while females did not show a relationship between PAF and heat pain (n = 32, r = −0.032). Subtle variations in experimental design and stimulus delivery could have influenced the PAF–pain relationship. Crucially, the present study did not calibrate the temperature of the PHP model based on participants’ thresholds for moderate pain. Consequently, the fixed temperature did not produce a moderately painful stimulus for all participants, in contrast to previous studies that calibrated to a moderately painful stimuli [10]. This discrepancy may account for the correlation observed between PAF and heat pain ratings among only the male participants in our study, as fixed stimuli can create ceiling or floor effects [94]. Previous literature has also acknowledged the potential for sex differences in the PAF–pain relationship [17].

In parallel, we saw no indication of a relationship between PAF and pain ratings during CPA. The introduction of the CPA model, specifically calibrated to a moderate pain threshold, provides further support for the notion that the relationship between PAF and pain could be specific to certain pain types [17,28] or possibly tissues targeted. Prolonged heat pain was pre-dominantly described as moderate/severe shooting, sharp, and hot pain, whereas prolonged pressure pain was predominantly described as mild/moderate throbbing, cramping, and aching in the present study. It is possible that the PAF–pain relationship is specific to particular pain models and protocols [12,17].

Previous experimental pain literature has suggested that slower PAF is related to higher pain sensitivity for PHP [10], capsaicin heat pain [8,10], and nerve growth factor (NGF) [9,13] models, while faster PAF has been related to higher pain sensitivity in tonic heat pain [7] and NGF [15]. Here we contribute to this literature by showing no relationship between baseline PAF and pain sensitivity during the CPA model. Further investigation of the PAF–pain relationship with a variety of prolonged pain models is warranted [12].

### 3.4 Study limitations and constraints

This study presents opportunities for future research due to certain limitations. First, the dose of nicotine administered was equivalent to one cigarette [95]; however, to enhance the efficacy of this research, exploring higher doses or dosing based on weight could be beneficial. Second, blood or saliva samples were not collected in the present study, therefore it is unclear whether participants absorbed similar amounts of nicotine. Future studies in this area should also be powered to perform sex-disaggregated analyses, as our findings indicate a relationship between baseline PAF and subsequent heat pain intensity only in males, and stronger nicotine modulation of PAF in female participants. The ability to detect changes in PAF could be considerably impacted by the frequency resolution used during Fourier Transformations, an element that is overlooked in recent methodological studies on PAF calculation [16,96]. Changes in PAF within individuals might be obscured or conflated by lower frequency resolutions, which should be considered further in future research.

### 3.5 Conclusion

In this study, we employed a rigorous double-blind, randomised, placebo-controlled design, pre-registered the analysis, and introduced a novel difference score 2W-LCS mediation model to examine the mediating role of changes in PAF on pain while controlling for pre-determined confounders. These methodological choices allowed for a comprehensive investigation into whether PAF plays a mechanistic role in pain sensitivity.

We have shown that, compared to placebo, nicotine gum produced statistically significant increases global PAF speed and decreases pain ratings during prolonged heat pain. However, the effects of nicotine on heat pain were not mediated by changes in PAF. Therefore, we find no evidence that PAF is causally related to pain. The implication of this is that changing PAF may not be a useful target for pain modulation, because pain sensitivity may not be dependent on state changes in PAF. However, this is the first study to test modulation of PAF as a mechanism for reducing pain intensity. Moreover, while effects on PAF appear small but robust, effects on heat pain required control for confounding to be observed. More randomised experiments on the effects of nicotine that use larger sample sizes, higher nicotine doses, and pharmacokinetic tracking, whilst also reducing mediator measurement error are needed to confirm the present findings and uncover whether these factors account for the lack of mediation effect seen here.

A growing body of evidence indicates a relationship between PAF and pain, including chronic pain studies [17,24,26–28,97], and experimental pain research from our own [8–12,14] and other groups [7,15–18]. Alongside validation of its clinical predictive utility [14,96], further research is needed to ascertain the causal nature of the relationship, explore PAF as a potential target for pain modulation, and gain deeper mechanistic insights into the PAF–pain relationship.

## 4 Methods

### 4.1 Design overview

The study employed a randomised, double-blind, placebo-controlled, parallel experimental design. Testing sessions were balanced for treatment (i.e. nicotine or placebo gum) and stratified by sex. Individuals were required to attend one 150 minute session. The experimental flow for this study is summarised in Figure 6.

**Figure 6:**
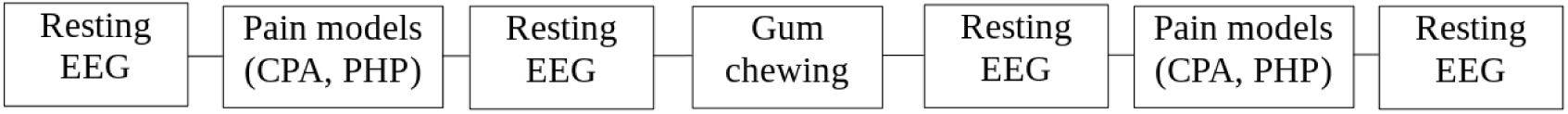
Timeline of experimental events. Electroencephalography (EEG) data were collected during eyes open and eyes closed resting states before and after each pain assessment. Prolonged pain assessments consisted of cuff pressure algometry (CPA) followed by phasic heat pain (PHP) models, and took place before and after gum chewing.

### 4.2 Participants

From June 2021 to March 2022, 62 individuals participated at Neuroscience Research Australia (NeuRA). Inclusion criteria for the study were for participants to be healthy, pain free, non-smoking, and between 18–45 years of age, inclusive. The sample size was informed by resource and feasibility constraints [98] and aligns with or exceeds that of comparable studies in the field [38], covering 60 participants in total, with two extra in case of incomplete data identified during later data analysis.

Non-smokers were defined as individuals who have smoked fewer than 100 cigarettes and/or e-cigarettes in their lifetime and have not smoked over the past 12 months. Exclusion criteria: previous use of nicotine gum; history of neurological, musculoskeletal, cardiovascular, or psychiatric conditions; other major diseases or disorders; current muscle pain; breastfeeding or pregnant; use of opioids, illicit drugs, or excessive alcohol in the past three months; or unable to provide informed consent.

### 4.3 Ethics approval and consent to participate

The Human Research Ethics Committee (HREC) Executive at the University of New South Wales (UNSW) approved this project (HC190466).

### 4.4 Experimental protocol

Participants were screened for eligibility by telephone and were asked to refrain from drinking coffee or caffeinated drinks two hours prior to the start of testing, as caffeine may impact nicotine absorption [99,100].

After signing informed consent, cool and warmth detection tests were conducted, followed by PHP familiarisation, then CPA calibration. During these introductions the participants were also familiarised with the VAS pain rating.

EEG capture started with the control condition of three cycles of warm temperatures, after which the remaining experimental flow is illustrated in Figure 6. The first resting state period was recorded, followed by the first pain assessments (i.e. CPA and PHP model), which were immediately followed by the second resting state. EEG recorded during the pain assessments was not analysed for the present article. Participants were randomly assigned to one of only two conditions, thus after the second resting state EEG, either nicotine or placebo gum was administered with the chew procedure. Following completion of the chew procedure, the third resting state, the second pain assessments, and the fourth resting state were conducted. The experiment ended with participants completing the gum chew experience questionnaire and debrief.

### 4.5 Data collection procedures

#### Questionnaires

Participants completed a set of questionnaires to assess demographics, medical history, general health, mental health, pain, and sleep. This data was collected electronically and stored using REDcap [101,102].

Questionnaires included the following National Institutes of Health common data elements (CDE) for pain biomarkers: Pain Catastrophizing Scale (PCS) [103]; Brief Pain Inventory (BPI), Pain Severity and 7-item Interference subscales [104,105]; SF-8 to assess general health [106]; Patient Health Questionnaire-2 (PHQ-2) to assess depression [107]; Generalised Anxiety Disorder Questionnaire-2 (GAD-2) to assess anxiety [108]; Pennebaker Inventory of Limbic Languidness (PILL) to assess reporting of physical symptoms and sensations [109]; Tobacco, Alcohol, Prescription medications, and other Substances (TAPS); sleep scale. The sleep scale asked “how would you rate your sleep last night?”, on a scale from 1 (i.e. “poor sleep quality”) to 10 (i.e. “excellent sleep quality”). In addition, a health history form, the Karolinska sleepiness scale (KSS) [110], Epworth sleepiness scale (ESS) [111], the perceived stress scale (PSS) [112], and questions regarding participants current level of weekly physical activity were included.

#### Electroencephalography

EEG was recorded from 63 active wet electrodes (actiCap, Brain Products GmbH, Germany), embedded in an elastic cap (EASYCAP, GmbH) in accordance with the international 10-10 system. Recordings were referenced online to ‘FCz’ and the ground electrode was placed on ‘FPz’. A sampling rate of 5000 Hz was used and impedance was kept below 25 kOhms throughout testing using electrolyte gel. EEG signal was amplified and digitised using an actiCHamp Plus DC amplifier linked to BrainVision Recorder software (version 1.22.0101, Brain Products GmbH, Gilching, Germany).

For each recording, resting state EEG was obtained during eyes open (three minutes) followed by eyes closed (three minutes). The eyes open recording took place in low lighting while participants rested their gaze on a white fixation cross with a black background displayed on a monitor. The eyes closed recording took place with the lights off. For both recordings participants were asked to rest in a comfortable seated position and try not to fall asleep.

#### Pain assessment

Two tonic (i.e. prolonged) pain assessments were conducted during the study, a phasic heat pain (PHP) model [10] and a cuff pressure algometry (CPA) model [59,113]. Pain intensity was sampled at 1 Hz during pain stimulation using a visual analogue scale (VAS), with anchors ‘no pain’ on the left to ‘worst pain imaginable’ on the right, custom MATLAB (version R2021a) code (source available: https://osf.io/pc4rq/). Participants were instructed to press the ‘STOP’ button on the screen if they became too uncomfortable and wanted to stop the pain assessments.

The Short Form McGill Pain Questionnaire 2 (SF-MPQ-2) [114] was used after each pain model to assess experienced qualities (e.g., throbbing, cramping, burning).

#### Heat pain

*Equipment.* Thermal stimuli were applied to the volar surface of the left forearm using a thermal-contact heat stimulator (27 mm diameter Medoc Pathways CHEPS Peltier device, Medoc Advanced Medical Systems Ltd., Ramat Yishai, Israel).

*Cool and warmth detection thresholds.* The thermode was placed on the volar surface of the left arm at a neutral temperature (32 °C). Participants were instructed to press a button held in their right hand when they sensed a change in temperature. Four areas were marked out by drawing three perpendicular lines equidistant from the elbow crease to 4 cm below the base of the palm. On each trial, the researcher moved through these locations on the forearm in a randomly generated sequence. There were three trials each for warmth (ascending temperatures) and cool (descending temperatures) detection thresholds. Participants were required to consistently detect the warmth and cool transitions in order to continue with the experiment.

*Heat pain threshold.* Participants were required to press the hand-held button with their right hand the moment they felt any pain as the thermode temperature increased at a rate of 1.5 °C/second. This pain threshold test was conducted three times and an average heat pain threshold was calculated.

*Familiarisation.* Participants were initially introduced to the PHP model (see below) by receiving two cycles of the heat pain, to ensure they would tolerate the maximum temperature of 46 °C during the experiment and to familiarise participants with using the VAS. If a participant was uncomfortable with receiving 46 °C, the temperature was decreased by 1 °C and the PHP familiarisation was repeated to ensure this temperature was tolerable.

*Phasic warmth.* Participants experienced three rounds of a phasic warm stimulus. Stimulus trains consisted of a rise phase of 2 °C/s, followed by a 40 second phase where a temperature was held at 36 °C, then a fall phase at a rate of 1 °C/s to reach 32 °C. This neutral temperature was then held for 20 seconds before the cycle repeated three times.

*Phasic heat pain (PHP) model.* The PHP model consisted of a series of five repetitions of phasic stimuli, each lasting one minute 21 seconds. Stimulus trains consisted of a rise phase of 2 °C/s, followed by a 40 second phase where a temperature was held at 46 °C, then a fall phase at a rate of 1 °C/s to reach 32 °C. This neutral temperature was then held for 20 seconds before the cycle repeated five times. The PHP model lasted six minutes 45 seconds, and participants rated their pain continuously on the VAS. Similar methodology using repetitive-phasic models have shown that this procedure reliably evokes tonic pain and sensitisation in healthy participants [8,10].

#### Pressure pain

*Equipment.* Pressure was applied using a standard double-chamber adult arm cuff (length 45cm, width 15cm) placed in the middle of the left lower leg over the head of the gastrocnemius soleus muscle.

*Familiarisation and calibration.* Participants underwent a pressure cuff calibration. Similar to previous literature [60,62], the pressure was gradually increased manually (30 mmHg/5 seconds) until the participant reported a moderate pain intensity of 4/10 on the VAS and pressure was automatically recorded. This gradual pressure increase was conducted three times with a two minute break between each test, to establish the average pressure required to evoke 4/10 pain for each participant. The maximum pressure limit for this study was 100 kPa (750 mmHg) [63]. This is well within safe limits, as previous studies have used maximums of 180 or 200 kPa without adverse effects [59–61].

*Cuff pressure algometry (CPA) model.* Pressure was increased manually (30 mmHg/5 seconds) until the individualised pressure was reached, then held constant for four minutes. Participants rated their pain continuously using the VAS.

#### Gum chewing

Nicotine was administered orally in the form of 4 mg of cinnamon flavoured Nicorette® gum (Johnson & Johnson Inc., Markham, Ontario, Canada); comparable to smoking a single cigarette, this produces a blood concentration of approximately 15-30 mg/ml [95]. The placebo gum was cinnamon flavoured Xyloburst® natural chewing gum, which, other than containing no nicotine, is of a similar flavour, size, and texture to the nicotine gum. At the end of the experiment, before being debriefed, participants completed a short questionnaire about the gum chewing experience to test perceptual matching between the nicotine and placebo gums.

To promote nicotine absorption, participants rinsed their mouth with a glass of water for 10 seconds prior to receiving the gum, as performed previously [99]. We used a protocol similar to Bowers and colleagues [38], following manufacturer’s guidelines of a total chewing time of 25 minutes. Audio cues to bite the gum occurred every 30 seconds for a total of 25 minutes; participants were asked to alternate between the left and right sides of the mouth, and to park the gum between the teeth and cheek between chews. Participants watched a wildlife documentary during the chewing to maintain attention; all participants watched the same documentary to control for its possible effects.

#### Randomisation and blinding

Participants were randomised in a 1:1 ratio to receive either nicotine gum or placebo gum. The allocation list was created a priori, independently of the primary experimenter, using blocked randomisation based on sex at birth. The allocation was blind to both the participant and experimenter. All participants were told that they were receiving nicotine gum to provide equal expectations. Participants were debriefed at the end of the experiment and asked whether they thought they had been given nicotine, placebo, or if they weren’t sure. Participants were unblinded by the independent researcher separately. The primary experimenter remained blinded with dummy variables until completion of analysis.

### 4.6 Data processing

#### EEG pre-processing

Blinded preprocessing of de-identified EEG was conducted with custom code in MATLAB 2020a (v.9.8.0.1417392, The MathWorks, Inc., Natick, MA, USA) using EEGLAB 2020.0 [115] and FieldTrip (v.20200607) [116].

Using EEGLAB, for four resting states, each participant’s data were downsampled to 500 Hz, filtered between 2 and 100 Hz using a linear finite impulse response (FIR) bandpass filter, and the three minute periods of eyes closed resting state were visually inspected and overtly noisy channels with ≥ 8 signal deviations (e.g., popping, baseline drift, line noise) were removed. The average number of channels removed were 0.94, 0.98, 0.74, and 0.87 (range: 0–4) for the four resting states, respectively. Data from the remaining channels were re-referenced to the common average reference. Removed channels were not interpolated for the pre-registered global and sensorimotor ROI averaged analyses, but were interpolated for an exploratory cluster based permutation analysis using the nearest neighbour average method in Fieldtrip.

Using FieldTrip, data were segmented into 5 second epochs, with no overlap, visually inspected using *ft_databrowser()*, and epochs containing any electrode, muscle, or motion artefacts were removed. From a maximum of 36 epochs (i.e. 3 mins), the average number of remaining epochs was 31.21, 31.33, 32.13, and 31.39 (range: 24–36) for each resting state, respectively. No resting state contained less than two minutes of data [96]. Principal component analysis using the *runica* method was applied to identify and remove components reflecting eye blinks and/or saccades. The average number of components removed per resting state was 1.05, 1.2, 1.1, 1.2 (range: 0–2), respectively.

#### Quantification of peak alpha frequency (PAF)

Each participant’s PAF was estimated for the four eyes closed resting states. FieldTrip was used to calculate the power spectral density in the range of 2–40 Hz for each epoch in 0.2 Hz bins. A Hanning taper was applied to the data prior to calculating spectra to reduce any edge artefacts [117]. The PAF for each power spectral density was estimated using the centre of gravity (CoG) method [118–122]:

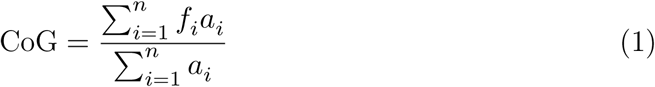

where 𝑓_𝑖_ is the 𝑖^th^ frequency bin, 𝑛 is the number of frequency bins within the frequency window (8–12 Hz or 9–1 Hz), and 𝑎_𝑖_ is the amplitude of the frequency for 𝑓_𝑖_. This equation states that PAF is the weighted sum of spectral estimates [118]. Global (i.e. average of all electrodes) and a sensorimotor region of interest were used (i.e. average of C3, C1, Cz, C2, and C4). PAF estimates for each participant at each time point were calculated. Out of 248 resting states recorded, 14 resting states had 4 ROI channels instead of 5. Importantly, no resting state had fewer than 4 channels for the sensorimotor ROI.

Contrary to some previous studies that used ICA to isolate sensory region alpha sources [8,15,18], we used pre-determined sensor level ROIs to improve reproducibility and reduce the potential for bias when individually selecting ICA components. Using sensor level ROIs may decrease the signal-to-noise ratio of the data; however, this approach has still been effective for observing the relationship between PAF and experimental pain [9,10].

The frequency range used to calculate PAF varies in the PAF–pain literature. The narrower 9–11 Hz band reduces the impact of 1/f noise (i.e. trend of higher power at lower frequencies) on the PAF estimations [8,117], however the narrow band may not produce representative PAF values for those with peaks near the lower or upper limits. Therefore, the wider 8–12 Hz band was used as the primary band of interest (Figure 2, plot A). All analyses were repeated with the narrower band and are displayed in Supplementary Material 3.4 and 3.6. Power was calculated using the wider 8–12 Hz band to display alpha on topographical plots.

### 4.7 Statistical analysis

Statistical analysis was conducted in RStudio (RStudio Team, 2021.9.1.372). A Guideline for Reporting Mediation Analysis (AGReMA) was followed for this article [123] and the analysis plan was pre-registered (https://osf.io/pc4rq/).

#### Mediation analysis

Using a structural equation modelling approach, a two-wave latent change score (2W-LCS) model – recommended by Valente and MacKinnon [64] – was used to assess whether nicotine changed PAF or pain (i.e. hypotheses 1–3).

Prior to modelling, a paired equivalence test satisfied that PAF_1𝑎_ (i.e. pre-gum before the first pain assessment) and PAF_1𝑏_ (i.e. pre-gum after the first pain assessment) were equivalent (𝑡(60) = 0.90, 𝑝 < .001), therefore PAF_1𝑎_ was used in the below models.

*Latent change score (LCS) specification.* Calculating the absolute difference between pre- and post-gum variables can result in regression to the mean [64]. With LCS specification, the second wave of PAF and pain (i.e. post-gum) can be set as a unit-weighted function of pre-gum and a new unobserved change variable (i.e. Δ_𝑃𝐴𝜞_ and Δ_𝑝𝑎𝑖𝑛_) to simulate subtracting pre-gum scores from post-gum scores on PAF and pain variables [64].

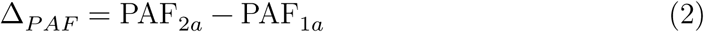

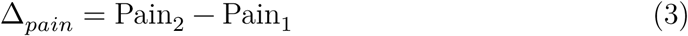

Simulation of the above equations was achieved via the following parameter constraints. Loading on Pain_2_ was fixed to one, while the mean and variance of Δ_𝑝𝑎𝑖𝑛_ were freely estimated. The path from Pain_1_ to Pain_2_ was fixed to 1, the mean (i.e. intercept) and variance of Pain_2_ were constrained to zero, and the mean and variance of Pain_1_ were freely estimated. The same approach was taken for Δ_𝑃𝐴𝜞_ . The covariances between PAF_1𝑎_ and Pain_1_ were freely estimated.

*Two-wave latent change score (2W-LCS) models.* The above latent change variables (Equation 2 and Equation 3) can then be used within 2W-LCS models to conduct mediation analysis. Two model variations were constructed to assess total, direct, and indirect effects of nicotine separately for latent change in mean ratings during PHP and CPA as outcomes (Table 4). Models were constructed in R (version 4.2.1) [124] using the lavaan package [125] with maximum likelihood estimation and 1000 bootstraps (seed: 2022) to compute standard errors. The mediation model syntax is available online (https://osf.io/pc4rq/).

**Table 4:** Overview of mediation models.

| Model | Intervention | Mediator | Outcome |
| --- | --- | --- | --- |
| 1.1 | Nicotine | Latent ΔPAF | Latent Δmean heat pain |
| 1.2 | Nicotine | Latent ΔPAF | Latent Δmean pressure pain |

*The ANCOVA-equivalent 2W-LCS model.* The first model variation used was the ANCOVA-equivalent model [64], the path diagram is displayed in Figure 7 and the below equations were used (Equation 4 and Equation 5):

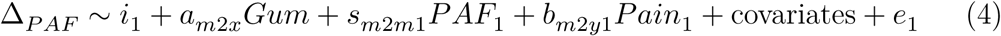

**Figure 7:**
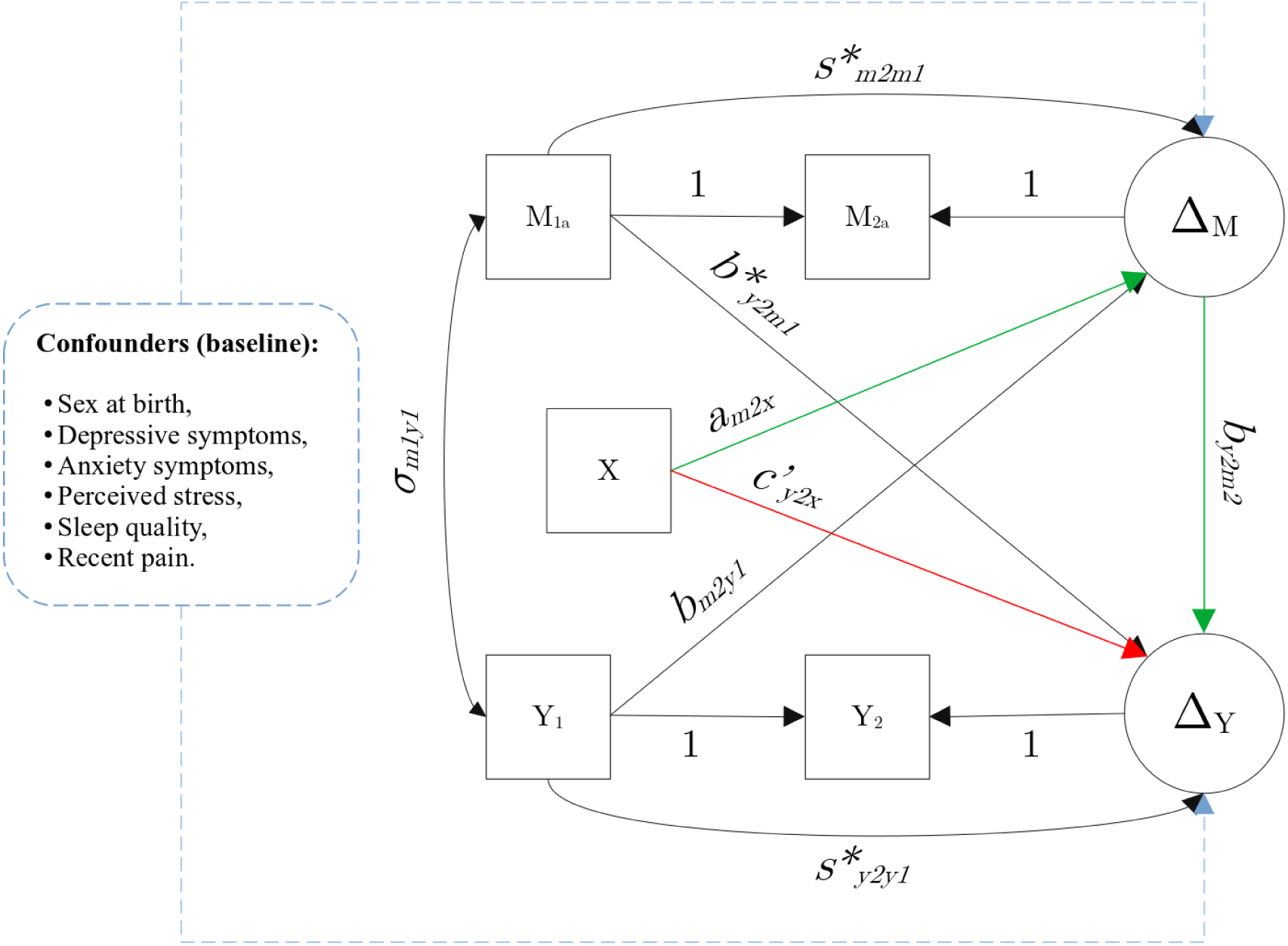
Adapted from Valente and MacKinnon [64]. The analysis of covariance (ANCOVA) equivalent pre-test/post-test control group design (two-wave) with latent change score (LCS) specification for the outcome (Y: pain) and mediating (M: PAF) variables. The direct effect of the intervention on the outcome is represented by the red line (𝑐’_𝑦2𝑥_). The intervention–mediator relationship is represented by a green line (𝑎_𝑚2𝑥_) from the intervention (i.e. X: nicotine vs. placebo) to the mediator (i.e. Δ_𝑀_). The mediator–outcome relationship is represented by another green line (𝑏_𝑦2𝑚2_) from the mediator (i.e. Δ_𝑀_) to the outcome (i.e Δ_𝑌_). Together the two green lines represent the indirect effect of the intervention on the outcome via the mediator. Time-lagged autoregression paths are represented by 𝑆∗_𝑚2𝑚1_ and 𝑆∗_𝑦2𝑦1_. Cross-lagged coupling paths are represented by 𝑏_𝑚2𝑦1_ and 𝑏∗_𝑦2𝑚1_. The covariance between Y and M at baseline is represented by 𝜎_𝑚1𝑦1_. The potential confounders of the mediator–outcome relationship were measured at baseline and are represented by the blue dashed lines. *Note.* ∗ *= latent; X = intervention (i.e. gum); M = mediator (i.e. PAF); Y = outcome (i.e. pain)*.

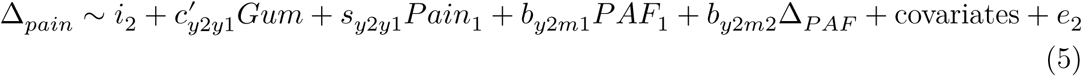

In this model, 𝑖 denotes the intercept and 𝑒 denotes error. Both 𝑠_𝑚2𝑚1_𝑀_1_ and 𝑠_𝑦2𝑦1_𝑌_1_ are time-lagged autoregressions referring to change in M and Y due to prior levels of M and Y, respectively. These autoregressions reflect that each variable influences itself over time. The 𝑏_𝑦2𝑚1_𝑀_1_ and 𝑏_𝑚2𝑦1_𝑌_1_ are cross-lagged coupling parameters, which refer to effects of previous levels of M on change in Y and effects of previous levels of Y on change in M. In other words, each variable influences the other variable at the subsequent time point. As displayed in Figure 7, X is predicting constant change in Y via the direct *c’*–path (Equation 5; 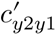) and constant change in M via the *a*–path (Equation 4; 𝑎_𝑚2𝑥_). Latent change in M is predicting later latent change in Y via the constrained *b*–path (Equation 5; 𝑏_𝑦2𝑚2_). The indirect effect was assessed by estimating the product of coefficients 𝑎𝑏 (𝑎_𝑚2𝑥_𝐸 ∗ 𝑏_𝑦2𝑚2_). The total effect was assessed by summation of the indirect and direct effects 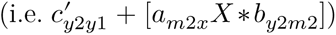.

*Difference score 2W-LCS model.* We compared the ANCOVA-equivalent model (Figure 7) to the difference score model (Figure 8); the former adjusts for the effects of pre-gum PAF and pain on post-gum PAF and pain, while the latter assumes those paths are equal to zero [64]. Removal of autoregressive (i.e. 𝑠_𝑚2𝑚1_𝑀_1_ and 𝑠_𝑦2𝑦1_𝑌_1_) and coupling parameters (i.e. 𝑏_𝑦2𝑚1_𝑀_1_ and 𝑏_𝑚2𝑦1_𝑌_1_) produces a simpler unconditional model of change [65,126]. The path diagram is displayed in Figure 8 and the below equations were used (Equation 6 and Equation 7):

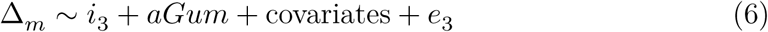

**Figure 8:**
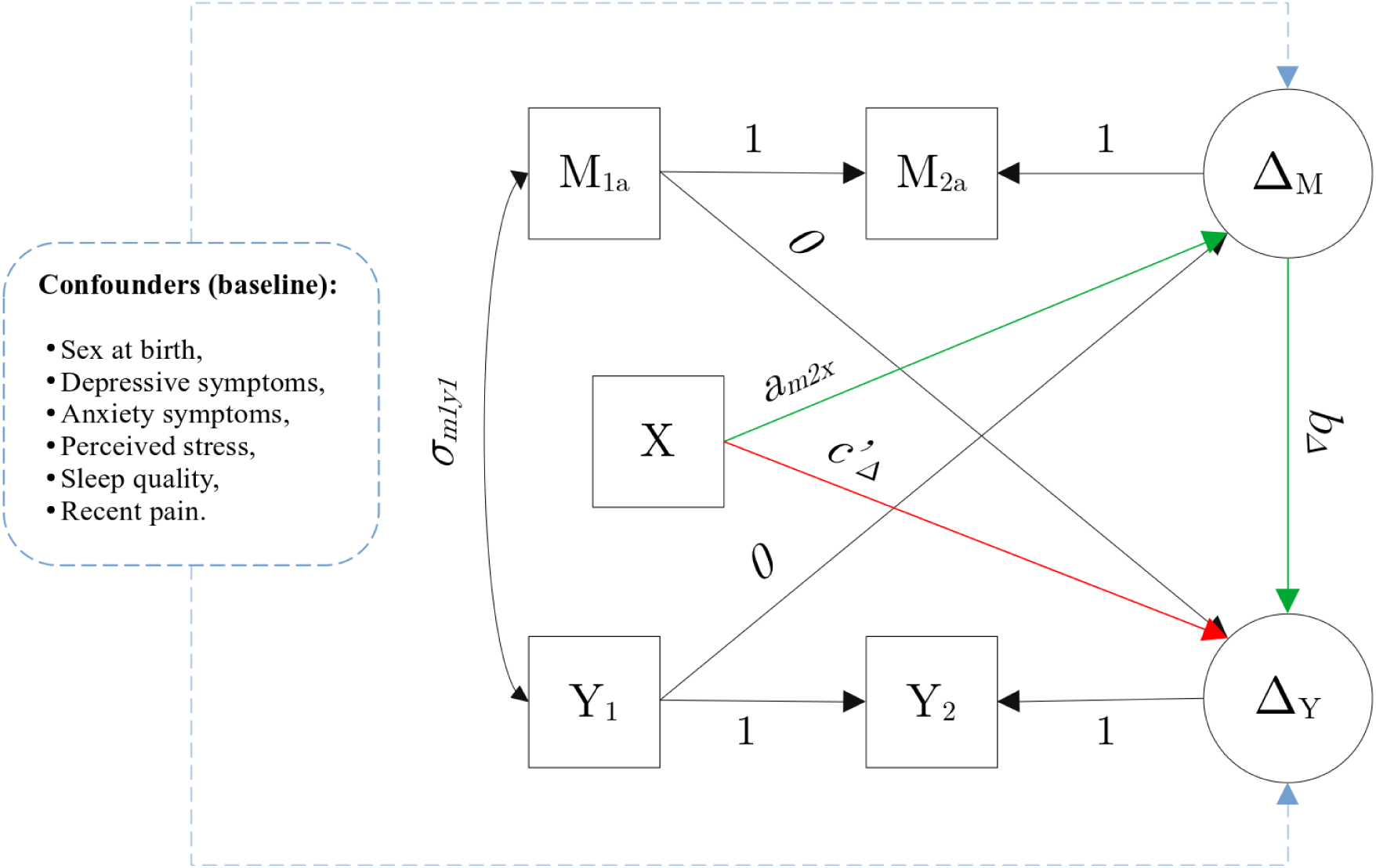
Adapted from Valante and MacKinnon [64]. The difference score model with latent change score specification for the outcome (Y) and mediating (M) variables. The direct effect of the intervention on the outcome is represented by the red line (𝑐’_Δ_). The intervention–mediator relationship is represented by a green line (𝑎_𝑚2𝑥_) from the intervention (i.e. X: nicotine vs. placebo) to the mediator (i.e. Δ_𝑀_). The mediator–outcome relationship is represented by another green line (𝑏_Δ_) from the mediator (i.e. Δ_𝑀_) to the outcome (i.e Δ_𝑌_). Together the two green lines represent the indirect effect of the intervention on the outcome via the mediator. Cross-lagged coupling paths are constrained to 0 and time-lagged autoregressions are constrained to 1. The covariance between Y and M at baseline is represented by 𝜎_𝑚1𝑦1_. The potential confounders of the mediator–outcome relationship are represented by the blue dashed lines. *Note. X = intervention (i.e. gum); M = mediator (i.e. PAF); Y = outcome (i.e. pain)*.

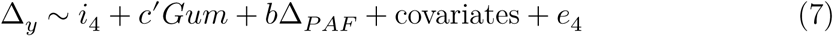

*Model fit and comparison.* Model fits were compared with 𝜒^2^statistic (p >0.05), comparative fit index (CFI), standardised root mean squared residual (SRMR), and root mean square error of approximation (RMSEA) [65,127]. For adequate model fit: CFI > 0.90, RMSEA < 0.08, and SRMR < 0.08 [128].

*Confounding.* The intervention–mediator (a–path) and intervention–outcome (c’– path) relationships are unconfounded due to the random allocation of nicotine or placebo gum. For the mediator–outcome (b–path) relationship, potential confounders were chosen by selecting variables measured prior to gum allocation that we assume to be a cause of the mediator, outcome, or both [129].

We assume that the mediator–outcome relationship is unconfounded after adjusting for the pre-determined (Supplementary Material 7.1; [130,131]) minimum sufficient set of observed confounders: sex at birth, depression symptoms, anxiety symptoms, perceived stress, sleep quality, and recent pain (Figure 7). Within the model, the variance of each covariate was freely estimated, along with their co-variance with baseline values of PAF and pain.

*Assumptions.* Multicollinearity assessment (𝑅^2^ cut-off of 0.7) did not suggest any covariates requiring removal. Using the *mvn()* function, we found violation of multivariate normality (Mardia skewness = 361.28, p <.001; Mardia kurtosis = 2.39, p = 0.017) as well as univariate normality for covariates 2, 3, 5, and 6 (W = 0.60-0.93, p < 0.0019), the impact of which was reduced by use of bootstrapping [132,133]. There was no systematically missing data and only complete datasets were used, incomplete sets were removed by listwise deletion.

*Mediation using PROCESS.* PROCESS is a statistics package in R for conducting mediation analysis [66], and is frequently used in the literature [134–136]. PROCESS analysis was conducted using and ANCOVA-equivalent model without the LCS specification, as construction of latent variables is not currently possible in this package [66]. Pain and PAF post-gum (i.e. PAF_2𝑎_ and pain_2_) were used as outcome measures rather than change in each, with PAF_1_ and pain_1_ included alongside the other covariates. The estimates produced by PROCESS were similar to the ANCOVA-equivalent 2W-LCS model in lavaan, and are displayed in the Supplementary Material 7.2.

*Cluster-based permutation analysis.* An exploratory non-parametric cluster-based permutation analysis [137,138] was conducted to assess where changes in global PAF occurred in sensor space, similar to those conduced by Furman and colleagues [80]. Tests conducted at the single electrode level were thresholded at 𝑝 < .01. If two or more spatially adjacent electrodes passed this threshold, they were grouped into a cluster, and each electrode within a cluster had to be adjacent to at least one other electrode from the overall cluster. Following the formation of clusters, test statistic values were summed across all electrodes within a cluster to obtain a composite test score for that cluster. For each potential cluster, this composite test score was then compared against a null distribution of 1000 generated scores produced by random shuffle of values for the dependent variable across conditions and participants at each electrode of the cluster. If a cluster’s composite test statistic surpassed the 99th percentile of the null distribution, that cluster was evaluated as significant.

#### PAF–pain relationship

To address whether faster sensorimotor or global PAF during pain-free resting states was associated with lower pain ratings during the first PHP and CPA models before gum chewing (i.e. hypothesis 4), correlations were used [7–10].

Bayesian correlations were conducted in R (‘BayesFactor’ package, default priors) between pain-free baseline 8–12 Hz PAF (i.e. global and sensorimotor) and pain measures (i.e. mean, max, AUC) during each model (i.e. PHP, CPA) prior to gum administration.

Median splits on PAF speed were conducted, and the difference in pain reported between PAF speed groups was assessed by independent t-tests for mean and AUC pain, with Mann Whitney-U test for max PHP pain, due to violation of normality (W = 0.89, p < .001). Results of median splits are displayed in Supplementary Material 6.1.

