## Supplementary_file_2026 for "Effects of nicotine compared to placebo gum on sensitivity to pain and mediating effects of peak alpha frequency"

### 1 Demographics

#### 1.1 Participant ancestry

Table 1: Count and percentage of participants self-reporting ancestry (N = 62). Mixed ancestry reported is presented in Table 2.

| Ancestry | Count | Percentage |
|---|---|---|
| Aboriginal or Torres Strait Islander | 0 | 0.00% |
| Other Oceanian | 1 | 1.61% |
| North-West European | 14 | 22.58% |
| Southern and Eastern European | 7 | 11.29% |
| North African and Middle Eastern | 2 | 3.23% |
| South-East Asian | 12 | 19.35% |
| North-East Asian | 5 | 8.06% |
| Southern and Central Asian | 6 | 9.68% |
| Peoples of the Americas | 3 | 4.84% |
| Sub-Saharan African | 1 | 1.61% |
| Prefer to self-describe | 3 | 4.84% |
| **Mixed** | 8 | 12.90% |

Table 2: Description and count of the mixed ancestry reported by participants (n = 8)

| Ancestry | Count |
|---|---|
| North-West European and Oceanian | 2 |
| North-West European and Southern and Eastern European | 2 |
| North-West European and South-East Asian | 1 |
| North-West European and North African and Middle Eastern | 1 |
| North-West European and Southern and Central Asian | 1 |
| South-East Asian and North-East Asian | 1 |

#### 2 Chewing gum experience and blinding

Participants were asked to rate their perception of the gum on several scales, to assess whether the nicotine and placebo gums were perceptually matched (Figure 1). Participants in the placebo group described their gum as a stronger flavour than those in the nicotine gum group ($BF_{10} = 1.68$), while the gums were reported as similarly pleasant ($BF_{10} = 0.70$), juicy ($BF_{10} = 0.26$), sweet ($BF_{10} = 0.43$), spicy ($BF_{10} = 0.37$), and bitter ($BF_{10} = 0.34$).

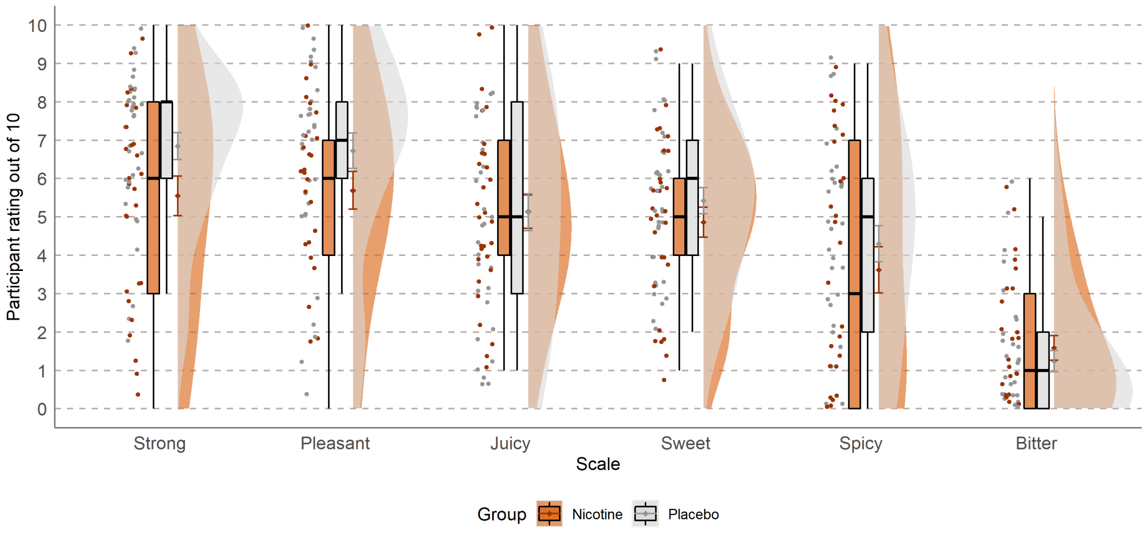

Figure 1: Participants' experience of the gum flavour, split into nicotine (orange, n = 29) and placebo (grey, n = 33) groups. Participants were asked, for example, to 'indicate on the scale how *sweet* the gum was:' on a scale of 0 "not at all" to 10 being "very *sweet*"

Eleven of 62 participants reported feeling some side effects from chewing gum. The proportion of participants reporting side effects from gum chewing was higher for the nicotine group (9 yes, 20 no) than the placebo group (2 yes, 31 no; $\chi^2(1) = 5.0, p = .025$). The most commonly reported side effect was a sore throat/mouth (5 participants), followed by tingling (4 participants), dizziness and nausea (3 participants for each), and other (1 participant; "Slight numbing/drying of the mucus membrane in the mouth").

A chi-squared test found no evidence for differences in the proportion of participants guessing at having nicotine, placebo, or not being sure between the two gum groups, $\chi^2(2) = 3.34, p = 0.19$ (Figure 2).

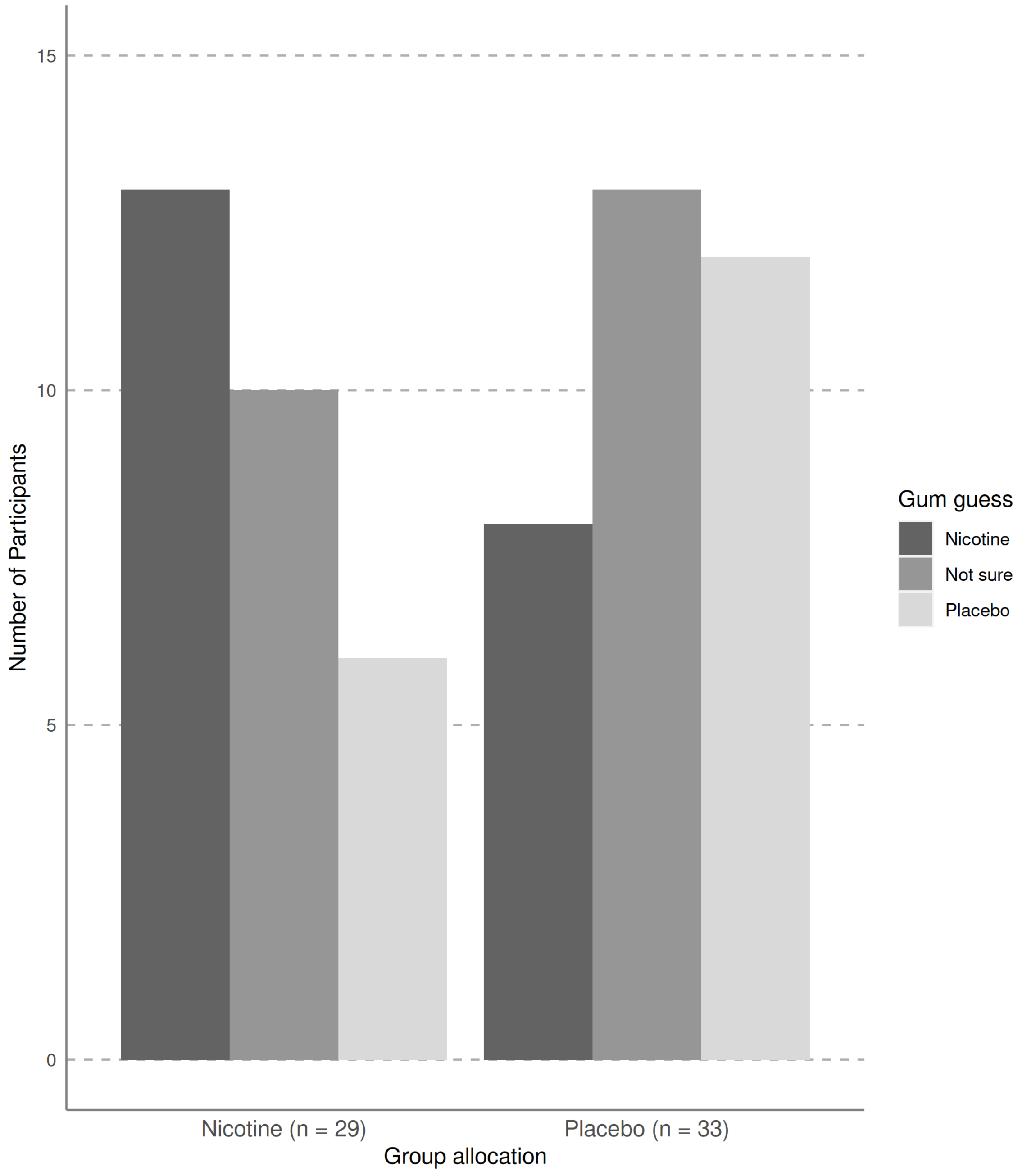

Figure 2: Participants' guess at which gum they were given after being debriefed. Actual group participants were allocated to on the x-axis, with the gum the participants' guessed they had been given out of nicotine (dark grey), placebo (light grey), or if they were not sure (mid-grey), $\chi^2(2) = 3.34, p = 0.19$

### 3 Mediation analysis

#### 3.1 ANCOVA-equivalent 2W-LCS model

The ANCOVA-equivalent 2W-LCS model examined whether latent change in global PAF (8–12 Hz) mediated any relationship between nicotine and change in pain ratings recorded during PHP and CPA. The model was refined with the addition of a specified corvariance between covariate 2 (i.e. depression symptoms) and covariate 3 (i.e. anxiety symptoms) by a recommendation from the *modindices()* function in the Lavaan package ($\delta\chi^2(-1) = -20.75, p < .001, \delta$CFI = 0.073, $\delta$RMSEA = -0.08, $\delta$SRMR = -0.023). Unstandardised regression coefficients are provided for PHP (Table 5) and CPA (Table 6).

For the ANCOVA-equivalent 2W-LCS model for PHP ratings ($\chi^2(20) = 23.72, p = 0.26$, CFI = 0.987 RMSEA = 0.056, SRMR = 0.078), the variance explained was $r^2 = 0.257$ for the nested model of change in PAF and $r^2 = 0.349$ for change in PHP ratings.

For the ANCOVA-equivalent 2W-LCS model for CPA ratings ($\chi^2(20) = 22.64, p = 0.31$, CFI = 0.987 RMSEA = 0.046, SRMR = 0.077), the variance explained was similar to the PHP model for change in PAF($r^2 = 0.244$), but lower than the PHP for change in CPA ratings ($r^2 = 0.219$).

There were no major differences in the corresponding estimates produced by the ANCOVA-equivalent and difference score models. However, there are several parameters freely estimated in the ANCOVA equivalent model that are not in the difference score model, as they are held constant: PAF and pain stability (i.e. auto-regressions), and the effect of baseline PAF on change in pain and baseline pain on change in PAF (i.e. cross-lagged). The estimate of PAF stability when controlling for gum group, pain, and confounding was significant, and expectedly similar, for both the PHP ($b$ = 1.02, p < .001, bootstrapped 95% CI: [0.92, 1.11]) and CPA models ($b$ = 1.03, p < .001, bootstrapped 95% CI: [0.95, 1.11]) . The estimate of pain stability when controlling for gum group, PAF, and confounding was also significant for both the PHP ($b$ = 1.02, p < .001, bootstrapped 95% CI: [0.92, 1.15]) and CPA models ($b$ = 0.80, p < .001, bootstrapped 95% CI: [0.48, 1.15]). Meanwhile, levels of PAF at baseline did not significantly impact change in PAF, and levels of pain at baseline did not significantly affect change in pain (Table 3 and Table 4). Taken together, this means that PAF speed and PHP ratings are very stable, while CPA ratings showed a general trend to decrease when controlling for all other factors, including intervention, regardless of the initial levels of pain or PAF speed. The significant stability of PAF and pain shown in this model, justifies use of the simpler difference score model, which fixes the stability of PAF and pain to 1.

Table 3: Mediation model (ANCOVA-equivalent 2W-LCS) using global peak alpha frequency (PAF; 8–12 Hz) and mean phasic heat pain (PHP) ratings. Covariate/confounding variables: sex (male/female), depressive symptoms, anxiety symptoms, perceived stress, sleep quality, and recent pain. Indirect (a+b path), direct (c'-path), and total effects of nicotine on change in pain calculated from model estimates. Latent change score factor is removed to retrieve ANCOVA estimates of PAF and pain stability ($S_{PAF}$ and $S_{Pain}$), and effect of baseline PAF on pain post-gum ($B_{y2m1}$).

| | | | label | estimate | SE | $Z$ | $p$-value | CI lower | CI upper |
|---|---|---|---|---|---|---|---|---|---|
| $PAF^1$ | ~~ | $Pain^1$ | | -0.110 | 0.136 | -0.809 | 0.419 | -0.371 | 0.172 |
| $\Delta_{PAF}$ | ~1 | *(intercept)* | | 0.042 | 0.458 | 0.092 | 0.926 | -0.825 | 1.024 |
| $\Delta_{PAF}$ | ~ | Gum | $a_{m2x}$ | 0.092 | 0.037 | 2.474 | 0.013* | 0.020 | 0.164 |
| $\Delta_{PAF}$ | ~ | $PAF^1$ | $s_{m2m1}$ | 0.017 | 0.046 | 0.365 | 0.715 | -0.077 | 0.108 |
| $\Delta_{PAF}$ | ~ | $Pain^1$ | $b_{m2y1}$ | -0.010 | 0.010 | -1.053 | 0.292 | -0.031 | 0.008 |
| $\Delta_{PAF}$ | ~ | sex | | -0.089 | 0.042 | -2.133 | 0.033 | -0.180 | -0.012 |
| $\Delta_{PAF}$ | ~ | depression | | -0.035 | 0.029 | -1.213 | 0.225 | -0.085 | 0.033 |
| $\Delta_{PAF}$ | ~ | anxiety | | 0.034 | 0.025 | 1.339 | 0.180 | -0.024 | 0.087 |
| $\Delta_{PAF}$ | ~ | stress | | -0.005 | 0.006 | -0.920 | 0.358 | -0.017 | 0.005 |
| $\Delta_{PAF}$ | ~ | sleep quality | | -0.014 | 0.013 | -1.081 | 0.280 | -0.038 | 0.015 |
| $\Delta_{PAF}$ | ~ | recent pain | | 0.018 | 0.024 | 0.753 | 0.452 | -0.030 | 0.061 |
| $\Delta_{Pain}$ | ~1 | *(intercept)* | | 1.649 | 2.899 | 0.569 | 0.569 | -4.049 | 7.217 |
| $\Delta_{Pain}$ | ~ | Gum | $c'_{y2x}$ | -0.685 | 0.290 | -2.358 | 0.018* | -1.319 | -0.186 |
| $\Delta_{Pain}$ | ~ | $\Delta_{PAF}$ | $b_{y2m2}$ | 0.288 | 0.867 | 0.332 | 0.740 | -1.344 | 2.064 |
| $\Delta_{Pain}$ | ~ | $PAF^1$ | $b_{y2m1}$ | -0.034 | 0.275 | -0.123 | 0.902 | -0.557 | 0.528 |
| $\Delta_{Pain}$ | ~ | $Pain^1$ | $s_{y2y1}$ | 0.019 | 0.057 | 0.333 | 0.739 | -0.085 | 0.148 |
| $\Delta_{Pain}$ | ~ | sex | | 0.165 | 0.319 | 0.517 | 0.605 | -0.422 | 0.832 |
| $\Delta_{Pain}$ | ~ | depression | | 0.303 | 0.173 | 1.746 | 0.081 | -0.047 | 0.651 |
| $\Delta_{Pain}$ | ~ | anxiety | | -0.071 | 0.165 | -0.433 | 0.665 | -0.523 | 0.128 |
| $\Delta_{Pain}$ | ~ | stress | | -0.094 | 0.049 | -1.908 | 0.056 | -0.188 | 0.004 |
| $\Delta_{Pain}$ | ~ | sleep quality | | -0.007 | 0.073 | -0.094 | 0.925 | -0.149 | 0.136 |
| $\Delta_{Pain}$ | ~ | recent pain | | -0.174 | 0.134 | -1.300 | 0.194 | -0.425 | 0.107 |
| $S_{PAF}$ | := | $s_{m2m1} + 1$ | $S_m$ | 1.017 | 0.046 | 22.267 | 0.000 | 0.923 | 1.108 |
| $S_{Pain}$ | := | $s_{y2y1} + 1$ | $S_y$ | 1.019 | 0.057 | 17.907 | 0.000 | 0.915 | 1.148 |
| $B_{y2m1}$ | := | $b_{y2m1} - b_{y2m2}$ | $B_y 2m1$ | -0.321 | 0.892 | -0.360 | 0.719 | -2.138 | 1.393 |
| Indirect | := | $a_{m2x} * b_{y2m2}$ | indirect | 0.026 | 0.090 | 0.294 | 0.769 | -0.107 | 0.256 |

| | | | label | estimate | SE | $Z$ | $p$-value | CI lower | CI upper |
|---|---|---|---|---|---|---|---|---|---|
| Direct | := | $c'_{y2x}$ | direct | -0.685 | 0.290 | -2.357 | 0.018 | -1.319 | -0.186 |
| Total | := | $c'_{y2x}$ + ($a_{m2x}$*$b_{y2m2}$) | total | -0.658 | 0.285 | -2.312 | 0.021* | -1.222 | -0.156 |

*Note. SE = standard error; CI = 95% bootstrapped confidence interval;* $\Delta_{PAF}$ *= change in mediator (i.e. peak alpha frequency);* $\Delta_{Pain}$ *= change in outcome (i.e. CPA pain);* $PAF^1$ *= baseline PAF, pre-gum; PAF^2 = PAF post-gum;* $Pain^1$ *baseline mean PHP rating, pre-gum;* $Pain^2$ *= mean PHP rating, post-gum;* $S_{PAF}$ *= stability of PAF;* $S_{PAF}$ *= stability of PAF.*

Table 4: Mediation model (ANCOVA-equivalent 2W-LCS) using global peak alpha frequency (PAF; 8–12 Hz) and mean cuff pressure algometry (CPA) pain ratings. Covariate/confounding variables: sex (male/female), depressive symptoms, anxiety symptoms, perceived stress, sleep quality, and recent pain. Indirect (a+b path), direct (c'-path), and total effects of nicotine on change in pain calculated from model estimates. Latent change score factor is removed to retrieve ANCOVA estimates of PAF and pain stability ($S_{PAF}$ and $S_{Pain}$), and effect of baseline PAF on pain post-gum ($B_{y2m1}$).

| | | | label | estimate | SE | $Z$ | $p$-value | CI lower | CI upper |
|---|---|---|---|---|---|---|---|---|---|
| $PAF^1$ | ~~ | $Pain^1$ | | -0.030 | 0.088 | -0.347 | 0.728 | -0.200 | 0.152 |
| $\Delta_{PAF}$ | ~1 | *(intercept)* | | -0.139 | 0.404 | -0.345 | 0.730 | -0.927 | 0.706 |
| $\Delta_{PAF}$ | ~ | Gum | $a_{m2x}$ | 0.094 | 0.037 | 2.516 | 0.012* | 0.018 | 0.166 |
| $\Delta_{PAF}$ | ~ | $PAF^1$ | $s_{m2m1}$ | 0.027 | 0.042 | 0.649 | 0.516 | -0.055 | 0.112 |
| $\Delta_{PAF}$ | ~ | $Pain^1$ | $b_{m2y1}$ | 0.000 | 0.013 | -0.018 | 0.985 | -0.026 | 0.025 |
| $\Delta_{PAF}$ | ~ | sex | | -0.072 | 0.036 | -1.995 | 0.046 | -0.152 | -0.003 |
| $\Delta_{PAF}$ | ~ | depressive | | -0.031 | 0.026 | -1.181 | 0.238 | -0.073 | 0.035 |
| $\Delta_{PAF}$ | ~ | anxiety | | 0.029 | 0.024 | 1.205 | 0.228 | -0.029 | 0.070 |
| $\Delta_{PAF}$ | ~ | stress | | -0.004 | 0.005 | -0.803 | 0.422 | -0.015 | 0.006 |
| $\Delta_{PAF}$ | ~ | sleep quality | | -0.014 | 0.013 | -1.053 | 0.292 | -0.037 | 0.015 |
| $\Delta_{PAF}$ | ~ | recent pain | | 0.017 | 0.022 | 0.788 | 0.431 | -0.028 | 0.059 |
| $\Delta_{Pain}$ | ~1 | *(intercept)* | | -9.414 | 5.837 | -1.613 | 0.107 | -20.165 | 3.692 |
| $\Delta_{Pain}$ | ~ | Gum | $c'_{y2x}$ | 0.151 | 0.457 | 0.330 | 0.741 | -0.764 | 1.000 |
| $\Delta_{Pain}$ | ~ | $\Delta_{PAF}$ | $b_{y2m2}$ | -1.523 | 1.816 | -0.839 | 0.402 | -5.330 | 1.755 |
| $\Delta_{Pain}$ | ~ | $PAF^1$ | $b_{y2m1}$ | 0.937 | 0.532 | 1.762 | 0.078 | -0.249 | 1.971 |
| $\Delta_{Pain}$ | ~ | $Pain^1$ | $s_{y2y1}$ | -0.205 | 0.177 | -1.160 | 0.246 | -0.524 | 0.151 |
| $\Delta_{Pain}$ | ~ | sex | | 0.744 | 0.430 | 1.730 | 0.084 | -0.098 | 1.575 |

| | | | label | estimate | SE | $Z$ | $p$-value | CI lower | CI upper |
|---|---|---|---|---|---|---|---|---|---|
| $\Delta_{Pain}$ | ~ | depressive | | 0.111 | 0.337 | 0.329 | 0.742 | -0.585 | 0.727 |
| $\Delta_{Pain}$ | ~ | anxiety | | -0.144 | 0.384 | -0.375 | 0.707 | -1.033 | 0.355 |
| $\Delta_{Pain}$ | ~ | stress | | -0.062 | 0.059 | -1.044 | 0.296 | -0.181 | 0.054 |
| $\Delta_{Pain}$ | ~ | sleep quality | | 0.046 | 0.131 | 0.353 | 0.724 | -0.205 | 0.299 |
| $\Delta_{Pain}$ | ~ | recent pain | | 0.078 | 0.254 | 0.308 | 0.758 | -0.389 | 0.572 |
| $S_{PAF}$ | := | $s_{m2m1} + 1$ | $S_m$ | 1.027 | 0.042 | 24.293 | 0.000 | 0.945 | 1.112 |
| $S_{Pain}$ | := | $s_{y2y1} + 1$ | $S_y$ | 0.795 | 0.177 | 4.495 | 0.000 | 0.476 | 1.151 |
| $B_y 2m1$ | := | $b_{y2m1} - b_{y2m2}$ | $B_{y2m1}$ | 2.460 | 1.921 | 1.281 | 0.200 | -1.031 | 6.447 |
| Indirect | := | $a_{m2x}*b_{y2m2}$ | indirect | -0.143 | 0.189 | -0.757 | 0.449 | -0.613 | 0.157 |
| Direct | := | $c'_{y2x}$ | direct | 0.151 | 0.457 | 0.330 | 0.741 | -0.764 | 1.000 |
| Total | := | $c'_{y2x} + (a_{m2x}*b_{y2m2})$ | total | 0.008 | 0.443 | 0.018 | 0.986 | -0.946 | 0.820 |

*Note. SE = standard error; CI = 95% bootstrapped confidence interval; $\Delta_{PAF}$ = change in mediator (i.e. peak alpha frequency); $\Delta_{Pain}$ = change in outcome (i.e. CPA pain); $PAF^1$ = baseline PAF, pre-gum; PAF^2 = PAF post-gum; $Pain^1$ baseline mean CPA rating, pre-gum; $Pain^2$ = mean CPA rating, post-gum; $S_{PAF}$ = stability of PAF; $S_{PAF}$ = stability of PAF.*

#### 3.2 Difference score 2W-LCS model estimates

##### *PHP mediation table*

Table 5: Overview of mediation model (difference score 2W-LCS) using global peak alpha frequency (PAF; 8–12 Hz) and mean pain ratings during phasic heat pain (PHP). Covariate/confounding variables are sex (male/female), depressive symptoms, anxiety symptoms, perceived stress, sleep quality, and recent pain. Indirect (a+b–path), direct (c'–path), and total effects of nicotine on change in pain are calculated from model estimates.

| | | | label | estimate | SE | $Z$ | $p$-value | CI lower | CI upper |
|---|---|---|---|---|---|---|---|---|---|
| $PAF^1$ | ~~ | $Pain^1$ | | -0.110 | 0.136 | -0.809 | 0.419 | -0.371 | 0.172 |
| $\Delta_{PAF}$ | ~1 | *(intercept)* | | 0.136 | 0.117 | 1.158 | 0.247 | -0.114 | 0.355 |
| $\Delta_{PAF}$ | ~ | Gum | $a_{m2x}$ | 0.085 | 0.036 | 2.356 | 0.018* | 0.017 | 0.154 |
| $\Delta_{PAF}$ | ~ | sex | | -0.077 | 0.037 | -2.089 | 0.037 | -0.156 | -0.011 |
| $\Delta_{PAF}$ | ~ | depression | | -0.030 | 0.027 | -1.102 | 0.270 | -0.074 | 0.038 |
| $\Delta_{PAF}$ | ~ | anxiety | | 0.029 | 0.024 | 1.198 | 0.231 | -0.033 | 0.068 |
| $\Delta_{PAF}$ | ~ | stress | | -0.005 | 0.005 | -0.835 | 0.404 | -0.016 | 0.006 |
| $\Delta_{PAF}$ | ~ | sleep quality | | -0.013 | 0.013 | -0.990 | 0.322 | -0.037 | 0.016 |
| $\Delta_{PAF}$ | ~ | recent pain | | 0.017 | 0.023 | 0.737 | 0.461 | -0.031 | 0.061 |
| $\Delta_{Pain}$ | ~1 | *(intercept)* | | 1.457 | 0.841 | 1.732 | 0.083 | -0.279 | 3.031 |
| $\Delta_{Pain}$ | ~ | Gum | $c'_{y2x}$ | -0.667 | 0.288 | -2.314 | 0.021* | -1.282 | -0.163 |
| $\Delta_{Pain}$ | ~ | $\Delta_{PAF}$ | $b_{y2m2}$ | 0.227 | 0.817 | 0.278 | 0.781 | -1.282 | 1.911 |
| $\Delta_{Pain}$ | ~ | sex | | 0.139 | 0.282 | 0.493 | 0.622 | -0.381 | 0.727 |
| $\Delta_{Pain}$ | ~ | depression | | 0.292 | 0.159 | 1.842 | 0.066 | -0.039 | 0.595 |
| $\Delta_{Pain}$ | ~ | anxiety | | -0.060 | 0.139 | -0.430 | 0.667 | -0.428 | 0.119 |
| $\Delta_{Pain}$ | ~ | stress | | -0.095 | 0.048 | -1.981 | 0.048 | -0.186 | -0.000047 |
| $\Delta_{Pain}$ | ~ | sleep quality | | -0.010 | 0.070 | -0.140 | 0.888 | -0.148 | 0.129 |
| $\Delta_{Pain}$ | ~ | recent pain | | -0.171 | 0.127 | -1.350 | 0.177 | -0.421 | 0.095 |
| Indirect | := | $a_{m2x}*b_{y2m2}$ | indirect | 0.019 | 0.079 | 0.246 | 0.805 | -0.093 | 0.232 |
| Direct | := | $c'_{y2x}$ | direct | -0.667 | 0.288 | -2.313 | 0.021 | -1.282 | -0.163 |
| Total | := | $c'_{y2x}$ + ($a_{m2x}*b_{y2m2}$) | total | -0.648 | 0.284 | -2.276 | 0.023* | -1.208 | -0.154 |

*Note. SE = standard error; CI = 95% bootstrapped confidence interval;* $\Delta_{PAF}$ *= change in mediator* *(i.e. peak alpha frequency);* $\Delta_{Pain}$ *= change in outcome (i.e. CPA pain);* $PAF^1$ *= baseline PAF, pre-gum;* $Pain^1$ *baseline mean PHP rating, pre-gum.*

##### *CPA mediation table*

Table 6: Overview of mediation model (difference score 2W-LCS) using global peak alpha frequency (PAF; 8–12 Hz) and mean pain ratings during cuff pressure algometry (CPA). Covariate/confounding variables are sex (male/female), depressive symptoms, anxiety symptoms, perceived stress, sleep quality, and recent pain. Indirect (a+b–path), direct (c'–path), and total effects of nicotine on change in pain are calculated from model estimates.

| | | | label | estimate | SE | *Z* | *p*-value | CI lower | CI upper |
|---|---|---|---|---|---|---|---|---|---|
| $PAF^1$ | ~~ | $Pain^1$ | | -0.030 | 0.088 | -0.347 | 0.728 | -0.200 | 0.152 |
| $\Delta_{PAF}$ | ~1 | *(intercept)* | | 0.137 | 0.119 | 1.150 | 0.250 | -0.103 | 0.372 |
| $\Delta_{PAF}$ | ~ | Gum | $a_{m2x}$ | 0.089 | 0.035 | 2.540 | 0.011 | 0.018 | 0.158 |
| $\Delta_{PAF}$ | ~ | sex | | -0.076 | 0.034 | -2.208 | 0.027 | -0.150 | -0.010 |
| $\Delta_{PAF}$ | ~ | depressive | | -0.031 | 0.026 | -1.203 | 0.229 | -0.074 | 0.034 |
| $\Delta_{PAF}$ | ~ | anxiety | | 0.028 | 0.023 | 1.207 | 0.228 | -0.029 | 0.067 |
| $\Delta_{PAF}$ | ~ | stress | | -0.004 | 0.005 | -0.818 | 0.414 | -0.015 | 0.005 |
| $\Delta_{PAF}$ | ~ | sleep quality | | -0.014 | 0.012 | -1.087 | 0.277 | -0.036 | 0.014 |
| $\Delta_{PAF}$ | ~ | recent pain | | 0.016 | 0.021 | 0.755 | 0.450 | -0.027 | 0.055 |
| $\Delta_{Pain}$ | ~1 | *(intercept)* | | -1.179 | 1.419 | -0.831 | 0.406 | -3.916 | 1.679 |
| $\Delta_{Pain}$ | ~ | Gum | $c'_{y2x}$ | -0.169 | 0.422 | -0.401 | 0.689 | -1.070 | 0.591 |
| $\Delta_{Pain}$ | ~ | $\Delta_{PAF}$ | $b_{y2m2}$ | -1.281 | 1.838 | -0.697 | 0.486 | -5.205 | 2.113 |
| $\Delta_{Pain}$ | ~ | sex | | 0.728 | 0.435 | 1.675 | 0.094 | -0.121 | 1.546 |
| $\Delta_{Pain}$ | ~ | depressive | | 0.188 | 0.346 | 0.541 | 0.588 | -0.542 | 0.819 |
| $\Delta_{Pain}$ | ~ | anxiety | | -0.208 | 0.395 | -0.526 | 0.599 | -1.138 | 0.313 |
| $\Delta_{Pain}$ | ~ | stress | | -0.056 | 0.065 | -0.864 | 0.387 | -0.177 | 0.077 |
| $\Delta_{Pain}$ | ~ | sleep quality | | 0.083 | 0.145 | 0.569 | 0.569 | -0.191 | 0.365 |
| $\Delta_{Pain}$ | ~ | recent pain | | 0.075 | 0.238 | 0.316 | 0.752 | -0.397 | 0.546 |
| Indirect | := | $a_{m2x} * b_{y2m2}$ | indirect | -0.114 | 0.182 | -0.626 | 0.531 | -0.613 | 0.181 |
| Direct | := | $c'_{y2x}$ | direct | -0.169 | 0.422 | -0.401 | 0.689 | -1.070 | 0.591 |
| Total | := | $c'_{y2x} + (a_{m2x} * b_{y2m2})$ | total | -0.283 | 0.418 | -0.676 | 0.499 | -1.128 | 0.468 |

*Note. SE = standard error; CI = 95% bootstrapped confidence interval;* $\Delta_{PAF}$ *= change in mediator (i.e. peak alpha frequency);* $\Delta_{Pain}$ *= change in outcome (i.e. CPA pain);* $PAF^1$ *= baseline PAF, pre-gum;* $Pain^1$ *baseline mean CPA rating, pre-gum.*

##### 3.3 Exploratory analysis of the influence of perceived stress on the effects of nicotine on change in PHP ratings

Due to the significant estimated effects of perceived stress on change in PHP ratings in the 2W-LCS mediation model, we also explored post-hoc effects of stress on change in PHP ratings. We found that there is strong evidence for a negative correlation between stress and change in PHP rating within the nicotine group (n = 28, $r = -0.39$, $BF_{10} = 13.65$; Figure 3) that is not present in the placebo group, with equivocal evidence (n = 32, $r = -0.14$, $BF_{10} = 0.46$). This suggests that those with higher baseline stress who had nicotine gum experienced greater decreases in PHP ratings. Note that there was less, but still sufficient evidence for this relationship within the nicotine group when the participant who was a potential outlier for change in PHP rating was removed (n = 27, $r = -0.32$, $BF_{10} = 1.45$).

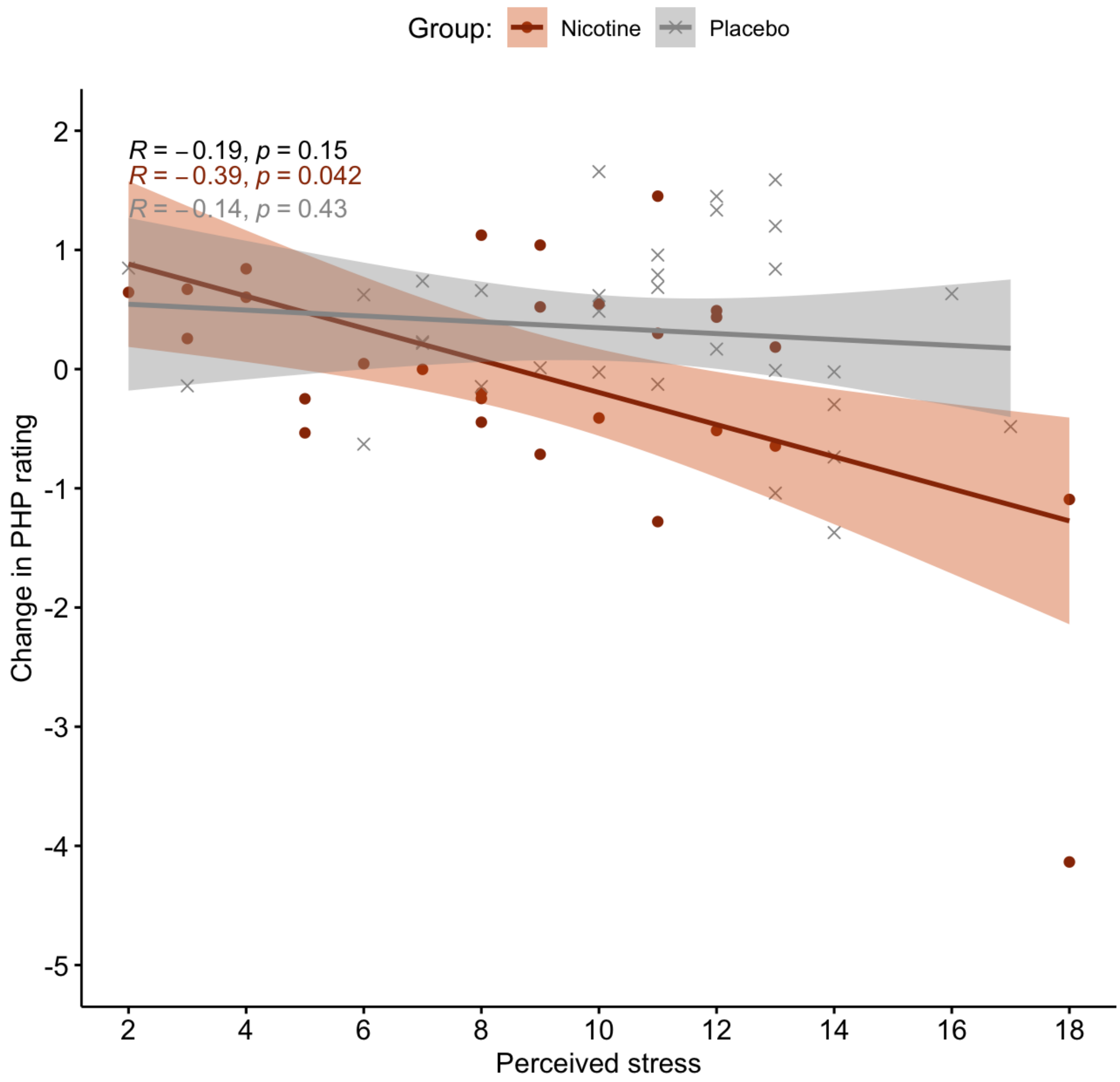

Figure 3: Spearman correlations of baseline perceived stress with change in phasic heat pain (PHP) ratings, suggest strong evidence for a negative relationship for the nicotine gum group in orange (n=28; $BF_{10} = 13.65$) but not for the placebo group in grey ($n = 32$; $BF_{10} = 0.46$). Regression lines and 95% confidence intervals.

##### 3.4 Exploratory analysis of the influence of sex on the effects of nicotine on change in PAF

***Assessment of sex at birth differences at baseline***

Due to the significant estimated effects of sex on change in PAF in the 2W-LCS mediation model, we also explored post-hoc effects of sex on change in PAF. Similar severity of pain was experienced by males (M = 4.40) and females (M = 4.75) during the first CPA assessment, with anecdotal evidence for the null hypothesis, $t(55.58) = 0.91, p = .37$, $BF_{10} = 0.37$. However, for the first PHP, females reported higher pain (M = 6.29) than males, with anecdotal evidence for this difference (M = 4.99; $t(60) = 2.25, p = .028$, $BF_{10} = 2.11$) and female participants required less pressure, with very strong evidence for this difference (M = 235.26 mmHg) than males (356.49 mmHg) to produce a 4/10 rating of pain, $t(60) = -4.52, p < .001$, $BF_{10} = 642.93$.

Table 7: Mean ($\pm SD$) baseline demographics, peak alpha frequency (PAF; 8–12 Hz), and pain of whole sample (N = 62) and split by sex. Wilcoxon or t-test used were appropriate.

| | N = 62 | Female (n = 32) | Male (n = 30) | Sex statistic | $BF_{10}$ |
|---|---|---|---|---|---|
| Age (years) | 26.45 (±7.28) | 24.53 (±6.52) | 28.5 (±7.59) | $W = 320, p = .024$ | |
| Weight (kg) | 68.16 (±13.42) | 59.50 (±8.76) | 77.39 (±11.21) | $t = -7.03, p < .001$ | 3714724 |
| Depressive | 0.53 (±0.97) | 0.69 (±1.00) | 0.37 (±0.93) | $W = 585, p = .065$ | |
| Anxiety | 0.69 (±1.06) | 0.91 (±1.25) | 0.47 (±0.78) | $W = 578.5, p = .12$ | |
| Stress | 9.82 (±3.81) | 10.41 (±3.46) | 9.20 (±4.12) | $t = 1.25, p = .22$ | 0.50 |
| Sleep quality | 6.97 (±1.77) | 6.88 (±1.68) | 7.07 (±1.89) | $t = -0.42, p = .67$ | 0.28 |
| PAF (global) | 9.956 (±0.405) | 10.019 (±0.441) | 9.888 (±0.357) | $t = 1.27, p = .21$ | 0.51 |
| PAF (sensorimotor) | 9.896 (±0.373) | 9.928 (±0.415) | 9.863 (±0.325) | $t = 0.68, p = .50$ | 0.31 |
| Heat pain threshold | 41.27 (±2.59) | 40.75 (±2.62) | 41.83 (±2.48) | $t = -1.67, p = .10$ | 0.83 |
| Pressure threshold | 293.92 (±121.21) | 235.26 (±96.21) | 356.49 (±114.72) | $t = -4.52, p < .001^{***}$ | 642.93 |
| PHP | 5.66 (±2.34) | 6.29 (±2.19) | 4.99 (±2.34) | $t = 2.25, p = .028$ | 2.11 |
| CPA | 4.58 (±1.53) | 4.75 (±1.77) | 4.40 (±1.24) | $t = 0.91, p = .37$ | 0.37 |

*Note. PAF = peak alpha frequency; PHP = phasic heat pain; CPA = cuff pressure algometry.*

***Effects of nicotine were greater in female than male participants***

RM-ANOVAs to assess the effects of time point and gum group on global PAF were conducted separately for male and female participants. For male participants (Figure 4: plot A), interactions

$(F(1,28) = 1.47, p = .24, \eta^2_G = 0.002$, $\mathrm{BF}_{10} = 0.11)$ and main effects of time point $(F(1,28) = 0.26, p = .61, \eta^2_G = 0.00036$, $\mathrm{BF}_{10} = 0.27)$ and gum $(F(1,28) = 0.58, p = .45, \eta^2_G = 0.020$, $\mathrm{BF}_{10} = 0.43)$ on global PAF were non-significant.

For female participants (Figure 4: plot B), there was a statistically significant interaction between the effects of nicotine and time point on global PAF $(F(1,30) = 6.92, p = .013, \eta^2_G = 0.005$, $\mathrm{BF}_{10} = 0.17)$ albeit with moderate evidence for the null hypothesis. Main effects showed that the time point had an independent effect on global PAF in females $(F(1,30) = 7.90, p = .009, \eta^2_G = 0.006$, $\mathrm{BF}_{10} = 0.29)$, but gum group did not $(F(1,30) = 0.99, p = .33, \eta^2_G = 0.031$, $\mathrm{BF}_{10} = 0.59)$, with moderate and anecdotal evidence for the null hypothesis, respectively.

Similarly to when looking at the whole sample, post-hoc pairwise comparisons assessing effects within each gum group for females suggested an effect of time (i.e. pre-post gum) on PAF within the nicotine group $(t(13) = -3.19, p = .007$, 95% CI: [-0.21, -0.041], $\mathrm{BF}_{10} = 0.36)$ that is not seen within the placebo group $(t(17) = -0.16, p = .88$, 95% CI: [-0.061, 0.053], $\mathrm{BF}_{10} = 0.24)$. The indicates that PAF increased within the female nicotine gum group after chewing but not the female placebo group. Post-hoc pairwise comparisons assessing effects within each time point do not suggest global PAF differences in females between groups pre-gum $(t(29.9) = -1.42, p = 0.17$, 95% CI: [-0.52, 0.094], $\mathrm{BF}_{10} = 0.70)$ or post-gum $(t(29.4) = -0.60, p = 0.56$, 95% CI: [-0.41, 0.23], $\mathrm{BF}_{10} = 0.39)$.

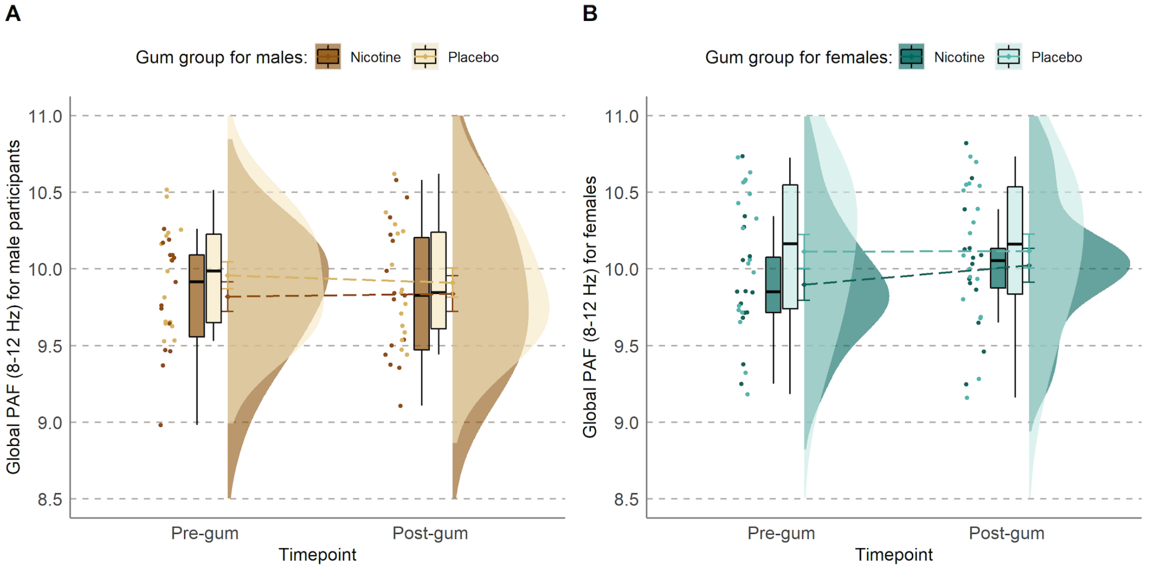

Figure 4: Global peak alpha frequency (PAF; 8–12 Hz) for all participants pre- and post-gum, split by nicotine (darker) and placebo (lighter) gum groups for A) males in brown and B) females in teal.

***Influence of sex in the effect of nicotine on PAF may be due to weight differences***

As nicotine absorption is impacted by weight, the effects of sex may be explained by significant differences in weight between female (M = 59.50 kg) and male (M = 77.39 kg) participants, for which there was extremely strong evidence for a difference from Bayesian analysis, $t(60) = -7.03, p < .001$, 95% CI: [-22.99, -12.80], $BF_{10}$ = 3714724.

When assessing the effects of group and sex on absolute change in global PAF (8–12 Hz) using a linear mixed regression model, there were significant effects of group ($b = -0.09514, p = .01$) and sex ($b = -0.078, p = .034$) on change in PAF speed. However, when weight was controlled for, there was no effect of weight ($b = -0.0015, p = .45$) or sex ($b = -0.052, p = .29$), but the effect of group remained significant (b = -0.096, p = .01).

In summary, within males, nicotine did not significantly increase global PAF compared to placebo, while within females, nicotine did significantly increase global PAF compared to placebo. However, the effect of sex was no longer present when weight was controlled for.

#### 3.5 Mediation global narrow

##### ***PHP***

Table 8: Overview of mediation model (difference score, 2W-LCS) using global PAF (9–11 Hz) and PHP. Covariate/confounding variables are sex (male/female), depressive symptoms, anxiety symptoms, perceived stress, sleep quality, and recent pain. $\chi^2(24) = 28.65, p = 0.23$, CFI = 0.98, RMSEA = 0.057, SRMR = 0.08.

| lhs | op | rhs | label | est | se | z | pvalue | ci.lower | ci.upper |
|---|---|---|---|---|---|---|---|---|---|
| deltam | =~ | m2 | | 1.000 | 0.000 | NA | NA | 1.000 | 1.000 |
| deltam | ~~ | deltam | | 0.006 | 0.001 | 4.955 | 0.000 | 0.003 | 0.007 |
| deltam | ~1 | | | 0.030 | 0.061 | 0.489 | 0.625 | -0.090 | 0.147 |
| m2 | ~ | m1 | | 1.000 | 0.000 | NA | NA | 1.000 | 1.000 |
| m2 | ~~ | m1 | | 0.000 | 0.000 | NA | NA | 0.000 | 0.000 |
| m2 | ~~ | m2 | | 0.000 | 0.000 | NA | NA | 0.000 | 0.000 |
| m2 | ~1 | | | 0.000 | 0.000 | NA | NA | 0.000 | 0.000 |
| m1 | ~1 | | | 10.000 | 0.026 | 384.968 | 0.000 | 9.948 | 10.047 |
| deltay | =~ | y2 | | 1.000 | 0.000 | NA | NA | 1.000 | 1.000 |
| deltay | ~~ | deltay | | 0.604 | 0.146 | 4.141 | 0.000 | 0.263 | 0.807 |
| deltay | ~1 | | | 1.520 | 0.813 | 1.869 | 0.062 | -0.185 | 2.955 |
| y2 | ~ | y1 | | 1.000 | 0.000 | NA | NA | 1.000 | 1.000 |
| y2 | ~~ | y1 | | 0.000 | 0.000 | NA | NA | 0.000 | 0.000 |
| y2 | ~~ | y2 | | 0.000 | 0.000 | NA | NA | 0.000 | 0.000 |
| y2 | ~1 | | | 0.000 | 0.000 | NA | NA | 0.000 | 0.000 |
| y1 | ~1 | | | 5.655 | 0.306 | 18.453 | 0.000 | 5.061 | 6.250 |
| m1 | ~~ | y1 | | -0.058 | 0.070 | -0.832 | 0.406 | -0.199 | 0.077 |
| m1 | ~~ | x | | -0.026 | 0.013 | -1.940 | 0.052 | -0.052 | 0.002 |
| y1 | ~~ | x | | 0.073 | 0.160 | 0.457 | 0.648 | -0.219 | 0.415 |
| m1 | ~~ | cov1 | | -0.022 | 0.013 | -1.652 | 0.098 | -0.049 | 0.004 |
| y1 | ~~ | cov1 | | -0.343 | 0.148 | -2.311 | 0.021 | -0.641 | -0.016 |
| m1 | ~~ | cov2 | | -0.004 | 0.021 | -0.191 | 0.848 | -0.051 | 0.038 |
| y1 | ~~ | cov2 | | -0.157 | 0.284 | -0.551 | 0.582 | -0.797 | 0.308 |
| cov2 | ~~ | cov3 | | 0.547 | 0.194 | 2.821 | 0.005 | 0.188 | 0.944 |
| m1 | ~~ | cov3 | | -0.024 | 0.020 | -1.189 | 0.235 | -0.063 | 0.017 |
| y1 | ~~ | cov3 | | 0.260 | 0.297 | 0.876 | 0.381 | -0.429 | 0.755 |
| m1 | ~~ | cov4 | | -0.043 | 0.111 | -0.390 | 0.696 | -0.269 | 0.163 |
| y1 | ~~ | cov4 | | -0.847 | 1.388 | -0.610 | 0.542 | -3.584 | 1.800 |
| m1 | ~~ | cov5 | | 0.029 | 0.042 | 0.690 | 0.490 | -0.061 | 0.110 |

| lhs | op | rhs | label | est | se | z | pvalue | ci.lower | ci.upper |
|---|---|---|---|---|---|---|---|---|---|
| y1 | ~~ | cov5 | | -0.326 | 0.436 | -0.746 | 0.456 | -1.108 | 0.645 |
| m1 | ~~ | cov6 | | -0.032 | 0.021 | -1.498 | 0.134 | -0.076 | 0.008 |
| y1 | ~~ | cov6 | | -0.015 | 0.217 | -0.070 | 0.944 | -0.457 | 0.405 |
| cov1 | ~~ | cov1 | | 0.249 | 0.008 | 30.382 | 0.000 | 0.222 | 0.250 |
| cov2 | ~~ | cov2 | | 0.916 | 0.276 | 3.316 | 0.001 | 0.403 | 1.489 |
| cov3 | ~~ | cov3 | | 1.116 | 0.462 | 2.417 | 0.016 | 0.473 | 2.110 |
| cov4 | ~~ | cov4 | | 14.612 | 2.401 | 6.085 | 0.000 | 9.560 | 19.181 |
| cov5 | ~~ | cov5 | | 3.183 | 0.655 | 4.858 | 0.000 | 1.903 | 4.415 |
| cov6 | ~~ | cov6 | | 0.616 | 0.087 | 7.101 | 0.000 | 0.450 | 0.781 |
| deltam | ~ | x | am2x | 0.029 | 0.022 | 1.352 | 0.176 | -0.015 | 0.068 |
| deltam | ~ | cov1 | | -0.039 | 0.022 | -1.771 | 0.077 | -0.087 | 0.002 |
| deltam | ~ | cov2 | | -0.012 | 0.015 | -0.771 | 0.441 | -0.035 | 0.027 |
| deltam | ~ | cov3 | | 0.023 | 0.015 | 1.599 | 0.110 | -0.013 | 0.051 |
| deltam | ~ | cov4 | | -0.002 | 0.003 | -0.688 | 0.491 | -0.009 | 0.004 |
| deltam | ~ | cov5 | | -0.004 | 0.007 | -0.585 | 0.559 | -0.018 | 0.010 |
| deltam | ~ | cov6 | | 0.022 | 0.013 | 1.675 | 0.094 | -0.006 | 0.045 |
| deltay | ~ | x | cy2x | -0.616 | 0.274 | -2.246 | 0.025 | -1.191 | -0.133 |
| deltay | ~ | deltam | by2m2 | -1.086 | 1.472 | -0.738 | 0.461 | -3.810 | 2.070 |
| deltay | ~ | cov1 | | 0.079 | 0.271 | 0.291 | 0.771 | -0.436 | 0.634 |
| deltay | ~ | cov2 | | 0.272 | 0.149 | 1.825 | 0.068 | -0.041 | 0.549 |
| deltay | ~ | cov3 | | -0.028 | 0.145 | -0.192 | 0.848 | -0.409 | 0.172 |
| deltay | ~ | cov4 | | -0.099 | 0.048 | -2.062 | 0.039 | -0.185 | -0.001 |
| deltay | ~ | cov5 | | -0.017 | 0.070 | -0.248 | 0.804 | -0.149 | 0.121 |
| deltay | ~ | cov6 | | -0.144 | 0.133 | -1.084 | 0.278 | -0.408 | 0.137 |
| m1 | ~~ | m1 | | 0.041 | 0.007 | 5.944 | 0.000 | 0.029 | 0.057 |
| y1 | ~~ | y1 | | 5.255 | 0.775 | 6.781 | 0.000 | 3.870 | 7.023 |
| x | ~~ | x | | 0.249 | 0.007 | 34.448 | 0.000 | 0.222 | 0.250 |
| x | ~1 | | | 0.467 | 0.064 | 7.279 | 0.000 | 0.334 | 0.583 |
| cov1 | ~1 | | | 0.467 | 0.064 | 7.267 | 0.000 | 0.333 | 0.583 |
| cov2 | ~1 | | | 0.517 | 0.121 | 4.267 | 0.000 | 0.300 | 0.767 |
| cov3 | ~1 | | | 0.683 | 0.139 | 4.912 | 0.000 | 0.433 | 0.967 |
| cov4 | ~1 | | | 9.767 | 0.497 | 19.640 | 0.000 | 8.800 | 10.766 |
| cov5 | ~1 | | | 6.983 | 0.232 | 30.110 | 0.000 | 6.533 | 7.433 |
| cov6 | ~1 | | | 0.983 | 0.102 | 9.628 | 0.000 | 0.783 | 1.183 |
| indirect | := | am2x*by2m2 | indirect | -0.032 | 0.056 | -0.562 | 0.574 | -0.165 | 0.082 |
| direct | := | cy2x | direct | -0.616 | 0.274 | -2.245 | 0.025 | -1.191 | -0.133 |

| lhs | op | rhs | label | est | se | z | pvalue | ci.lower | ci.upper |
|---|---|---|---|---|---|---|---|---|---|
| total | := | cy2x+ (am2x*by2m2) | total | -0.648 | 0.284 | -2.276 | 0.023 | -1.208 | -0.154 |

##### *CPA*

Table 9: Overview of mediation model (difference score, 2W-LCS) using global PAF (9–11 Hz) and CPA. Covariate/confounding variables are sex (male/female), depressive symptoms, anxiety symptoms, perceived stress, sleep quality, and recent pain. $\chi^2(24) = 31.13, p = 0.15$, CFI = 0.96, RMSEA = 0.069, SRMR = 0.085.

| lhs | op | rhs | label | est | se | z | pvalue | ci.lower | ci.upper |
|---|---|---|---|---|---|---|---|---|---|
| deltam | =~ | m2 | | 1.000 | 0.000 | NA | NA | 1.000 | 1.000 |
| deltam | ~~ | deltam | | 0.006 | 0.001 | 5.457 | 0.000 | 0.003 | 0.007 |
| deltam | ~1 | | | 0.028 | 0.063 | 0.441 | 0.659 | -0.092 | 0.152 |
| m2 | ~ | m1 | | 1.000 | 0.000 | NA | NA | 1.000 | 1.000 |
| m2 | ~~ | m1 | | 0.000 | 0.000 | NA | NA | 0.000 | 0.000 |
| m2 | ~~ | m2 | | 0.000 | 0.000 | NA | NA | 0.000 | 0.000 |
| m2 | ~1 | | | 0.000 | 0.000 | NA | NA | 0.000 | 0.000 |
| m1 | ~1 | | | 10.000 | 0.026 | 382.686 | 0.000 | 9.947 | 10.050 |
| deltay | =~ | y2 | | 1.000 | 0.000 | NA | NA | 1.000 | 1.000 |
| deltay | ~~ | deltay | | 2.201 | 0.394 | 5.580 | 0.000 | 1.045 | 2.594 |
| deltay | ~1 | | | -1.287 | 1.383 | -0.930 | 0.352 | -3.984 | 1.342 |
| y2 | ~ | y1 | | 1.000 | 0.000 | NA | NA | 1.000 | 1.000 |
| y2 | ~~ | y1 | | 0.000 | 0.000 | NA | NA | 0.000 | 0.000 |
| y2 | ~~ | y2 | | 0.000 | 0.000 | NA | NA | 0.000 | 0.000 |
| y2 | ~1 | | | 0.000 | 0.000 | NA | NA | 0.000 | 0.000 |
| y1 | ~1 | | | 4.580 | 0.193 | 23.680 | 0.000 | 4.202 | 4.947 |
| m1 | ~~ | y1 | | -0.006 | 0.042 | -0.131 | 0.896 | -0.083 | 0.084 |
| m1 | ~~ | x | | -0.023 | 0.013 | -1.823 | 0.068 | -0.048 | 0.002 |
| y1 | ~~ | x | | 0.159 | 0.098 | 1.625 | 0.104 | -0.041 | 0.337 |
| m1 | ~~ | cov1 | | -0.022 | 0.013 | -1.676 | 0.094 | -0.050 | 0.003 |
| y1 | ~~ | cov1 | | -0.124 | 0.095 | -1.301 | 0.193 | -0.309 | 0.061 |
| m1 | ~~ | cov2 | | -0.010 | 0.021 | -0.465 | 0.642 | -0.049 | 0.032 |
| y1 | ~~ | cov2 | | -0.271 | 0.196 | -1.384 | 0.166 | -0.703 | 0.086 |
| cov2 | ~~ | cov3 | | 0.566 | 0.187 | 3.035 | 0.002 | 0.205 | 0.944 |
| m1 | ~~ | cov3 | | -0.029 | 0.020 | -1.429 | 0.153 | -0.069 | 0.012 |
| y1 | ~~ | cov3 | | -0.115 | 0.243 | -0.475 | 0.635 | -0.658 | 0.295 |
| m1 | ~~ | cov4 | | -0.036 | 0.108 | -0.337 | 0.736 | -0.265 | 0.169 |
| y1 | ~~ | cov4 | | -0.226 | 0.731 | -0.309 | 0.757 | -1.645 | 1.206 |
| m1 | ~~ | cov5 | | 0.026 | 0.042 | 0.625 | 0.532 | -0.061 | 0.110 |
| y1 | ~~ | cov5 | | -0.373 | 0.388 | -0.962 | 0.336 | -1.144 | 0.303 |
| m1 | ~~ | cov6 | | -0.033 | 0.021 | -1.543 | 0.123 | -0.077 | 0.007 |

| lhs | op | rhs | label | est | se | z | pvalue | ci.lower | ci.upper |
|---|---|---|---|---|---|---|---|---|---|
| y1 | ~~ | cov6 | | -0.178 | 0.142 | -1.250 | 0.211 | -0.468 | 0.097 |
| cov1 | ~~ | cov1 | | 0.250 | 0.006 | 44.155 | 0.000 | 0.229 | 0.250 |
| cov2 | ~~ | cov2 | | 0.926 | 0.263 | 3.523 | 0.000 | 0.432 | 1.489 |
| cov3 | ~~ | cov3 | | 1.116 | 0.434 | 2.571 | 0.010 | 0.476 | 2.046 |
| cov4 | ~~ | cov4 | | 14.307 | 2.304 | 6.210 | 0.000 | 9.642 | 18.652 |
| cov5 | ~~ | cov5 | | 3.096 | 0.645 | 4.796 | 0.000 | 1.939 | 4.450 |
| cov6 | ~~ | cov6 | | 0.627 | 0.085 | 7.399 | 0.000 | 0.467 | 0.805 |
| deltam | ~ | x | am2x | 0.037 | 0.021 | 1.743 | 0.081 | -0.006 | 0.077 |
| deltam | ~ | cov1 | | -0.044 | 0.021 | -2.091 | 0.037 | -0.088 | -0.002 |
| deltam | ~ | cov2 | | -0.015 | 0.015 | -1.039 | 0.299 | -0.040 | 0.019 |
| deltam | ~ | cov3 | | 0.021 | 0.014 | 1.529 | 0.126 | -0.010 | 0.048 |
| deltam | ~ | cov4 | | -0.002 | 0.003 | -0.706 | 0.480 | -0.009 | 0.004 |
| deltam | ~ | cov5 | | -0.004 | 0.007 | -0.591 | 0.554 | -0.017 | 0.011 |
| deltam | ~ | cov6 | | 0.024 | 0.013 | 1.939 | 0.052 | -0.002 | 0.048 |
| deltay | ~ | x | cy2x | -0.192 | 0.412 | -0.467 | 0.640 | -1.054 | 0.566 |
| deltay | ~ | deltam | by2m2 | -2.424 | 2.713 | -0.894 | 0.372 | -8.062 | 2.579 |
| deltay | ~ | cov1 | | 0.719 | 0.426 | 1.686 | 0.092 | -0.112 | 1.535 |
| deltay | ~ | cov2 | | 0.190 | 0.347 | 0.548 | 0.584 | -0.541 | 0.824 |
| deltay | ~ | cov3 | | -0.193 | 0.393 | -0.490 | 0.624 | -1.103 | 0.328 |
| deltay | ~ | cov4 | | -0.056 | 0.062 | -0.901 | 0.367 | -0.175 | 0.063 |
| deltay | ~ | cov5 | | 0.090 | 0.144 | 0.626 | 0.532 | -0.179 | 0.384 |
| deltay | ~ | cov6 | | 0.114 | 0.241 | 0.472 | 0.637 | -0.360 | 0.593 |
| m1 | ~~ | m1 | | 0.040 | 0.007 | 5.786 | 0.000 | 0.029 | 0.055 |
| y1 | ~~ | y1 | | 2.288 | 0.413 | 5.537 | 0.000 | 1.616 | 3.194 |
| x | ~~ | x | | 0.249 | 0.007 | 36.005 | 0.000 | 0.224 | 0.250 |
| x | ~1 | | | 0.468 | 0.063 | 7.392 | 0.000 | 0.339 | 0.581 |
| cov1 | ~1 | | | 0.484 | 0.063 | 7.707 | 0.000 | 0.355 | 0.613 |
| cov2 | ~1 | | | 0.532 | 0.121 | 4.409 | 0.000 | 0.306 | 0.774 |
| cov3 | ~1 | | | 0.694 | 0.134 | 5.185 | 0.000 | 0.452 | 0.968 |
| cov4 | ~1 | | | 9.823 | 0.486 | 20.196 | 0.000 | 8.839 | 10.758 |
| cov5 | ~1 | | | 6.968 | 0.223 | 31.179 | 0.000 | 6.516 | 7.403 |
| cov6 | ~1 | | | 0.952 | 0.103 | 9.280 | 0.000 | 0.758 | 1.161 |
| indirect | := | am2x*by2m2 | indirect | -0.090 | 0.123 | -0.737 | 0.461 | -0.390 | 0.113 |
| direct | := | cy2x | direct | -0.192 | 0.412 | -0.467 | 0.641 | -1.054 | 0.566 |
| total | := | cy2x+ (am2x*by2m2) | total | -0.283 | 0.418 | -0.676 | 0.499 | -1.128 | 0.468 |

#### 3.6 Mediation SM wide

##### *PHP*

Table 10: Overview of mediation model using sensorimotor PAF (8–12 Hz) and PHP. Covariate/confounding variables are sex (male/female), depressive symptoms, anxiety symptoms, perceived stress, sleep quality, and recent pain. $\chi^2(24) = 24.74, p = 0.42$, CFI = 0.997, RMSEA = 0.023, SRMR = 0.078.

| lhs | op | rhs | label | est | se | z | pvalue | ci.lower | ci.upper |
|---|---|---|---|---|---|---|---|---|---|
| deltam | =~ | m2 | | 1.000 | 0.000 | NA | NA | 1.000 | 1.000 |
| deltam | ~~ | deltam | | 0.021 | 0.004 | 5.757 | 0.000 | 0.011 | 0.025 |
| deltam | ~1 | | | 0.173 | 0.140 | 1.231 | 0.218 | -0.105 | 0.433 |
| m2 | ~ | m1 | | 1.000 | 0.000 | NA | NA | 1.000 | 1.000 |
| m2 | ~~ | m1 | | 0.000 | 0.000 | NA | NA | 0.000 | 0.000 |
| m2 | ~~ | m2 | | 0.000 | 0.000 | NA | NA | 0.000 | 0.000 |
| m2 | ~1 | | | 0.000 | 0.000 | NA | NA | 0.000 | 0.000 |
| m1 | ~1 | | | 9.900 | 0.048 | 207.936 | 0.000 | 9.805 | 9.991 |
| deltay | =~ | y2 | | 1.000 | 0.000 | NA | NA | 1.000 | 1.000 |
| deltay | ~~ | deltay | | 0.609 | 0.150 | 4.071 | 0.000 | 0.261 | 0.826 |
| deltay | ~1 | | | 1.537 | 0.839 | 1.831 | 0.067 | -0.231 | 3.065 |
| y2 | ~ | y1 | | 1.000 | 0.000 | NA | NA | 1.000 | 1.000 |
| y2 | ~~ | y1 | | 0.000 | 0.000 | NA | NA | 0.000 | 0.000 |
| y2 | ~~ | y2 | | 0.000 | 0.000 | NA | NA | 0.000 | 0.000 |
| y2 | ~1 | | | 0.000 | 0.000 | NA | NA | 0.000 | 0.000 |
| y1 | ~1 | | | 5.655 | 0.306 | 18.453 | 0.000 | 5.061 | 6.250 |
| m1 | ~~ | y1 | | -0.130 | 0.124 | -1.048 | 0.295 | -0.367 | 0.126 |
| m1 | ~~ | x | | -0.041 | 0.025 | -1.619 | 0.105 | -0.088 | 0.010 |
| y1 | ~~ | x | | 0.073 | 0.160 | 0.457 | 0.648 | -0.219 | 0.415 |
| m1 | ~~ | cov1 | | -0.015 | 0.026 | -0.575 | 0.565 | -0.070 | 0.034 |
| y1 | ~~ | cov1 | | -0.343 | 0.148 | -2.311 | 0.021 | -0.641 | -0.016 |
| m1 | ~~ | cov2 | | -0.008 | 0.046 | -0.166 | 0.869 | -0.103 | 0.084 |
| y1 | ~~ | cov2 | | -0.157 | 0.284 | -0.551 | 0.582 | -0.797 | 0.308 |
| cov2 | ~~ | cov3 | | 0.547 | 0.194 | 2.821 | 0.005 | 0.188 | 0.944 |
| m1 | ~~ | cov3 | | -0.031 | 0.043 | -0.714 | 0.475 | -0.114 | 0.056 |
| y1 | ~~ | cov3 | | 0.260 | 0.297 | 0.876 | 0.381 | -0.429 | 0.755 |
| m1 | ~~ | cov4 | | 0.070 | 0.215 | 0.327 | 0.744 | -0.360 | 0.478 |
| y1 | ~~ | cov4 | | -0.847 | 1.388 | -0.610 | 0.542 | -3.584 | 1.800 |
| m1 | ~~ | cov5 | | 0.023 | 0.090 | 0.251 | 0.802 | -0.175 | 0.191 |

| lhs | op | rhs | label | est | se | z | pvalue | ci.lower | ci.upper |
|---|---|---|---|---|---|---|---|---|---|
| y1 | ~~ | cov5 | | -0.326 | 0.436 | -0.746 | 0.456 | -1.108 | 0.645 |
| m1 | ~~ | cov6 | | -0.049 | 0.041 | -1.182 | 0.237 | -0.128 | 0.031 |
| y1 | ~~ | cov6 | | -0.015 | 0.217 | -0.070 | 0.944 | -0.457 | 0.405 |
| cov1 | ~~ | cov1 | | 0.249 | 0.008 | 30.382 | 0.000 | 0.222 | 0.250 |
| cov2 | ~~ | cov2 | | 0.916 | 0.276 | 3.316 | 0.001 | 0.403 | 1.489 |
| cov3 | ~~ | cov3 | | 1.116 | 0.462 | 2.417 | 0.016 | 0.473 | 2.110 |
| cov4 | ~~ | cov4 | | 14.612 | 2.401 | 6.085 | 0.000 | 9.560 | 19.181 |
| cov5 | ~~ | cov5 | | 3.183 | 0.655 | 4.858 | 0.000 | 1.903 | 4.415 |
| cov6 | ~~ | cov6 | | 0.616 | 0.087 | 7.101 | 0.000 | 0.450 | 0.781 |
| deltam | ~ | x | am2x | 0.061 | 0.038 | 1.604 | 0.109 | -0.008 | 0.132 |
| deltam | ~ | cov1 | | -0.083 | 0.041 | -2.020 | 0.043 | -0.171 | -0.010 |
| deltam | ~ | cov2 | | -0.019 | 0.031 | -0.614 | 0.539 | -0.069 | 0.054 |
| deltam | ~ | cov3 | | 0.012 | 0.027 | 0.457 | 0.647 | -0.043 | 0.070 |
| deltam | ~ | cov4 | | -0.009 | 0.006 | -1.438 | 0.150 | -0.022 | 0.003 |
| deltam | ~ | cov5 | | -0.010 | 0.016 | -0.653 | 0.514 | -0.039 | 0.025 |
| deltam | ~ | cov6 | | 0.024 | 0.025 | 0.958 | 0.338 | -0.027 | 0.071 |
| deltay | ~ | x | cy2x | -0.630 | 0.285 | -2.212 | 0.027 | -1.220 | -0.150 |
| deltay | ~ | deltam | by2m2 | -0.285 | 0.758 | -0.376 | 0.707 | -1.689 | 1.248 |
| deltay | ~ | cov1 | | 0.098 | 0.282 | 0.347 | 0.729 | -0.415 | 0.682 |
| deltay | ~ | cov2 | | 0.280 | 0.154 | 1.812 | 0.070 | -0.048 | 0.563 |
| deltay | ~ | cov3 | | -0.050 | 0.138 | -0.362 | 0.718 | -0.419 | 0.126 |
| deltay | ~ | cov4 | | -0.099 | 0.048 | -2.044 | 0.041 | -0.189 | 0.001 |
| deltay | ~ | cov5 | | -0.016 | 0.070 | -0.225 | 0.822 | -0.154 | 0.128 |
| deltay | ~ | cov6 | | -0.161 | 0.127 | -1.266 | 0.205 | -0.412 | 0.100 |
| m1 | ~~ | m1 | | 0.141 | 0.024 | 5.949 | 0.000 | 0.099 | 0.191 |
| y1 | ~~ | y1 | | 5.255 | 0.775 | 6.781 | 0.000 | 3.870 | 7.022 |
| x | ~~ | x | | 0.249 | 0.007 | 34.448 | 0.000 | 0.222 | 0.250 |
| x | ~1 | | | 0.467 | 0.064 | 7.279 | 0.000 | 0.334 | 0.583 |
| cov1 | ~1 | | | 0.467 | 0.064 | 7.267 | 0.000 | 0.333 | 0.583 |
| cov2 | ~1 | | | 0.517 | 0.121 | 4.267 | 0.000 | 0.300 | 0.767 |
| cov3 | ~1 | | | 0.683 | 0.139 | 4.912 | 0.000 | 0.433 | 0.967 |
| cov4 | ~1 | | | 9.767 | 0.497 | 19.640 | 0.000 | 8.800 | 10.766 |
| cov5 | ~1 | | | 6.983 | 0.232 | 30.110 | 0.000 | 6.533 | 7.433 |
| cov6 | ~1 | | | 0.983 | 0.102 | 9.628 | 0.000 | 0.783 | 1.183 |
| indirect | := | am2x*by2m2 | indirect | -0.017 | 0.052 | -0.334 | 0.738 | -0.100 | 0.125 |
| direct | := | cy2x | direct | -0.630 | 0.285 | -2.211 | 0.027 | -1.220 | -0.150 |

| lhs | op | rhs | label | est | se | z | pvalue | ci.lower | ci.upper |
|---|---|---|---|---|---|---|---|---|---|
| total | := | cy2x+ (am2x*by2m2) | total | -0.648 | 0.284 | -2.276 | 0.023 | -1.208 | -0.154 |

##### *CPA*

Table 11: Overview of mediation model using sensorimotor PAF (8–12 Hz) and CPA. Covariate/confounding variables are sex (male/female), depressive symptoms, anxiety symptoms, perceived stress, sleep quality, and recent pain. $\chi^2(24) = 28.54, p = 0.24$, CFI = 0.97, RMSEA = 0.055, SRMR = 0.084.

| lhs | op | rhs | label | est | se | z | pvalue | ci.lower | ci.upper |
|---|---|---|---|---|---|---|---|---|---|
| deltam | =~ | m2 | | 1.000 | 0.000 | NA | NA | 1.000 | 1.000 |
| deltam | ~~ | deltam | | 0.021 | 0.003 | 6.246 | 0.000 | 0.012 | 0.025 |
| deltam | ~1 | | | 0.174 | 0.140 | 1.248 | 0.212 | -0.108 | 0.450 |
| m2 | ~ | m1 | | 1.000 | 0.000 | NA | NA | 1.000 | 1.000 |
| m2 | ~~ | m1 | | 0.000 | 0.000 | NA | NA | 0.000 | 0.000 |
| m2 | ~~ | m2 | | 0.000 | 0.000 | NA | NA | 0.000 | 0.000 |
| m2 | ~1 | | | 0.000 | 0.000 | NA | NA | 0.000 | 0.000 |
| m1 | ~1 | | | 9.897 | 0.049 | 202.366 | 0.000 | 9.801 | 9.986 |
| deltay | =~ | y2 | | 1.000 | 0.000 | NA | NA | 1.000 | 1.000 |
| deltay | ~~ | deltay | | 2.211 | 0.401 | 5.508 | 0.000 | 1.025 | 2.602 |
| deltay | ~1 | | | -1.163 | 1.410 | -0.825 | 0.409 | -3.962 | 1.654 |
| y2 | ~ | y1 | | 1.000 | 0.000 | NA | NA | 1.000 | 1.000 |
| y2 | ~~ | y1 | | 0.000 | 0.000 | NA | NA | 0.000 | 0.000 |
| y2 | ~~ | y2 | | 0.000 | 0.000 | NA | NA | 0.000 | 0.000 |
| y2 | ~1 | | | 0.000 | 0.000 | NA | NA | 0.000 | 0.000 |
| y1 | ~1 | | | 4.580 | 0.193 | 23.680 | 0.000 | 4.202 | 4.947 |
| m1 | ~~ | y1 | | -0.042 | 0.077 | -0.542 | 0.588 | -0.189 | 0.109 |
| m1 | ~~ | x | | -0.037 | 0.023 | -1.578 | 0.114 | -0.080 | 0.010 |
| y1 | ~~ | x | | 0.159 | 0.098 | 1.625 | 0.104 | -0.041 | 0.337 |
| m1 | ~~ | cov1 | | -0.018 | 0.026 | -0.721 | 0.471 | -0.071 | 0.030 |
| y1 | ~~ | cov1 | | -0.124 | 0.095 | -1.301 | 0.193 | -0.309 | 0.061 |
| m1 | ~~ | cov2 | | -0.018 | 0.046 | -0.391 | 0.696 | -0.103 | 0.081 |
| y1 | ~~ | cov2 | | -0.271 | 0.196 | -1.385 | 0.166 | -0.703 | 0.086 |
| cov2 | ~~ | cov3 | | 0.566 | 0.187 | 3.035 | 0.002 | 0.205 | 0.944 |
| m1 | ~~ | cov3 | | -0.040 | 0.044 | -0.909 | 0.363 | -0.140 | 0.047 |
| y1 | ~~ | cov3 | | -0.115 | 0.243 | -0.475 | 0.635 | -0.658 | 0.295 |
| m1 | ~~ | cov4 | | 0.064 | 0.213 | 0.298 | 0.765 | -0.355 | 0.460 |
| y1 | ~~ | cov4 | | -0.226 | 0.731 | -0.309 | 0.757 | -1.645 | 1.206 |
| m1 | ~~ | cov5 | | 0.024 | 0.087 | 0.278 | 0.781 | -0.160 | 0.196 |
| y1 | ~~ | cov5 | | -0.373 | 0.388 | -0.962 | 0.336 | -1.144 | 0.303 |
| m1 | ~~ | cov6 | | -0.044 | 0.040 | -1.091 | 0.275 | -0.127 | 0.035 |

| lhs | op | rhs | label | est | se | z | pvalue | ci.lower | ci.upper |
|---|---|---|---|---|---|---|---|---|---|
| y1 | ~~ | cov6 | | -0.178 | 0.142 | -1.250 | 0.211 | -0.468 | 0.097 |
| cov1 | ~~ | cov1 | | 0.250 | 0.006 | 44.155 | 0.000 | 0.229 | 0.250 |
| cov2 | ~~ | cov2 | | 0.926 | 0.263 | 3.523 | 0.000 | 0.432 | 1.489 |
| cov3 | ~~ | cov3 | | 1.116 | 0.434 | 2.571 | 0.010 | 0.476 | 2.046 |
| cov4 | ~~ | cov4 | | 14.307 | 2.304 | 6.210 | 0.000 | 9.642 | 18.652 |
| cov5 | ~~ | cov5 | | 3.096 | 0.645 | 4.796 | 0.000 | 1.939 | 4.450 |
| cov6 | ~~ | cov6 | | 0.627 | 0.085 | 7.399 | 0.000 | 0.467 | 0.805 |
| deltam | ~ | x | am2x | 0.071 | 0.037 | 1.912 | 0.056 | -0.004 | 0.142 |
| deltam | ~ | cov1 | | -0.082 | 0.038 | -2.181 | 0.029 | -0.165 | -0.013 |
| deltam | ~ | cov2 | | -0.022 | 0.029 | -0.755 | 0.450 | -0.068 | 0.049 |
| deltam | ~ | cov3 | | 0.010 | 0.025 | 0.379 | 0.704 | -0.040 | 0.064 |
| deltam | ~ | cov4 | | -0.008 | 0.006 | -1.353 | 0.176 | -0.021 | 0.004 |
| deltam | ~ | cov5 | | -0.011 | 0.015 | -0.749 | 0.454 | -0.038 | 0.022 |
| deltam | ~ | cov6 | | 0.023 | 0.023 | 0.992 | 0.321 | -0.027 | 0.070 |
| deltay | ~ | x | cy2x | -0.205 | 0.411 | -0.499 | 0.618 | -1.100 | 0.549 |
| deltay | ~ | deltam | by2m2 | -1.098 | 1.657 | -0.663 | 0.508 | -4.661 | 1.745 |
| deltay | ~ | cov1 | | 0.735 | 0.442 | 1.664 | 0.096 | -0.143 | 1.547 |
| deltay | ~ | cov2 | | 0.203 | 0.345 | 0.588 | 0.556 | -0.541 | 0.830 |
| deltay | ~ | cov3 | | -0.233 | 0.386 | -0.605 | 0.545 | -1.162 | 0.290 |
| deltay | ~ | cov4 | | -0.060 | 0.067 | -0.892 | 0.372 | -0.187 | 0.069 |
| deltay | ~ | cov5 | | 0.088 | 0.145 | 0.605 | 0.545 | -0.175 | 0.370 |
| deltay | ~ | cov6 | | 0.080 | 0.242 | 0.331 | 0.740 | -0.391 | 0.569 |
| m1 | ~~ | m1 | | 0.137 | 0.023 | 5.967 | 0.000 | 0.097 | 0.184 |
| y1 | ~~ | y1 | | 2.288 | 0.413 | 5.537 | 0.000 | 1.616 | 3.194 |
| x | ~~ | x | | 0.249 | 0.007 | 36.005 | 0.000 | 0.224 | 0.250 |
| x | ~1 | | | 0.468 | 0.063 | 7.392 | 0.000 | 0.339 | 0.581 |
| cov1 | ~1 | | | 0.484 | 0.063 | 7.707 | 0.000 | 0.355 | 0.613 |
| cov2 | ~1 | | | 0.532 | 0.121 | 4.409 | 0.000 | 0.306 | 0.774 |
| cov3 | ~1 | | | 0.694 | 0.134 | 5.185 | 0.000 | 0.452 | 0.968 |
| cov4 | ~1 | | | 9.823 | 0.486 | 20.196 | 0.000 | 8.839 | 10.758 |
| cov5 | ~1 | | | 6.968 | 0.223 | 31.179 | 0.000 | 6.516 | 7.403 |
| cov6 | ~1 | | | 0.952 | 0.103 | 9.280 | 0.000 | 0.758 | 1.161 |
| indirect | := | am2x*by2m2 | indirect | -0.078 | 0.135 | -0.577 | 0.564 | -0.440 | 0.128 |
| direct | := | cy2x | direct | -0.205 | 0.411 | -0.498 | 0.618 | -1.100 | 0.549 |
| total | := | cy2x+ (am2x*by2m2) | total | -0.283 | 0.418 | -0.676 | 0.499 | -1.128 | 0.468 |

#### 3.7 Mediation SM narrow

##### ***PHP***

Table 12: Overview of mediation model (difference score, 2W-LCS) using sensorimotor PAF (9–11 Hz) and PHP. Covariate/confounding variables are sex (male/female), depressive symptoms, anxiety symptoms, perceived stress, sleep quality, and recent pain. $\chi^2(24) = 26.50, p = 0.33$, CFI = 0.99, RMSEA = 0.042, SRMR = 0.080.

| lhs | op | rhs | label | est | se | z | pvalue | ci.lower | ci.upper |
|---|---|---|---|---|---|---|---|---|---|
| deltam | =~ | m2 | | 1.000 | 0.000 | NA | NA | 1.000 | 1.000 |
| deltam | ~~ | deltam | | 0.008 | 0.001 | 5.505 | 0.000 | 0.004 | 0.010 |
| deltam | ~1 | | | 0.060 | 0.082 | 0.729 | 0.466 | -0.104 | 0.214 |
| m2 | ~ | m1 | | 1.000 | 0.000 | NA | NA | 1.000 | 1.000 |
| m2 | ~~ | m1 | | 0.000 | 0.000 | NA | NA | 0.000 | 0.000 |
| m2 | ~~ | m2 | | 0.000 | 0.000 | NA | NA | 0.000 | 0.000 |
| m2 | ~1 | | | 0.000 | 0.000 | NA | NA | 0.000 | 0.000 |
| m1 | ~1 | | | 9.971 | 0.024 | 420.912 | 0.000 | 9.927 | 10.019 |
| deltay | =~ | y2 | | 1.000 | 0.000 | NA | NA | 1.000 | 1.000 |
| deltay | ~~ | deltay | | 0.585 | 0.140 | 4.187 | 0.000 | 0.256 | 0.778 |
| deltay | ~1 | | | 1.596 | 0.823 | 1.939 | 0.053 | -0.204 | 3.040 |
| y2 | ~ | y1 | | 1.000 | 0.000 | NA | NA | 1.000 | 1.000 |
| y2 | ~~ | y1 | | 0.000 | 0.000 | NA | NA | 0.000 | 0.000 |
| y2 | ~~ | y2 | | 0.000 | 0.000 | NA | NA | 0.000 | 0.000 |
| y2 | ~1 | | | 0.000 | 0.000 | NA | NA | 0.000 | 0.000 |
| y1 | ~1 | | | 5.655 | 0.298 | 18.991 | 0.000 | 5.046 | 6.233 |
| m1 | ~~ | y1 | | -0.080 | 0.064 | -1.256 | 0.209 | -0.222 | 0.043 |
| m1 | ~~ | x | | -0.021 | 0.012 | -1.709 | 0.087 | -0.046 | 0.002 |
| y1 | ~~ | x | | 0.073 | 0.170 | 0.430 | 0.667 | -0.223 | 0.461 |
| m1 | ~~ | cov1 | | -0.014 | 0.012 | -1.165 | 0.244 | -0.037 | 0.010 |
| y1 | ~~ | cov1 | | -0.343 | 0.144 | -2.383 | 0.017 | -0.620 | -0.058 |
| m1 | ~~ | cov2 | | 0.003 | 0.020 | 0.149 | 0.882 | -0.037 | 0.042 |
| y1 | ~~ | cov2 | | -0.157 | 0.274 | -0.573 | 0.567 | -0.750 | 0.345 |
| cov2 | ~~ | cov3 | | 0.547 | 0.192 | 2.853 | 0.004 | 0.188 | 0.927 |
| m1 | ~~ | cov3 | | -0.018 | 0.019 | -0.954 | 0.340 | -0.053 | 0.022 |
| y1 | ~~ | cov3 | | 0.260 | 0.301 | 0.866 | 0.387 | -0.404 | 0.793 |
| m1 | ~~ | cov4 | | -0.033 | 0.102 | -0.325 | 0.745 | -0.236 | 0.167 |
| y1 | ~~ | cov4 | | -0.847 | 1.390 | -0.609 | 0.542 | -3.365 | 2.115 |
| m1 | ~~ | cov5 | | 0.021 | 0.039 | 0.534 | 0.593 | -0.056 | 0.099 |

| lhs | op | rhs | label | est | se | z | pvalue | ci.lower | ci.upper |
|---|---|---|---|---|---|---|---|---|---|
| y1 | ~~ | cov5 | | -0.326 | 0.455 | -0.716 | 0.474 | -1.172 | 0.610 |
| m1 | ~~ | cov6 | | -0.040 | 0.020 | -1.959 | 0.050 | -0.080 | -0.001 |
| y1 | ~~ | cov6 | | -0.015 | 0.225 | -0.067 | 0.947 | -0.455 | 0.393 |
| cov1 | ~~ | cov1 | | 0.249 | 0.007 | 35.110 | 0.000 | 0.227 | 0.250 |
| cov2 | ~~ | cov2 | | 0.916 | 0.277 | 3.311 | 0.001 | 0.428 | 1.503 |
| cov3 | ~~ | cov3 | | 1.116 | 0.457 | 2.444 | 0.015 | 0.476 | 2.083 |
| cov4 | ~~ | cov4 | | 14.612 | 2.449 | 5.967 | 0.000 | 9.877 | 19.432 |
| cov5 | ~~ | cov5 | | 3.183 | 0.651 | 4.891 | 0.000 | 1.970 | 4.503 |
| cov6 | ~~ | cov6 | | 0.616 | 0.083 | 7.413 | 0.000 | 0.449 | 0.781 |
| deltam | ~ | x | am2x | 0.022 | 0.024 | 0.901 | 0.367 | -0.029 | 0.069 |
| deltam | ~ | cov1 | | -0.042 | 0.025 | -1.661 | 0.097 | -0.099 | 0.006 |
| deltam | ~ | cov2 | | -0.002 | 0.019 | -0.094 | 0.925 | -0.033 | 0.043 |
| deltam | ~ | cov3 | | 0.011 | 0.019 | 0.575 | 0.566 | -0.020 | 0.054 |
| deltam | ~ | cov4 | | -0.004 | 0.004 | -1.139 | 0.255 | -0.012 | 0.003 |
| deltam | ~ | cov5 | | -0.006 | 0.009 | -0.695 | 0.487 | -0.022 | 0.012 |
| deltam | ~ | cov6 | | 0.034 | 0.014 | 2.387 | 0.017 | 0.006 | 0.061 |
| deltay | ~ | x | cy2x | -0.608 | 0.278 | -2.188 | 0.029 | -1.145 | -0.103 |
| deltay | ~ | deltam | by2m2 | -1.815 | 1.296 | -1.400 | 0.162 | -4.362 | 0.676 |
| deltay | ~ | cov1 | | 0.045 | 0.251 | 0.179 | 0.858 | -0.426 | 0.571 |
| deltay | ~ | cov2 | | 0.282 | 0.148 | 1.907 | 0.056 | -0.033 | 0.537 |
| deltay | ~ | cov3 | | -0.033 | 0.133 | -0.251 | 0.802 | -0.372 | 0.167 |
| deltay | ~ | cov4 | | -0.104 | 0.049 | -2.122 | 0.034 | -0.191 | -0.007 |
| deltay | ~ | cov5 | | -0.024 | 0.068 | -0.350 | 0.727 | -0.147 | 0.129 |
| deltay | ~ | cov6 | | -0.106 | 0.129 | -0.828 | 0.408 | -0.361 | 0.157 |
| m1 | ~~ | m1 | | 0.035 | 0.007 | 5.334 | 0.000 | 0.023 | 0.050 |
| y1 | ~~ | y1 | | 5.255 | 0.814 | 6.456 | 0.000 | 3.869 | 7.038 |
| x | ~~ | x | | 0.249 | 0.008 | 33.010 | 0.000 | 0.222 | 0.250 |
| x | ~1 | | | 0.467 | 0.065 | 7.185 | 0.000 | 0.333 | 0.583 |
| cov1 | ~1 | | | 0.467 | 0.064 | 7.338 | 0.000 | 0.350 | 0.583 |
| cov2 | ~1 | | | 0.517 | 0.124 | 4.173 | 0.000 | 0.300 | 0.783 |
| cov3 | ~1 | | | 0.683 | 0.130 | 5.257 | 0.000 | 0.467 | 0.950 |
| cov4 | ~1 | | | 9.767 | 0.491 | 19.898 | 0.000 | 8.800 | 10.750 |
| cov5 | ~1 | | | 6.983 | 0.230 | 30.338 | 0.000 | 6.517 | 7.417 |
| cov6 | ~1 | | | 0.983 | 0.101 | 9.714 | 0.000 | 0.784 | 1.200 |
| indirect | := | am2x*by2m2 | indirect | -0.040 | 0.056 | -0.712 | 0.476 | -0.167 | 0.067 |
| direct | := | cy2x | direct | -0.608 | 0.278 | -2.187 | 0.029 | -1.145 | -0.103 |

| lhs | op | rhs | label | est | se | z | pvalue | ci.lower | ci.upper |
|---|---|---|---|---|---|---|---|---|---|
| total | := | cy2x+ (am2x*by2m2) | total | -0.648 | 0.289 | -2.244 | 0.025 | -1.193 | -0.114 |

***CPA***

Table 13: Overview of mediation model (difference score, 2W-LCS) using sensorimotor PAF (9–11 Hz) and CPA. Covariate/confounding variables are sex (male/female), depressive symptoms, anxiety symptoms, perceived stress, sleep quality, and recent pain. $\chi^2(24) = 29.60, p = 0.20$, CFI = 0.96, RMSEA = 0.061, SRMR = 0.084.

| lhs | op | rhs | label | est | se | z | pvalue | ci.lower | ci.upper |
|---|---|---|---|---|---|---|---|---|---|
| deltam | =~ | m2 | | 1.000 | 0.000 | NA | NA | 1.000 | 1.000 |
| deltam | ~~ | deltam | | 0.008 | 0.001 | 6.039 | 0.000 | 0.005 | 0.010 |
| deltam | ~1 | | | 0.059 | 0.082 | 0.721 | 0.471 | -0.090 | 0.226 |
| m2 | ~ | m1 | | 1.000 | 0.000 | NA | NA | 1.000 | 1.000 |
| m2 | ~~ | m1 | | 0.000 | 0.000 | NA | NA | 0.000 | 0.000 |
| m2 | ~~ | m2 | | 0.000 | 0.000 | NA | NA | 0.000 | 0.000 |
| m2 | ~1 | | | 0.000 | 0.000 | NA | NA | 0.000 | 0.000 |
| m1 | ~1 | | | 9.971 | 0.024 | 410.988 | 0.000 | 9.923 | 10.018 |
| deltay | =~ | y2 | | 1.000 | 0.000 | NA | NA | 1.000 | 1.000 |
| deltay | ~~ | deltay | | 2.193 | 0.395 | 5.551 | 0.000 | 1.037 | 2.606 |
| deltay | ~1 | | | -1.221 | 1.376 | -0.887 | 0.375 | -3.899 | 1.350 |
| y2 | ~ | y1 | | 1.000 | 0.000 | NA | NA | 1.000 | 1.000 |
| y2 | ~~ | y1 | | 0.000 | 0.000 | NA | NA | 0.000 | 0.000 |
| y2 | ~~ | y2 | | 0.000 | 0.000 | NA | NA | 0.000 | 0.000 |
| y2 | ~1 | | | 0.000 | 0.000 | NA | NA | 0.000 | 0.000 |
| y1 | ~1 | | | 4.580 | 0.193 | 23.680 | 0.000 | 4.202 | 4.947 |
| m1 | ~~ | y1 | | -0.014 | 0.037 | -0.376 | 0.707 | -0.082 | 0.061 |
| m1 | ~~ | x | | -0.019 | 0.012 | -1.598 | 0.110 | -0.042 | 0.005 |
| y1 | ~~ | x | | 0.159 | 0.098 | 1.625 | 0.104 | -0.041 | 0.337 |
| m1 | ~~ | cov1 | | -0.015 | 0.012 | -1.216 | 0.224 | -0.040 | 0.009 |
| y1 | ~~ | cov1 | | -0.124 | 0.095 | -1.301 | 0.193 | -0.309 | 0.061 |
| m1 | ~~ | cov2 | | -0.001 | 0.019 | -0.075 | 0.940 | -0.037 | 0.039 |
| y1 | ~~ | cov2 | | -0.271 | 0.196 | -1.385 | 0.166 | -0.703 | 0.086 |
| cov2 | ~~ | cov3 | | 0.566 | 0.187 | 3.035 | 0.002 | 0.205 | 0.944 |
| m1 | ~~ | cov3 | | -0.022 | 0.019 | -1.167 | 0.243 | -0.055 | 0.018 |
| y1 | ~~ | cov3 | | -0.115 | 0.243 | -0.475 | 0.635 | -0.658 | 0.295 |
| m1 | ~~ | cov4 | | -0.031 | 0.100 | -0.315 | 0.753 | -0.240 | 0.153 |
| y1 | ~~ | cov4 | | -0.226 | 0.731 | -0.309 | 0.757 | -1.645 | 1.206 |
| m1 | ~~ | cov5 | | 0.020 | 0.038 | 0.531 | 0.595 | -0.057 | 0.095 |
| y1 | ~~ | cov5 | | -0.373 | 0.388 | -0.962 | 0.336 | -1.144 | 0.303 |
| m1 | ~~ | cov6 | | -0.039 | 0.020 | -1.981 | 0.048 | -0.080 | -0.001 |

| lhs | op | rhs | label | est | se | z | pvalue | ci.lower | ci.upper |
|---|---|---|---|---|---|---|---|---|---|
| y1 | ~~ | cov6 | | -0.178 | 0.142 | -1.250 | 0.211 | -0.468 | 0.097 |
| cov1 | ~~ | cov1 | | 0.250 | 0.006 | 44.155 | 0.000 | 0.229 | 0.250 |
| cov2 | ~~ | cov2 | | 0.926 | 0.263 | 3.523 | 0.000 | 0.432 | 1.489 |
| cov3 | ~~ | cov3 | | 1.116 | 0.434 | 2.571 | 0.010 | 0.476 | 2.046 |
| cov4 | ~~ | cov4 | | 14.307 | 2.304 | 6.210 | 0.000 | 9.642 | 18.652 |
| cov5 | ~~ | cov5 | | 3.096 | 0.645 | 4.796 | 0.000 | 1.939 | 4.450 |
| cov6 | ~~ | cov6 | | 0.627 | 0.085 | 7.399 | 0.000 | 0.467 | 0.805 |
| deltam | ~ | x | am2x | 0.035 | 0.025 | 1.416 | 0.157 | -0.015 | 0.083 |
| deltam | ~ | cov1 | | -0.046 | 0.025 | -1.811 | 0.070 | -0.098 | 0.002 |
| deltam | ~ | cov2 | | -0.007 | 0.018 | -0.377 | 0.706 | -0.036 | 0.032 |
| deltam | ~ | cov3 | | 0.007 | 0.018 | 0.423 | 0.673 | -0.023 | 0.049 |
| deltam | ~ | cov4 | | -0.004 | 0.004 | -1.019 | 0.308 | -0.012 | 0.004 |
| deltam | ~ | cov5 | | -0.007 | 0.009 | -0.754 | 0.451 | -0.023 | 0.010 |
| deltam | ~ | cov6 | | 0.035 | 0.015 | 2.372 | 0.018 | 0.003 | 0.062 |
| deltay | ~ | x | cy2x | -0.203 | 0.403 | -0.504 | 0.614 | -1.026 | 0.514 |
| deltay | ~ | deltam | by2m2 | -2.257 | 2.405 | -0.938 | 0.348 | -7.513 | 2.055 |
| deltay | ~ | cov1 | | 0.722 | 0.425 | 1.699 | 0.089 | -0.103 | 1.532 |
| deltay | ~ | cov2 | | 0.212 | 0.345 | 0.614 | 0.539 | -0.540 | 0.858 |
| deltay | ~ | cov3 | | -0.227 | 0.379 | -0.600 | 0.549 | -1.143 | 0.282 |
| deltay | ~ | cov4 | | -0.059 | 0.062 | -0.952 | 0.341 | -0.179 | 0.062 |
| deltay | ~ | cov5 | | 0.085 | 0.145 | 0.589 | 0.556 | -0.182 | 0.384 |
| deltay | ~ | cov6 | | 0.134 | 0.249 | 0.541 | 0.589 | -0.328 | 0.622 |
| m1 | ~~ | m1 | | 0.034 | 0.007 | 5.142 | 0.000 | 0.023 | 0.049 |
| y1 | ~~ | y1 | | 2.288 | 0.413 | 5.537 | 0.000 | 1.616 | 3.194 |
| x | ~~ | x | | 0.249 | 0.007 | 36.005 | 0.000 | 0.224 | 0.250 |
| x | ~1 | | | 0.468 | 0.063 | 7.392 | 0.000 | 0.339 | 0.581 |
| cov1 | ~1 | | | 0.484 | 0.063 | 7.707 | 0.000 | 0.355 | 0.613 |
| cov2 | ~1 | | | 0.532 | 0.121 | 4.409 | 0.000 | 0.306 | 0.774 |
| cov3 | ~1 | | | 0.694 | 0.134 | 5.185 | 0.000 | 0.452 | 0.968 |
| cov4 | ~1 | | | 9.823 | 0.486 | 20.196 | 0.000 | 8.839 | 10.758 |
| cov5 | ~1 | | | 6.968 | 0.223 | 31.179 | 0.000 | 6.516 | 7.403 |
| cov6 | ~1 | | | 0.952 | 0.103 | 9.280 | 0.000 | 0.758 | 1.161 |
| indirect | := | am2x*by2m2 | indirect | -0.079 | 0.106 | -0.746 | 0.456 | -0.343 | 0.090 |
| direct | := | cy2x | direct | -0.203 | 0.404 | -0.504 | 0.614 | -1.026 | 0.514 |
| total | := | cy2x+ (am2x*by2m2) | total | -0.283 | 0.418 | -0.676 | 0.499 | -1.128 | 0.468 |

#### 3.8 Mediation parietal cluster wide

##### *PHP*

Table 14: Overview of mediation model (difference score, 2W-LCS) using PHP and PAF (8–12 Hz) from a parietal cluster of electrodes identified by cluster based permutation analyses. Covariate/confounding variables are sex (male/female), depressive symptoms, anxiety symptoms, perceived stress, sleep quality, and recent pain. $\chi^2(24) = 25.98, p = 0.35$, CFI = 0.99, RMSEA = 0.037, SRMR = 0.079.

| lhs | op | rhs | label | est | se | z | pvalue | ci.lower | ci.upper |
|---|---|---|---|---|---|---|---|---|---|
| deltam | =~ | m2 | | 1.000 | 0.000 | NA | NA | 1.000 | 1.000 |
| deltam | ~~ | deltam | | 0.017 | 0.003 | 5.816 | 0.000 | 0.009 | 0.020 |
| deltam | ~1 | | | 0.117 | 0.121 | 0.969 | 0.332 | -0.121 | 0.369 |
| m2 | ~ | m1 | | 1.000 | 0.000 | NA | NA | 1.000 | 1.000 |
| m2 | ~~ | m1 | | 0.000 | 0.000 | NA | NA | 0.000 | 0.000 |
| m2 | ~~ | m2 | | 0.000 | 0.000 | NA | NA | 0.000 | 0.000 |
| m2 | ~1 | | | 0.000 | 0.000 | NA | NA | 0.000 | 0.000 |
| m1 | ~1 | | | 10.000 | 0.055 | 182.616 | 0.000 | 9.891 | 10.104 |
| deltay | =~ | y2 | | 1.000 | 0.000 | NA | NA | 1.000 | 1.000 |
| deltay | ~~ | deltay | | 0.610 | 0.151 | 4.036 | 0.000 | 0.251 | 0.797 |
| deltay | ~1 | | | 1.462 | 0.862 | 1.697 | 0.090 | -0.307 | 3.077 |
| y2 | ~ | y1 | | 1.000 | 0.000 | NA | NA | 1.000 | 1.000 |
| y2 | ~~ | y1 | | 0.000 | 0.000 | NA | NA | 0.000 | 0.000 |
| y2 | ~~ | y2 | | 0.000 | 0.000 | NA | NA | 0.000 | 0.000 |
| y2 | ~1 | | | 0.000 | 0.000 | NA | NA | 0.000 | 0.000 |
| y1 | ~1 | | | 5.655 | 0.306 | 18.505 | 0.000 | 5.037 | 6.250 |
| m1 | ~~ | y1 | | -0.109 | 0.144 | -0.756 | 0.450 | -0.396 | 0.162 |
| m1 | ~~ | x | | -0.050 | 0.028 | -1.757 | 0.079 | -0.104 | 0.009 |
| y1 | ~~ | x | | 0.073 | 0.167 | 0.437 | 0.662 | -0.238 | 0.410 |
| m1 | ~~ | cov1 | | -0.046 | 0.029 | -1.590 | 0.112 | -0.108 | 0.009 |
| y1 | ~~ | cov1 | | -0.343 | 0.150 | -2.288 | 0.022 | -0.634 | -0.027 |
| m1 | ~~ | cov2 | | -0.007 | 0.050 | -0.141 | 0.888 | -0.107 | 0.090 |
| y1 | ~~ | cov2 | | -0.157 | 0.266 | -0.588 | 0.556 | -0.733 | 0.344 |
| cov2 | ~~ | cov3 | | 0.547 | 0.193 | 2.828 | 0.005 | 0.184 | 0.939 |
| m1 | ~~ | cov3 | | -0.057 | 0.049 | -1.168 | 0.243 | -0.156 | 0.045 |
| y1 | ~~ | cov3 | | 0.260 | 0.289 | 0.902 | 0.367 | -0.362 | 0.781 |
| m1 | ~~ | cov4 | | 0.071 | 0.248 | 0.288 | 0.774 | -0.403 | 0.545 |
| y1 | ~~ | cov4 | | -0.847 | 1.294 | -0.655 | 0.513 | -3.253 | 1.871 |
| m1 | ~~ | cov5 | | 0.056 | 0.101 | 0.553 | 0.580 | -0.171 | 0.242 |
| y1 | ~~ | cov5 | | -0.326 | 0.444 | -0.733 | 0.463 | -1.176 | 0.666 |
| m1 | ~~ | cov6 | | -0.046 | 0.047 | -0.980 | 0.327 | -0.136 | 0.053 |

| lhs | op | rhs | label | est | se | z | pvalue | ci.lower | ci.upper |
|---|---|---|---|---|---|---|---|---|---|
| y1 | ~~ | cov6 | | -0.015 | 0.216 | -0.070 | 0.944 | -0.450 | 0.382 |
| cov1 | ~~ | cov1 | | 0.249 | 0.007 | 33.410 | 0.000 | 0.222 | 0.250 |
| cov2 | ~~ | cov2 | | 0.916 | 0.281 | 3.264 | 0.001 | 0.416 | 1.500 |
| cov3 | ~~ | cov3 | | 1.116 | 0.451 | 2.474 | 0.013 | 0.481 | 2.050 |
| cov4 | ~~ | cov4 | | 14.612 | 2.293 | 6.374 | 0.000 | 9.917 | 19.114 |
| cov5 | ~~ | cov5 | | 3.183 | 0.658 | 4.839 | 0.000 | 1.949 | 4.543 |
| cov6 | ~~ | cov6 | | 0.616 | 0.087 | 7.112 | 0.000 | 0.449 | 0.783 |
| deltam | ~ | x | am2x | 0.105 | 0.035 | 3.011 | 0.003 | 0.033 | 0.168 |
| deltam | ~ | cov1 | | -0.040 | 0.037 | -1.107 | 0.268 | -0.118 | 0.023 |
| deltam | ~ | cov2 | | -0.024 | 0.024 | -1.011 | 0.312 | -0.063 | 0.036 |
| deltam | ~ | cov3 | | 0.026 | 0.024 | 1.049 | 0.294 | -0.040 | 0.064 |
| deltam | ~ | cov4 | | -0.003 | 0.005 | -0.512 | 0.608 | -0.013 | 0.007 |
| deltam | ~ | cov5 | | -0.016 | 0.014 | -1.178 | 0.239 | -0.044 | 0.013 |
| deltam | ~ | cov6 | | 0.034 | 0.022 | 1.561 | 0.118 | -0.011 | 0.076 |
| deltay | ~ | x | cy2x | -0.671 | 0.311 | -2.156 | 0.031 | -1.290 | -0.107 |
| deltay | ~ | deltam | by2m2 | 0.218 | 0.827 | 0.264 | 0.792 | -1.319 | 1.865 |
| deltay | ~ | cov1 | | 0.130 | 0.273 | 0.478 | 0.633 | -0.383 | 0.684 |
| deltay | ~ | cov2 | | 0.291 | 0.166 | 1.749 | 0.080 | -0.052 | 0.592 |
| deltay | ~ | cov3 | | -0.059 | 0.141 | -0.418 | 0.676 | -0.419 | 0.116 |
| deltay | ~ | cov4 | | -0.096 | 0.049 | -1.955 | 0.051 | -0.187 | 0.000 |
| deltay | ~ | cov5 | | -0.009 | 0.070 | -0.132 | 0.895 | -0.153 | 0.136 |
| deltay | ~ | cov6 | | -0.175 | 0.130 | -1.347 | 0.178 | -0.418 | 0.091 |
| m1 | ~~ | m1 | | 0.189 | 0.034 | 5.584 | 0.000 | 0.135 | 0.264 |
| y1 | ~~ | y1 | | 5.255 | 0.764 | 6.882 | 0.000 | 3.865 | 6.948 |
| x | ~~ | x | | 0.249 | 0.008 | 31.998 | 0.000 | 0.222 | 0.250 |
| x | ~1 | | | 0.467 | 0.062 | 7.514 | 0.000 | 0.333 | 0.583 |
| cov1 | ~1 | | | 0.467 | 0.063 | 7.366 | 0.000 | 0.333 | 0.600 |
| cov2 | ~1 | | | 0.517 | 0.130 | 3.987 | 0.000 | 0.300 | 0.800 |
| cov3 | ~1 | | | 0.683 | 0.134 | 5.083 | 0.000 | 0.433 | 0.967 |
| cov4 | ~1 | | | 9.767 | 0.503 | 19.401 | 0.000 | 8.800 | 10.767 |
| cov5 | ~1 | | | 6.983 | 0.231 | 30.290 | 0.000 | 6.517 | 7.433 |
| cov6 | ~1 | | | 0.983 | 0.097 | 10.172 | 0.000 | 0.800 | 1.167 |
| indirect | := | am2x*by2m2 | indirect | 0.023 | 0.088 | 0.261 | 0.794 | -0.121 | 0.229 |
| direct | := | cy2x | direct | -0.671 | 0.311 | -2.155 | 0.031 | -1.290 | -0.107 |
| total | := | cy2x+ (am2x*by2m2) | total | -0.648 | 0.303 | -2.137 | 0.033 | -1.257 | -0.095 |

##### *CPA*

Table 15: Overview of mediation model (difference score, 2W-LCS) using CPA and PAF (8–12 Hz) from a parietal cluster of electrodes identified by cluster based permutation analyses. Covariate/confounding variables are sex (male/female), depressive symptoms, anxiety symptoms, perceived stress, sleep quality, and recent pain. $\chi^2(24) = 31.05, p = 0.15$, CFI = 0.97, RMSEA = 0.069, SRMR = 0.087.

| lhs | op | rhs | label | est | se | z | pvalue | ci.lower | ci.upper |
|---|---|---|---|---|---|---|---|---|---|
| deltam | =~ | m2 | | 1.000 | 0.000 | NA | NA | 1.000 | 1.000 |
| deltam | ~~ | deltam | | 0.016 | 0.003 | 5.529 | 0.000 | 0.009 | 0.020 |
| deltam | ~1 | | | 0.120 | 0.111 | 1.074 | 0.283 | -0.103 | 0.343 |
| m2 | ~ | m1 | | 1.000 | 0.000 | NA | NA | 1.000 | 1.000 |
| m2 | ~~ | m1 | | 0.000 | 0.000 | NA | NA | 0.000 | 0.000 |
| m2 | ~~ | m2 | | 0.000 | 0.000 | NA | NA | 0.000 | 0.000 |
| m2 | ~1 | | | 0.000 | 0.000 | NA | NA | 0.000 | 0.000 |
| m1 | ~1 | | | 9.999 | 0.055 | 182.260 | 0.000 | 9.891 | 10.109 |
| deltay | =~ | y2 | | 1.000 | 0.000 | NA | NA | 1.000 | 1.000 |
| deltay | ~~ | deltay | | 2.228 | 0.414 | 5.377 | 0.000 | 1.046 | 2.594 |
| deltay | ~1 | | | -1.273 | 1.469 | -0.867 | 0.386 | -3.949 | 1.630 |
| y2 | ~ | y1 | | 1.000 | 0.000 | NA | NA | 1.000 | 1.000 |
| y2 | ~~ | y1 | | 0.000 | 0.000 | NA | NA | 0.000 | 0.000 |
| y2 | ~~ | y2 | | 0.000 | 0.000 | NA | NA | 0.000 | 0.000 |
| y2 | ~1 | | | 0.000 | 0.000 | NA | NA | 0.000 | 0.000 |
| y1 | ~1 | | | 4.580 | 0.187 | 24.429 | 0.000 | 4.223 | 4.954 |
| m1 | ~~ | y1 | | -0.031 | 0.100 | -0.312 | 0.755 | -0.213 | 0.164 |
| m1 | ~~ | x | | -0.044 | 0.027 | -1.622 | 0.105 | -0.100 | 0.008 |
| y1 | ~~ | x | | 0.159 | 0.100 | 1.586 | 0.113 | -0.037 | 0.358 |
| m1 | ~~ | cov1 | | -0.048 | 0.028 | -1.697 | 0.090 | -0.105 | 0.008 |
| y1 | ~~ | cov1 | | -0.124 | 0.099 | -1.244 | 0.213 | -0.316 | 0.063 |
| m1 | ~~ | cov2 | | -0.020 | 0.049 | -0.406 | 0.685 | -0.117 | 0.078 |
| y1 | ~~ | cov2 | | -0.271 | 0.202 | -1.344 | 0.179 | -0.679 | 0.123 |
| cov2 | ~~ | cov3 | | 0.566 | 0.181 | 3.123 | 0.002 | 0.219 | 0.936 |
| m1 | ~~ | cov3 | | -0.069 | 0.048 | -1.432 | 0.152 | -0.160 | 0.028 |
| y1 | ~~ | cov3 | | -0.115 | 0.253 | -0.456 | 0.648 | -0.629 | 0.350 |
| m1 | ~~ | cov4 | | 0.080 | 0.230 | 0.347 | 0.729 | -0.391 | 0.541 |
| y1 | ~~ | cov4 | | -0.226 | 0.757 | -0.298 | 0.765 | -1.753 | 1.220 |
| m1 | ~~ | cov5 | | 0.051 | 0.097 | 0.522 | 0.602 | -0.162 | 0.227 |
| y1 | ~~ | cov5 | | -0.373 | 0.404 | -0.923 | 0.356 | -1.138 | 0.459 |
| m1 | ~~ | cov6 | | -0.046 | 0.047 | -0.982 | 0.326 | -0.138 | 0.043 |
| y1 | ~~ | cov6 | | -0.178 | 0.145 | -1.227 | 0.220 | -0.469 | 0.094 |
| cov1 | ~~ | cov1 | | 0.250 | 0.006 | 41.697 | 0.000 | 0.229 | 0.250 |
| cov2 | ~~ | cov2 | | 0.926 | 0.268 | 3.454 | 0.001 | 0.450 | 1.482 |

| lhs | op | rhs | label | est | se | z | pvalue | ci.lower | ci.upper |
|---|---|---|---|---|---|---|---|---|---|
| cov3 | ~~ | cov3 | | 1.116 | 0.426 | 2.621 | 0.009 | 0.504 | 1.990 |
| cov4 | ~~ | cov4 | | 14.307 | 2.418 | 5.916 | 0.000 | 9.652 | 19.060 |
| cov5 | ~~ | cov5 | | 3.096 | 0.621 | 4.983 | 0.000 | 1.939 | 4.379 |
| cov6 | ~~ | cov6 | | 0.627 | 0.085 | 7.393 | 0.000 | 0.465 | 0.790 |
| deltam | ~ | x | am2x | 0.106 | 0.032 | 3.283 | 0.001 | 0.041 | 0.166 |
| deltam | ~ | cov1 | | -0.037 | 0.035 | -1.045 | 0.296 | -0.111 | 0.029 |
| deltam | ~ | cov2 | | -0.024 | 0.026 | -0.922 | 0.357 | -0.068 | 0.034 |
| deltam | ~ | cov3 | | 0.025 | 0.024 | 1.074 | 0.283 | -0.035 | 0.061 |
| deltam | ~ | cov4 | | -0.002 | 0.005 | -0.450 | 0.652 | -0.013 | 0.008 |
| deltam | ~ | cov5 | | -0.017 | 0.013 | -1.322 | 0.186 | -0.041 | 0.010 |
| deltam | ~ | cov6 | | 0.031 | 0.020 | 1.581 | 0.114 | -0.009 | 0.068 |
| deltay | ~ | x | cy2x | -0.210 | 0.440 | -0.479 | 0.632 | -1.122 | 0.586 |
| deltay | ~ | deltam | by2m2 | -0.679 | 1.811 | -0.375 | 0.708 | -4.808 | 2.474 |
| deltay | ~ | cov1 | | 0.800 | 0.419 | 1.911 | 0.056 | -0.047 | 1.612 |
| deltay | ~ | cov2 | | 0.211 | 0.344 | 0.613 | 0.540 | -0.527 | 0.826 |
| deltay | ~ | cov3 | | -0.227 | 0.414 | -0.548 | 0.584 | -1.226 | 0.321 |
| deltay | ~ | cov4 | | -0.052 | 0.067 | -0.780 | 0.435 | -0.173 | 0.092 |
| deltay | ~ | cov5 | | 0.088 | 0.152 | 0.580 | 0.562 | -0.225 | 0.370 |
| deltay | ~ | cov6 | | 0.076 | 0.261 | 0.293 | 0.770 | -0.424 | 0.622 |
| m1 | ~~ | m1 | | 0.186 | 0.032 | 5.864 | 0.000 | 0.129 | 0.258 |
| y1 | ~~ | y1 | | 2.288 | 0.411 | 5.563 | 0.000 | 1.581 | 3.227 |
| x | ~~ | x | | 0.249 | 0.007 | 33.987 | 0.000 | 0.224 | 0.250 |
| x | ~1 | | | 0.468 | 0.064 | 7.345 | 0.000 | 0.339 | 0.597 |
| cov1 | ~1 | | | 0.484 | 0.064 | 7.569 | 0.000 | 0.355 | 0.613 |
| cov2 | ~1 | | | 0.532 | 0.121 | 4.396 | 0.000 | 0.306 | 0.774 |
| cov3 | ~1 | | | 0.694 | 0.130 | 5.317 | 0.000 | 0.435 | 0.952 |
| cov4 | ~1 | | | 9.823 | 0.486 | 20.210 | 0.000 | 8.871 | 10.838 |
| cov5 | ~1 | | | 6.968 | 0.227 | 30.758 | 0.000 | 6.516 | 7.435 |
| cov6 | ~1 | | | 0.952 | 0.100 | 9.562 | 0.000 | 0.758 | 1.161 |
| indirect | := | am2x*by2m2 | indirect | -0.072 | 0.197 | -0.367 | 0.714 | -0.552 | 0.265 |
| direct | := | cy2x | direct | -0.210 | 0.440 | -0.478 | 0.632 | -1.122 | 0.586 |
| total | := | cy2x+ (am2x*by2m2) | total | -0.283 | 0.435 | -0.650 | 0.516 | -1.203 | 0.493 |

#### 3.9 Mediation frontal cluster wide

##### *PHP*

Table 16: Overview of mediation model (difference score, 2W-LCS) using PHP and PAF (8–12 Hz) from a right-frontal cluster of electrodes identified by cluster based permutation analyses. Covariate/confounding variables are sex (male/female), depressive symptoms, anxiety symptoms, perceived stress, sleep quality, and recent pain. $\chi^2(24) = 26.08, p = 0.35$, CFI = 0.99, RMSEA = 0.038, SRMR = 0.079.

| lhs | op | rhs | label | est | se | z | pvalue | ci.lower | ci.upper |
|---|---|---|---|---|---|---|---|---|---|
| deltam | =~ | m2 | | 1.000 | 0.000 | NA | NA | 1.000 | 1.000 |
| deltam | ~~ | deltam | | 0.019 | 0.004 | 5.246 | 0.000 | 0.010 | 0.024 |
| deltam | ~1 | | | 0.107 | 0.118 | 0.907 | 0.364 | -0.151 | 0.334 |
| m2 | ~ | m1 | | 1.000 | 0.000 | NA | NA | 1.000 | 1.000 |
| m2 | ~~ | m1 | | 0.000 | 0.000 | NA | NA | 0.000 | 0.000 |
| m2 | ~~ | m2 | | 0.000 | 0.000 | NA | NA | 0.000 | 0.000 |
| m2 | ~1 | | | 0.000 | 0.000 | NA | NA | 0.000 | 0.000 |
| m1 | ~1 | | | 9.965 | 0.055 | 182.349 | 0.000 | 9.858 | 10.071 |
| deltay | =~ | y2 | | 1.000 | 0.000 | NA | NA | 1.000 | 1.000 |
| deltay | ~~ | deltay | | 0.610 | 0.150 | 4.074 | 0.000 | 0.240 | 0.789 |
| deltay | ~1 | | | 1.484 | 0.882 | 1.683 | 0.092 | -0.310 | 3.130 |
| y2 | ~ | y1 | | 1.000 | 0.000 | NA | NA | 1.000 | 1.000 |
| y2 | ~~ | y1 | | 0.000 | 0.000 | NA | NA | 0.000 | 0.000 |
| y2 | ~~ | y2 | | 0.000 | 0.000 | NA | NA | 0.000 | 0.000 |
| y2 | ~1 | | | 0.000 | 0.000 | NA | NA | 0.000 | 0.000 |
| y1 | ~1 | | | 5.655 | 0.306 | 18.505 | 0.000 | 5.037 | 6.250 |
| m1 | ~~ | y1 | | -0.127 | 0.149 | -0.851 | 0.395 | -0.431 | 0.152 |
| m1 | ~~ | x | | -0.057 | 0.028 | -2.030 | 0.042 | -0.112 | 0.001 |
| y1 | ~~ | x | | 0.073 | 0.167 | 0.437 | 0.662 | -0.238 | 0.410 |
| m1 | ~~ | cov1 | | -0.040 | 0.029 | -1.372 | 0.170 | -0.101 | 0.015 |
| y1 | ~~ | cov1 | | -0.343 | 0.150 | -2.288 | 0.022 | -0.634 | -0.027 |
| m1 | ~~ | cov2 | | -0.004 | 0.050 | -0.072 | 0.942 | -0.105 | 0.091 |
| y1 | ~~ | cov2 | | -0.157 | 0.266 | -0.588 | 0.556 | -0.733 | 0.344 |
| cov2 | ~~ | cov3 | | 0.547 | 0.193 | 2.828 | 0.005 | 0.184 | 0.939 |
| m1 | ~~ | cov3 | | -0.039 | 0.048 | -0.816 | 0.415 | -0.137 | 0.060 |
| y1 | ~~ | cov3 | | 0.260 | 0.289 | 0.902 | 0.367 | -0.362 | 0.781 |
| m1 | ~~ | cov4 | | -0.012 | 0.247 | -0.050 | 0.960 | -0.478 | 0.456 |
| y1 | ~~ | cov4 | | -0.847 | 1.294 | -0.655 | 0.513 | -3.253 | 1.871 |
| m1 | ~~ | cov5 | | 0.050 | 0.102 | 0.491 | 0.623 | -0.177 | 0.240 |
| y1 | ~~ | cov5 | | -0.326 | 0.444 | -0.733 | 0.463 | -1.176 | 0.666 |
| m1 | ~~ | cov6 | | -0.045 | 0.048 | -0.949 | 0.343 | -0.137 | 0.051 |

| lhs | op | rhs | label | est | se | z | pvalue | ci.lower | ci.upper |
|---|---|---|---|---|---|---|---|---|---|
| y1 | ~~ | cov6 | | -0.015 | 0.216 | -0.070 | 0.944 | -0.450 | 0.382 |
| cov1 | ~~ | cov1 | | 0.249 | 0.007 | 33.410 | 0.000 | 0.222 | 0.250 |
| cov2 | ~~ | cov2 | | 0.916 | 0.281 | 3.264 | 0.001 | 0.416 | 1.500 |
| cov3 | ~~ | cov3 | | 1.116 | 0.451 | 2.474 | 0.013 | 0.481 | 2.050 |
| cov4 | ~~ | cov4 | | 14.612 | 2.293 | 6.374 | 0.000 | 9.917 | 19.114 |
| cov5 | ~~ | cov5 | | 3.183 | 0.658 | 4.839 | 0.000 | 1.949 | 4.543 |
| cov6 | ~~ | cov6 | | 0.616 | 0.087 | 7.112 | 0.000 | 0.449 | 0.783 |
| deltam | ~ | x | am2x | 0.134 | 0.040 | 3.325 | 0.001 | 0.047 | 0.202 |
| deltam | ~ | cov1 | | -0.049 | 0.039 | -1.239 | 0.215 | -0.128 | 0.028 |
| deltam | ~ | cov2 | | -0.047 | 0.024 | -1.965 | 0.049 | -0.084 | 0.008 |
| deltam | ~ | cov3 | | 0.020 | 0.027 | 0.713 | 0.476 | -0.052 | 0.055 |
| deltam | ~ | cov4 | | 0.001 | 0.006 | 0.180 | 0.857 | -0.011 | 0.012 |
| deltam | ~ | cov5 | | -0.019 | 0.013 | -1.487 | 0.137 | -0.042 | 0.008 |
| deltam | ~ | cov6 | | 0.015 | 0.026 | 0.590 | 0.555 | -0.033 | 0.065 |
| deltay | ~ | x | cy2x | -0.652 | 0.277 | -2.359 | 0.018 | -1.211 | -0.132 |
| deltay | ~ | deltam | by2m2 | 0.036 | 0.918 | 0.039 | 0.969 | -1.878 | 1.793 |
| deltay | ~ | cov1 | | 0.123 | 0.266 | 0.463 | 0.644 | -0.395 | 0.650 |
| deltay | ~ | cov2 | | 0.287 | 0.157 | 1.833 | 0.067 | -0.041 | 0.569 |
| deltay | ~ | cov3 | | -0.054 | 0.142 | -0.381 | 0.704 | -0.417 | 0.127 |
| deltay | ~ | cov4 | | -0.097 | 0.048 | -2.007 | 0.045 | -0.187 | -0.002 |
| deltay | ~ | cov5 | | -0.012 | 0.074 | -0.165 | 0.869 | -0.154 | 0.139 |
| deltay | ~ | cov6 | | -0.168 | 0.127 | -1.329 | 0.184 | -0.395 | 0.103 |
| m1 | ~~ | m1 | | 0.188 | 0.033 | 5.648 | 0.000 | 0.136 | 0.263 |
| y1 | ~~ | y1 | | 5.255 | 0.764 | 6.882 | 0.000 | 3.865 | 6.948 |
| x | ~~ | x | | 0.249 | 0.008 | 31.998 | 0.000 | 0.222 | 0.250 |
| x | ~1 | | | 0.467 | 0.062 | 7.514 | 0.000 | 0.333 | 0.583 |
| cov1 | ~1 | | | 0.467 | 0.063 | 7.366 | 0.000 | 0.333 | 0.600 |
| cov2 | ~1 | | | 0.517 | 0.130 | 3.987 | 0.000 | 0.300 | 0.800 |
| cov3 | ~1 | | | 0.683 | 0.134 | 5.083 | 0.000 | 0.433 | 0.967 |
| cov4 | ~1 | | | 9.767 | 0.503 | 19.401 | 0.000 | 8.800 | 10.767 |
| cov5 | ~1 | | | 6.983 | 0.231 | 30.290 | 0.000 | 6.517 | 7.433 |
| cov6 | ~1 | | | 0.983 | 0.097 | 10.172 | 0.000 | 0.800 | 1.167 |
| indirect | := | am2x*by2m2 | indirect | 0.005 | 0.122 | 0.040 | 0.968 | -0.234 | 0.257 |
| direct | := | cy2x | direct | -0.652 | 0.277 | -2.357 | 0.018 | -1.211 | -0.132 |
| total | := | cy2x+ (am2x*by2m2) | total | -0.648 | 0.303 | -2.137 | 0.033 | -1.257 | -0.095 |

##### *CPA*

Table 17: Overview of mediation model (difference score, 2W-LCS) using CPA and PAF (8–12 Hz) from a right-frontal cluster of electrodes identified by cluster based permutation analyses. Covariate/confounding variables are sex (male/female), depressive symptoms, anxiety symptoms, perceived stress, sleep quality, and recent pain. $\chi^2(24) = 29.60, p = 0.20$, CFI = 0.97, RMSEA = 0.061, SRMR = 0.086.

| lhs | op | rhs | label | est | se | z | pvalue | ci.lower | ci.upper |
|---|---|---|---|---|---|---|---|---|---|
| deltam | =~ | m2 | | 1.000 | 0.000 | NA | NA | 1.000 | 1.000 |
| deltam | ~~ | deltam | | 0.019 | 0.003 | 5.549 | 0.000 | 0.010 | 0.023 |
| deltam | ~1 | | | 0.102 | 0.113 | 0.905 | 0.365 | -0.128 | 0.316 |
| m2 | ~ | m1 | | 1.000 | 0.000 | NA | NA | 1.000 | 1.000 |
| m2 | ~~ | m1 | | 0.000 | 0.000 | NA | NA | 0.000 | 0.000 |
| m2 | ~~ | m2 | | 0.000 | 0.000 | NA | NA | 0.000 | 0.000 |
| m2 | ~1 | | | 0.000 | 0.000 | NA | NA | 0.000 | 0.000 |
| m1 | ~1 | | | 9.964 | 0.054 | 184.025 | 0.000 | 9.859 | 10.076 |
| deltay | =~ | y2 | | 1.000 | 0.000 | NA | NA | 1.000 | 1.000 |
| deltay | ~~ | deltay | | 2.236 | 0.412 | 5.432 | 0.000 | 1.070 | 2.591 |
| deltay | ~1 | | | -1.362 | 1.454 | -0.936 | 0.349 | -4.096 | 1.523 |
| y2 | ~ | y1 | | 1.000 | 0.000 | NA | NA | 1.000 | 1.000 |
| y2 | ~~ | y1 | | 0.000 | 0.000 | NA | NA | 0.000 | 0.000 |
| y2 | ~~ | y2 | | 0.000 | 0.000 | NA | NA | 0.000 | 0.000 |
| y2 | ~1 | | | 0.000 | 0.000 | NA | NA | 0.000 | 0.000 |
| y1 | ~1 | | | 4.580 | 0.187 | 24.429 | 0.000 | 4.223 | 4.954 |
| m1 | ~~ | y1 | | -0.023 | 0.103 | -0.224 | 0.823 | -0.209 | 0.179 |
| m1 | ~~ | x | | -0.052 | 0.027 | -1.905 | 0.057 | -0.108 | 0.001 |
| y1 | ~~ | x | | 0.159 | 0.100 | 1.586 | 0.113 | -0.037 | 0.358 |
| m1 | ~~ | cov1 | | -0.042 | 0.028 | -1.492 | 0.136 | -0.101 | 0.013 |
| y1 | ~~ | cov1 | | -0.124 | 0.099 | -1.244 | 0.213 | -0.316 | 0.063 |
| m1 | ~~ | cov2 | | -0.015 | 0.049 | -0.313 | 0.754 | -0.111 | 0.083 |
| y1 | ~~ | cov2 | | -0.271 | 0.202 | -1.344 | 0.179 | -0.679 | 0.123 |
| cov2 | ~~ | cov3 | | 0.566 | 0.181 | 3.123 | 0.002 | 0.219 | 0.936 |
| m1 | ~~ | cov3 | | -0.050 | 0.046 | -1.072 | 0.284 | -0.139 | 0.043 |
| y1 | ~~ | cov3 | | -0.115 | 0.253 | -0.456 | 0.648 | -0.629 | 0.350 |
| m1 | ~~ | cov4 | | -0.005 | 0.229 | -0.020 | 0.984 | -0.487 | 0.449 |
| y1 | ~~ | cov4 | | -0.226 | 0.757 | -0.298 | 0.765 | -1.753 | 1.220 |
| m1 | ~~ | cov5 | | 0.046 | 0.099 | 0.467 | 0.640 | -0.168 | 0.221 |
| y1 | ~~ | cov5 | | -0.373 | 0.404 | -0.923 | 0.356 | -1.138 | 0.459 |
| m1 | ~~ | cov6 | | -0.044 | 0.047 | -0.942 | 0.346 | -0.139 | 0.045 |
| y1 | ~~ | cov6 | | -0.178 | 0.145 | -1.227 | 0.220 | -0.469 | 0.094 |
| cov1 | ~~ | cov1 | | 0.250 | 0.006 | 41.697 | 0.000 | 0.229 | 0.250 |

| lhs | op | rhs | label | est | se | z | pvalue | ci.lower | ci.upper |
|---|---|---|---|---|---|---|---|---|---|
| cov2 | ~~ | cov2 | | 0.926 | 0.268 | 3.454 | 0.001 | 0.450 | 1.482 |
| cov3 | ~~ | cov3 | | 1.116 | 0.426 | 2.621 | 0.009 | 0.504 | 1.990 |
| cov4 | ~~ | cov4 | | 14.307 | 2.418 | 5.916 | 0.000 | 9.652 | 19.060 |
| cov5 | ~~ | cov5 | | 3.096 | 0.621 | 4.983 | 0.000 | 1.939 | 4.379 |
| cov6 | ~~ | cov6 | | 0.627 | 0.085 | 7.393 | 0.000 | 0.465 | 0.790 |
| deltam | ~ | x | am2x | 0.138 | 0.036 | 3.778 | 0.000 | 0.055 | 0.200 |
| deltam | ~ | cov1 | | -0.058 | 0.038 | -1.514 | 0.130 | -0.134 | 0.013 |
| deltam | ~ | cov2 | | -0.049 | 0.024 | -2.044 | 0.041 | -0.089 | 0.009 |
| deltam | ~ | cov3 | | 0.018 | 0.027 | 0.680 | 0.497 | -0.054 | 0.050 |
| deltam | ~ | cov4 | | 0.001 | 0.006 | 0.098 | 0.922 | -0.011 | 0.011 |
| deltam | ~ | cov5 | | -0.018 | 0.012 | -1.519 | 0.129 | -0.041 | 0.010 |
| deltam | ~ | cov6 | | 0.021 | 0.023 | 0.905 | 0.366 | -0.028 | 0.063 |
| deltay | ~ | x | cy2x | -0.292 | 0.459 | -0.637 | 0.524 | -1.233 | 0.539 |
| deltay | ~ | deltam | by2m2 | 0.071 | 1.636 | 0.043 | 0.966 | -3.745 | 3.062 |
| deltay | ~ | cov1 | | 0.829 | 0.423 | 1.963 | 0.050 | -0.021 | 1.636 |
| deltay | ~ | cov2 | | 0.231 | 0.338 | 0.682 | 0.495 | -0.547 | 0.819 |
| deltay | ~ | cov3 | | -0.245 | 0.415 | -0.590 | 0.555 | -1.243 | 0.304 |
| deltay | ~ | cov4 | | -0.050 | 0.066 | -0.770 | 0.441 | -0.165 | 0.092 |
| deltay | ~ | cov5 | | 0.101 | 0.152 | 0.665 | 0.506 | -0.210 | 0.386 |
| deltay | ~ | cov6 | | 0.054 | 0.254 | 0.210 | 0.833 | -0.441 | 0.571 |
| m1 | ~~ | m1 | | 0.184 | 0.031 | 5.939 | 0.000 | 0.129 | 0.254 |
| y1 | ~~ | y1 | | 2.288 | 0.411 | 5.563 | 0.000 | 1.581 | 3.227 |
| x | ~~ | x | | 0.249 | 0.007 | 33.987 | 0.000 | 0.224 | 0.250 |
| x | ~1 | | | 0.468 | 0.064 | 7.345 | 0.000 | 0.339 | 0.597 |
| cov1 | ~1 | | | 0.484 | 0.064 | 7.569 | 0.000 | 0.355 | 0.613 |
| cov2 | ~1 | | | 0.532 | 0.121 | 4.396 | 0.000 | 0.306 | 0.774 |
| cov3 | ~1 | | | 0.694 | 0.130 | 5.317 | 0.000 | 0.435 | 0.952 |
| cov4 | ~1 | | | 9.823 | 0.486 | 20.210 | 0.000 | 8.871 | 10.838 |
| cov5 | ~1 | | | 6.968 | 0.227 | 30.758 | 0.000 | 6.516 | 7.435 |
| cov6 | ~1 | | | 0.952 | 0.100 | 9.562 | 0.000 | 0.758 | 1.161 |
| indirect | := | am2x*by2m2 | indirect | 0.010 | 0.220 | 0.044 | 0.965 | -0.482 | 0.441 |
| direct | := | cy2x | direct | -0.292 | 0.459 | -0.637 | 0.524 | -1.233 | 0.539 |
| total | := | cy2x+ (am2x*by2m2) | total | -0.283 | 0.435 | -0.650 | 0.516 | -1.203 | 0.493 |

#### 3.10 Mediation parietal-frontal clusters wide

##### ***PHP***

Table 18: Overview of mediation model (difference score, 2W-LCS) using PHP and PAF (8–12 Hz) from the average of all electrodes in the parietal and the right-frontal clusters identified by cluster based permutation analyses. Covariate/confounding variables are sex (male/female), depressive symptoms, anxiety symptoms, perceived stress, sleep quality, and recent pain. $\chi^2(24) = 26.03, p = 0.35$, CFI = 0.99, RMSEA = 0.038, SRMR = 0.079.

| lhs | op | rhs | label | est | se | z | pvalue | ci.lower | ci.upper |
|---|---|---|---|---|---|---|---|---|---|
| deltam | =~ | m2 | | 1.000 | 0.000 | NA | NA | 1.000 | 1.000 |
| deltam | ~~ | deltam | | 0.016 | 0.003 | 5.595 | 0.000 | 0.008 | 0.020 |
| deltam | ~1 | | | 0.114 | 0.117 | 0.981 | 0.326 | -0.136 | 0.353 |
| m2 | ~ | m1 | | 1.000 | 0.000 | NA | NA | 1.000 | 1.000 |
| m2 | ~~ | m1 | | 0.000 | 0.000 | NA | NA | 0.000 | 0.000 |
| m2 | ~~ | m2 | | 0.000 | 0.000 | NA | NA | 0.000 | 0.000 |
| m2 | ~1 | | | 0.000 | 0.000 | NA | NA | 0.000 | 0.000 |
| m1 | ~1 | | | 9.991 | 0.055 | 183.013 | 0.000 | 9.882 | 10.096 |
| deltay | =~ | y2 | | 1.000 | 0.000 | NA | NA | 1.000 | 1.000 |
| deltay | ~~ | deltay | | 0.610 | 0.151 | 4.034 | 0.000 | 0.248 | 0.793 |
| deltay | ~1 | | | 1.468 | 0.870 | 1.687 | 0.092 | -0.317 | 3.121 |
| y2 | ~ | y1 | | 1.000 | 0.000 | NA | NA | 1.000 | 1.000 |
| y2 | ~~ | y1 | | 0.000 | 0.000 | NA | NA | 0.000 | 0.000 |
| y2 | ~~ | y2 | | 0.000 | 0.000 | NA | NA | 0.000 | 0.000 |
| y2 | ~1 | | | 0.000 | 0.000 | NA | NA | 0.000 | 0.000 |
| y1 | ~1 | | | 5.655 | 0.306 | 18.505 | 0.000 | 5.037 | 6.250 |
| m1 | ~~ | y1 | | -0.113 | 0.145 | -0.782 | 0.434 | -0.406 | 0.159 |
| m1 | ~~ | x | | -0.052 | 0.028 | -1.829 | 0.067 | -0.106 | 0.006 |
| y1 | ~~ | x | | 0.073 | 0.167 | 0.437 | 0.662 | -0.238 | 0.410 |
| m1 | ~~ | cov1 | | -0.045 | 0.029 | -1.539 | 0.124 | -0.106 | 0.010 |
| y1 | ~~ | cov1 | | -0.343 | 0.150 | -2.288 | 0.022 | -0.634 | -0.027 |
| m1 | ~~ | cov2 | | -0.006 | 0.050 | -0.125 | 0.901 | -0.104 | 0.089 |
| y1 | ~~ | cov2 | | -0.157 | 0.266 | -0.588 | 0.556 | -0.733 | 0.344 |
| cov2 | ~~ | cov3 | | 0.547 | 0.193 | 2.828 | 0.005 | 0.184 | 0.939 |
| m1 | ~~ | cov3 | | -0.052 | 0.048 | -1.085 | 0.278 | -0.151 | 0.048 |
| y1 | ~~ | cov3 | | 0.260 | 0.289 | 0.902 | 0.367 | -0.362 | 0.781 |
| m1 | ~~ | cov4 | | 0.050 | 0.247 | 0.204 | 0.838 | -0.417 | 0.526 |
| y1 | ~~ | cov4 | | -0.847 | 1.294 | -0.655 | 0.513 | -3.253 | 1.871 |
| m1 | ~~ | cov5 | | 0.054 | 0.101 | 0.539 | 0.590 | -0.172 | 0.238 |
| y1 | ~~ | cov5 | | -0.326 | 0.444 | -0.733 | 0.463 | -1.176 | 0.666 |
| m1 | ~~ | cov6 | | -0.046 | 0.047 | -0.974 | 0.330 | -0.136 | 0.052 |

| lhs | op | rhs | label | est | se | z | pvalue | ci.lower | ci.upper |
|---|---|---|---|---|---|---|---|---|---|
| y1 | ~~ | cov6 | | -0.015 | 0.216 | -0.070 | 0.944 | -0.450 | 0.382 |
| cov1 | ~~ | cov1 | | 0.249 | 0.007 | 33.410 | 0.000 | 0.222 | 0.250 |
| cov2 | ~~ | cov2 | | 0.916 | 0.281 | 3.264 | 0.001 | 0.416 | 1.500 |
| cov3 | ~~ | cov3 | | 1.116 | 0.451 | 2.474 | 0.013 | 0.481 | 2.050 |
| cov4 | ~~ | cov4 | | 14.612 | 2.293 | 6.374 | 0.000 | 9.917 | 19.114 |
| cov5 | ~~ | cov5 | | 3.183 | 0.658 | 4.839 | 0.000 | 1.949 | 4.543 |
| cov6 | ~~ | cov6 | | 0.616 | 0.087 | 7.112 | 0.000 | 0.449 | 0.783 |
| deltam | ~ | x | am2x | 0.113 | 0.035 | 3.219 | 0.001 | 0.038 | 0.173 |
| deltam | ~ | cov1 | | -0.043 | 0.036 | -1.184 | 0.236 | -0.117 | 0.021 |
| deltam | ~ | cov2 | | -0.030 | 0.023 | -1.285 | 0.199 | -0.067 | 0.027 |
| deltam | ~ | cov3 | | 0.024 | 0.024 | 0.990 | 0.322 | -0.041 | 0.060 |
| deltam | ~ | cov4 | | -0.002 | 0.005 | -0.336 | 0.737 | -0.013 | 0.008 |
| deltam | ~ | cov5 | | -0.017 | 0.013 | -1.278 | 0.201 | -0.044 | 0.011 |
| deltam | ~ | cov6 | | 0.029 | 0.022 | 1.331 | 0.183 | -0.016 | 0.073 |
| deltay | ~ | x | cy2x | -0.667 | 0.303 | -2.201 | 0.028 | -1.276 | -0.116 |
| deltay | ~ | deltam | by2m2 | 0.177 | 0.877 | 0.201 | 0.840 | -1.529 | 1.867 |
| deltay | ~ | cov1 | | 0.129 | 0.271 | 0.476 | 0.634 | -0.387 | 0.681 |
| deltay | ~ | cov2 | | 0.291 | 0.164 | 1.771 | 0.077 | -0.048 | 0.588 |
| deltay | ~ | cov3 | | -0.057 | 0.141 | -0.408 | 0.683 | -0.422 | 0.118 |
| deltay | ~ | cov4 | | -0.096 | 0.049 | -1.960 | 0.050 | -0.190 | 0.001 |
| deltay | ~ | cov5 | | -0.010 | 0.071 | -0.138 | 0.891 | -0.153 | 0.138 |
| deltay | ~ | cov6 | | -0.173 | 0.129 | -1.344 | 0.179 | -0.405 | 0.093 |
| m1 | ~~ | m1 | | 0.188 | 0.033 | 5.605 | 0.000 | 0.135 | 0.260 |
| y1 | ~~ | y1 | | 5.255 | 0.764 | 6.882 | 0.000 | 3.865 | 6.948 |
| x | ~~ | x | | 0.249 | 0.008 | 31.998 | 0.000 | 0.222 | 0.250 |
| x | ~1 | | | 0.467 | 0.062 | 7.514 | 0.000 | 0.333 | 0.583 |
| cov1 | ~1 | | | 0.467 | 0.063 | 7.366 | 0.000 | 0.333 | 0.600 |
| cov2 | ~1 | | | 0.517 | 0.130 | 3.987 | 0.000 | 0.300 | 0.800 |
| cov3 | ~1 | | | 0.683 | 0.134 | 5.083 | 0.000 | 0.433 | 0.967 |
| cov4 | ~1 | | | 9.767 | 0.503 | 19.401 | 0.000 | 8.800 | 10.767 |
| cov5 | ~1 | | | 6.983 | 0.231 | 30.290 | 0.000 | 6.517 | 7.433 |
| cov6 | ~1 | | | 0.983 | 0.097 | 10.172 | 0.000 | 0.800 | 1.167 |
| indirect | := | am2x*by2m2 | indirect | 0.020 | 0.099 | 0.202 | 0.840 | -0.152 | 0.250 |
| direct | := | cy2x | direct | -0.667 | 0.303 | -2.199 | 0.028 | -1.276 | -0.116 |
| total | := | cy2x+ (am2x*by2m2) | total | -0.648 | 0.303 | -2.137 | 0.033 | -1.257 | -0.095 |

***CPA***

Table 19: Overview of mediation model (difference score, 2W-LCS) using CPA and PAF (8–12 Hz) from the average of all electrodes in the parietal and the right-frontal clusters identified by cluster based permutation analyses. Covariate/confounding variables are sex (male/female), depressive symptoms, anxiety symptoms, perceived stress, sleep quality, and recent pain. $\chi^2(24) = 30.56, p = 0.17$, CFI = 0.97, RMSEA = 0.066, SRMR = 0.086.

| lhs | op | rhs | label | est | se | z | pvalue | ci.lower | ci.upper |
|---|---|---|---|---|---|---|---|---|---|
| deltam | =~ | m2 | | 1.000 | 0.000 | NA | NA | 1.000 | 1.000 |
| deltam | ~~ | deltam | | 0.016 | 0.003 | 5.401 | 0.000 | 0.008 | 0.020 |
| deltam | ~1 | | | 0.115 | 0.108 | 1.063 | 0.288 | -0.100 | 0.328 |
| m2 | ~ | m1 | | 1.000 | 0.000 | NA | NA | 1.000 | 1.000 |
| m2 | ~~ | m1 | | 0.000 | 0.000 | NA | NA | 0.000 | 0.000 |
| m2 | ~~ | m2 | | 0.000 | 0.000 | NA | NA | 0.000 | 0.000 |
| m2 | ~1 | | | 0.000 | 0.000 | NA | NA | 0.000 | 0.000 |
| m1 | ~1 | | | 9.991 | 0.055 | 183.155 | 0.000 | 9.883 | 10.099 |
| deltay | =~ | y2 | | 1.000 | 0.000 | NA | NA | 1.000 | 1.000 |
| deltay | ~~ | deltay | | 2.232 | 0.414 | 5.387 | 0.000 | 1.049 | 2.597 |
| deltay | ~1 | | | -1.296 | 1.468 | -0.883 | 0.377 | -4.008 | 1.649 |
| y2 | ~ | y1 | | 1.000 | 0.000 | NA | NA | 1.000 | 1.000 |
| y2 | ~~ | y1 | | 0.000 | 0.000 | NA | NA | 0.000 | 0.000 |
| y2 | ~~ | y2 | | 0.000 | 0.000 | NA | NA | 0.000 | 0.000 |
| y2 | ~1 | | | 0.000 | 0.000 | NA | NA | 0.000 | 0.000 |
| y1 | ~1 | | | 4.580 | 0.187 | 24.429 | 0.000 | 4.223 | 4.954 |
| m1 | ~~ | y1 | | -0.029 | 0.101 | -0.290 | 0.772 | -0.212 | 0.168 |
| m1 | ~~ | x | | -0.046 | 0.027 | -1.697 | 0.090 | -0.102 | 0.006 |
| y1 | ~~ | x | | 0.159 | 0.100 | 1.586 | 0.113 | -0.037 | 0.358 |
| m1 | ~~ | cov1 | | -0.046 | 0.028 | -1.650 | 0.099 | -0.104 | 0.009 |
| y1 | ~~ | cov1 | | -0.124 | 0.099 | -1.244 | 0.213 | -0.316 | 0.063 |
| m1 | ~~ | cov2 | | -0.019 | 0.049 | -0.385 | 0.700 | -0.115 | 0.078 |
| y1 | ~~ | cov2 | | -0.271 | 0.202 | -1.344 | 0.179 | -0.679 | 0.123 |
| cov2 | ~~ | cov3 | | 0.566 | 0.181 | 3.123 | 0.002 | 0.219 | 0.936 |
| m1 | ~~ | cov3 | | -0.064 | 0.047 | -1.349 | 0.177 | -0.155 | 0.032 |
| y1 | ~~ | cov3 | | -0.115 | 0.253 | -0.456 | 0.648 | -0.629 | 0.350 |
| m1 | ~~ | cov4 | | 0.059 | 0.229 | 0.256 | 0.798 | -0.424 | 0.511 |
| y1 | ~~ | cov4 | | -0.226 | 0.757 | -0.298 | 0.765 | -1.753 | 1.220 |
| m1 | ~~ | cov5 | | 0.050 | 0.098 | 0.510 | 0.610 | -0.166 | 0.224 |
| y1 | ~~ | cov5 | | -0.373 | 0.404 | -0.923 | 0.356 | -1.138 | 0.459 |
| m1 | ~~ | cov6 | | -0.045 | 0.047 | -0.974 | 0.330 | -0.135 | 0.045 |
| y1 | ~~ | cov6 | | -0.178 | 0.145 | -1.227 | 0.220 | -0.469 | 0.094 |
| cov1 | ~~ | cov1 | | 0.250 | 0.006 | 41.697 | 0.000 | 0.229 | 0.250 |

| lhs | op | rhs | label | est | se | z | pvalue | ci.lower | ci.upper |
|---|---|---|---|---|---|---|---|---|---|
| cov2 | ~~ | cov2 | | 0.926 | 0.268 | 3.454 | 0.001 | 0.450 | 1.482 |
| cov3 | ~~ | cov3 | | 1.116 | 0.426 | 2.621 | 0.009 | 0.504 | 1.990 |
| cov4 | ~~ | cov4 | | 14.307 | 2.418 | 5.916 | 0.000 | 9.652 | 19.060 |
| cov5 | ~~ | cov5 | | 3.096 | 0.621 | 4.983 | 0.000 | 1.939 | 4.379 |
| cov6 | ~~ | cov6 | | 0.627 | 0.085 | 7.393 | 0.000 | 0.465 | 0.790 |
| deltam | ~ | x | am2x | 0.114 | 0.032 | 3.575 | 0.000 | 0.045 | 0.170 |
| deltam | ~ | cov1 | | -0.042 | 0.034 | -1.216 | 0.224 | -0.117 | 0.021 |
| deltam | ~ | cov2 | | -0.030 | 0.025 | -1.221 | 0.222 | -0.071 | 0.025 |
| deltam | ~ | cov3 | | 0.024 | 0.024 | 1.000 | 0.317 | -0.040 | 0.056 |
| deltam | ~ | cov4 | | -0.002 | 0.005 | -0.312 | 0.755 | -0.013 | 0.009 |
| deltam | ~ | cov5 | | -0.017 | 0.012 | -1.403 | 0.161 | -0.041 | 0.009 |
| deltam | ~ | cov6 | | 0.029 | 0.020 | 1.449 | 0.147 | -0.011 | 0.067 |
| deltay | ~ | x | cy2x | -0.225 | 0.446 | -0.503 | 0.615 | -1.124 | 0.595 |
| deltay | ~ | deltam | by2m2 | -0.509 | 1.853 | -0.275 | 0.784 | -4.637 | 2.724 |
| deltay | ~ | cov1 | | 0.804 | 0.421 | 1.908 | 0.056 | -0.045 | 1.617 |
| deltay | ~ | cov2 | | 0.212 | 0.342 | 0.619 | 0.536 | -0.536 | 0.823 |
| deltay | ~ | cov3 | | -0.232 | 0.415 | -0.559 | 0.576 | -1.235 | 0.316 |
| deltay | ~ | cov4 | | -0.051 | 0.066 | -0.771 | 0.440 | -0.171 | 0.092 |
| deltay | ~ | cov5 | | 0.091 | 0.153 | 0.597 | 0.551 | -0.219 | 0.375 |
| deltay | ~ | cov6 | | 0.070 | 0.260 | 0.268 | 0.789 | -0.427 | 0.615 |
| m1 | ~~ | m1 | | 0.184 | 0.031 | 5.890 | 0.000 | 0.128 | 0.254 |
| y1 | ~~ | y1 | | 2.288 | 0.411 | 5.563 | 0.000 | 1.581 | 3.227 |
| x | ~~ | x | | 0.249 | 0.007 | 33.987 | 0.000 | 0.224 | 0.250 |
| x | ~1 | | | 0.468 | 0.064 | 7.345 | 0.000 | 0.339 | 0.597 |
| cov1 | ~1 | | | 0.484 | 0.064 | 7.569 | 0.000 | 0.355 | 0.613 |
| cov2 | ~1 | | | 0.532 | 0.121 | 4.396 | 0.000 | 0.306 | 0.774 |
| cov3 | ~1 | | | 0.694 | 0.130 | 5.317 | 0.000 | 0.435 | 0.952 |
| cov4 | ~1 | | | 9.823 | 0.486 | 20.210 | 0.000 | 8.871 | 10.838 |
| cov5 | ~1 | | | 6.968 | 0.227 | 30.758 | 0.000 | 6.516 | 7.435 |
| cov6 | ~1 | | | 0.952 | 0.100 | 9.562 | 0.000 | 0.758 | 1.161 |
| indirect | := | am2x*by2m2 | indirect | -0.058 | 0.213 | -0.273 | 0.785 | -0.572 | 0.327 |
| direct | := | cy2x | direct | -0.225 | 0.447 | -0.503 | 0.615 | -1.124 | 0.595 |
| total | := | cy2x+ (am2x*by2m2) | total | -0.283 | 0.435 | -0.650 | 0.516 | -1.203 | 0.493 |

#### 3.11 Mediation sensorimotor independent component (IC) PAF wide

An automated independent component analysis identification was conducted to select one component containing alpha oscillations. This automatic pipeline conducted an independent component analysis 10 times for each resting state, and selected the component with the highest correlation to a template created of a sensorimotor alpha component (Figure 5).

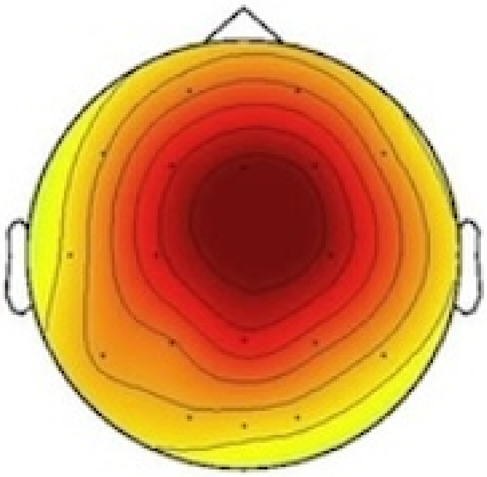

Figure 5: Template sensorimotor component used for automated independent component extraction.

The results of the PHP or CPA mediation models were not substantially different using the PAF calculated from independent components than that using the global PAF. For the PHP model, the total effect (b = -0.648, p = .033) and direct effects (b = -0.666, p = .035) were still significant, and there was still not significant indirect effect (b = 0.018, p = .726). The general fit was reduced, as although the CFI was above 0.90, akin to the original model, the RMSEA and SRMR were not below 0.08, unlike the original models (Little, 2013). For the CPA model, there were still no significant total (b = -0.371, p = .357), direct (b = -0.364, p = .386), or indirect effects (b = -0.007, p = .906), and the model fit also decreased, with CFI below 0.90 and RMSEA and SRMR above 0.08. See supplementary material (3.11). Note that still no correlations were seen between this IC sensorimotor PAF and pain (PHP: r = 0.11, p = .4; CPA: r = -0.064, p = .63).

Interestingly, in both models, there was now no longer a significant a-path (PHP: b = 0.08, p = 0.292; CPA: b = 0.039, p = 0.575), unlike previously observed (PHP: b = 0.085, p = 0.018; CPA: b = 0.089, p = 0.011). We interpret this as supporting the previously highlighted difference between finding an effect on PAF globally but not in a sensorimotor ROI (and now a sensorimotor IC), justifying the exploratory CBPA and the suggestion in the discussion to explore methodology.

##### *PHP*

Table 20: Overview of mediation model (difference score, 2W-LCS) using PHP and PAF (8–12 Hz) from a sensorimotor component extracted from an independent component analysis identified from an automated pipeline. Covariate/confounding variables are sex (male/female), depressive symptoms, anxiety symptoms, perceived stress, sleep quality, and recent pain. $\chi^2(24) = 41.24, p = 0.016$, CFI = 0.902, RMSEA = 0.109, SRMR = 0.124.

| lhs | op | rhs | label | est | se | z | pvalue | ci.lower | ci.upper |
|---|---|---|---|---|---|---|---|---|---|
| deltam | =~ | m2 | | 1.000 | 0.000 | NA | NA | 1.000 | 1.000 |
| deltam | ~~ | deltam | | 0.067 | 0.015 | 4.388 | 0.000 | 0.033 | 0.091 |
| deltam | ~1 | | | 0.033 | 0.190 | 0.176 | 0.860 | -0.351 | 0.412 |
| m2 | ~ | m1 | | 1.000 | 0.000 | NA | NA | 1.000 | 1.000 |
| m2 | ~~ | m1 | | 0.000 | 0.000 | NA | NA | 0.000 | 0.000 |
| m2 | ~~ | m2 | | 0.000 | 0.000 | NA | NA | 0.000 | 0.000 |
| m2 | ~1 | | | 0.000 | 0.000 | NA | NA | 0.000 | 0.000 |
| m1 | ~1 | | | 9.929 | 0.033 | 300.665 | 0.000 | 9.865 | 9.991 |
| deltay | =~ | y2 | | 1.000 | 0.000 | NA | NA | 1.000 | 1.000 |
| deltay | ~~ | deltay | | 0.607 | 0.151 | 4.023 | 0.000 | 0.251 | 0.801 |
| deltay | ~1 | | | 1.480 | 0.833 | 1.776 | 0.076 | -0.247 | 3.091 |
| y2 | ~ | y1 | | 1.000 | 0.000 | NA | NA | 1.000 | 1.000 |
| y2 | ~~ | y1 | | 0.000 | 0.000 | NA | NA | 0.000 | 0.000 |
| y2 | ~~ | y2 | | 0.000 | 0.000 | NA | NA | 0.000 | 0.000 |
| y2 | ~1 | | | 0.000 | 0.000 | NA | NA | 0.000 | 0.000 |
| y1 | ~1 | | | 5.655 | 0.306 | 18.505 | 0.000 | 5.037 | 6.250 |
| m1 | ~~ | y1 | | 0.082 | 0.075 | 1.092 | 0.275 | -0.064 | 0.227 |
| m1 | ~~ | x | | -0.034 | 0.017 | -1.917 | 0.055 | -0.066 | 0.001 |
| y1 | ~~ | x | | 0.073 | 0.167 | 0.437 | 0.662 | -0.238 | 0.410 |
| m1 | ~~ | cov1 | | -0.021 | 0.017 | -1.279 | 0.201 | -0.057 | 0.009 |
| y1 | ~~ | cov1 | | -0.343 | 0.150 | -2.288 | 0.022 | -0.634 | -0.027 |
| m1 | ~~ | cov2 | | 0.004 | 0.033 | 0.106 | 0.915 | -0.062 | 0.067 |
| y1 | ~~ | cov2 | | -0.157 | 0.266 | -0.588 | 0.556 | -0.733 | 0.344 |
| cov2 | ~~ | cov3 | | 0.547 | 0.193 | 2.828 | 0.005 | 0.184 | 0.939 |
| m1 | ~~ | cov3 | | -0.005 | 0.036 | -0.139 | 0.890 | -0.080 | 0.063 |
| y1 | ~~ | cov3 | | 0.260 | 0.289 | 0.902 | 0.367 | -0.362 | 0.781 |
| m1 | ~~ | cov4 | | 0.051 | 0.118 | 0.435 | 0.664 | -0.181 | 0.288 |
| y1 | ~~ | cov4 | | -0.847 | 1.294 | -0.655 | 0.513 | -3.253 | 1.871 |
| m1 | ~~ | cov5 | | -0.016 | 0.062 | -0.259 | 0.796 | -0.142 | 0.104 |
| y1 | ~~ | cov5 | | -0.326 | 0.444 | -0.733 | 0.463 | -1.176 | 0.666 |

| lhs | op | rhs | label | est | se | z | pvalue | ci.lower | ci.upper |
|---|---|---|---|---|---|---|---|---|---|
| m1 | ~~ | cov6 | | -0.002 | 0.030 | -0.074 | 0.941 | -0.063 | 0.055 |
| y1 | ~~ | cov6 | | -0.015 | 0.216 | -0.070 | 0.944 | -0.450 | 0.382 |
| cov1 | ~~ | cov1 | | 0.249 | 0.007 | 33.410 | 0.000 | 0.222 | 0.250 |
| cov2 | ~~ | cov2 | | 0.916 | 0.281 | 3.264 | 0.001 | 0.416 | 1.500 |
| cov3 | ~~ | cov3 | | 1.116 | 0.451 | 2.474 | 0.013 | 0.481 | 2.050 |
| cov4 | ~~ | cov4 | | 14.612 | 2.293 | 6.374 | 0.000 | 9.917 | 19.114 |
| cov5 | ~~ | cov5 | | 3.183 | 0.658 | 4.839 | 0.000 | 1.949 | 4.543 |
| cov6 | ~~ | cov6 | | 0.616 | 0.087 | 7.112 | 0.000 | 0.449 | 0.783 |
| deltam | ~ | x | am2x | 0.080 | 0.076 | 1.054 | 0.292 | -0.070 | 0.231 |
| deltam | ~ | cov1 | | -0.070 | 0.073 | -0.971 | 0.332 | -0.222 | 0.076 |
| deltam | ~ | cov2 | | -0.047 | 0.037 | -1.262 | 0.207 | -0.125 | 0.024 |
| deltam | ~ | cov3 | | -0.007 | 0.037 | -0.192 | 0.848 | -0.111 | 0.055 |
| deltam | ~ | cov4 | | -0.006 | 0.009 | -0.641 | 0.522 | -0.022 | 0.013 |
| deltam | ~ | cov5 | | 0.006 | 0.020 | 0.299 | 0.765 | -0.035 | 0.044 |
| deltam | ~ | cov6 | | -0.002 | 0.057 | -0.036 | 0.972 | -0.127 | 0.104 |
| deltay | ~ | x | cy2x | -0.666 | 0.315 | -2.113 | 0.035 | -1.302 | -0.104 |
| deltay | ~ | deltam | by2m2 | 0.227 | 0.441 | 0.515 | 0.607 | -0.842 | 0.967 |
| deltay | ~ | cov1 | | 0.137 | 0.268 | 0.513 | 0.608 | -0.407 | 0.658 |
| deltay | ~ | cov2 | | 0.296 | 0.164 | 1.801 | 0.072 | -0.038 | 0.585 |
| deltay | ~ | cov3 | | -0.052 | 0.139 | -0.371 | 0.711 | -0.421 | 0.129 |
| deltay | ~ | cov4 | | -0.095 | 0.048 | -1.980 | 0.048 | -0.184 | -0.004 |
| deltay | ~ | cov5 | | -0.014 | 0.068 | -0.210 | 0.834 | -0.151 | 0.130 |
| deltay | ~ | cov6 | | -0.167 | 0.125 | -1.341 | 0.180 | -0.393 | 0.087 |
| m1 | ~~ | m1 | | 0.065 | 0.014 | 4.565 | 0.000 | 0.038 | 0.095 |
| y1 | ~~ | y1 | | 5.255 | 0.764 | 6.882 | 0.000 | 3.865 | 6.948 |
| x | ~~ | x | | 0.249 | 0.008 | 31.998 | 0.000 | 0.222 | 0.250 |
| x | ~1 | | | 0.467 | 0.062 | 7.514 | 0.000 | 0.333 | 0.583 |
| cov1 | ~1 | | | 0.467 | 0.063 | 7.366 | 0.000 | 0.333 | 0.600 |
| cov2 | ~1 | | | 0.517 | 0.130 | 3.987 | 0.000 | 0.300 | 0.800 |
| cov3 | ~1 | | | 0.683 | 0.134 | 5.083 | 0.000 | 0.433 | 0.967 |
| cov4 | ~1 | | | 9.767 | 0.503 | 19.401 | 0.000 | 8.800 | 10.767 |
| cov5 | ~1 | | | 6.983 | 0.231 | 30.290 | 0.000 | 6.517 | 7.433 |
| cov6 | ~1 | | | 0.983 | 0.097 | 10.172 | 0.000 | 0.800 | 1.167 |
| indirect | := | am2x*by2m2 | indirect | 0.018 | 0.052 | 0.350 | 0.726 | -0.077 | 0.155 |
| direct | := | cy2x | direct | -0.666 | 0.315 | -2.112 | 0.035 | -1.302 | -0.104 |
| total | := | cy2x+(am2x*by2m2) | total | -0.648 | 0.303 | -2.137 | 0.033 | -1.257 | -0.095 |

***CPA***

Table 21: Overview of mediation model (difference score, 2W-LCS) using CPA and PAF (8–12 Hz) from a sensorimotor component extracted from an independent component analysis identified from an automated pipeline. Covariate/confounding variables are sex (male/female), depressive symptoms, anxiety symptoms, perceived stress, sleep quality, and recent pain. $\chi^2(24) = 38.53, p = 0.031$, CFI = 0.799, RMSEA = 0.1, SRMR = 0.119.

| lhs | op | rhs | label | est | se | z | pvalue | ci.lower | ci.upper |
|---|---|---|---|---|---|---|---|---|---|
| deltam | =~ | m2 | | 1.000 | 0.000 | NA | NA | 1.000 | 1.000 |
| deltam | ~~ | deltam | | 0.052 | 0.013 | 4.001 | 0.000 | 0.025 | 0.075 |
| deltam | ~1 | | | 0.098 | 0.151 | 0.646 | 0.519 | -0.242 | 0.366 |
| m2 | ~ | m1 | | 1.000 | 0.000 | NA | NA | 1.000 | 1.000 |
| m2 | ~~ | m1 | | 0.000 | 0.000 | NA | NA | 0.000 | 0.000 |
| m2 | ~~ | m2 | | 0.000 | 0.000 | NA | NA | 0.000 | 0.000 |
| m2 | ~1 | | | 0.000 | 0.000 | NA | NA | 0.000 | 0.000 |
| m1 | ~1 | | | 9.932 | 0.032 | 309.116 | 0.000 | 9.870 | 9.993 |
| deltay | =~ | y2 | | 1.000 | 0.000 | NA | NA | 1.000 | 1.000 |
| deltay | ~~ | deltay | | 2.153 | 0.403 | 5.343 | 0.000 | 0.977 | 2.577 |
| deltay | ~1 | | | -1.415 | 1.343 | -1.054 | 0.292 | -3.984 | 1.309 |
| y2 | ~ | y1 | | 1.000 | 0.000 | NA | NA | 1.000 | 1.000 |
| y2 | ~~ | y1 | | 0.000 | 0.000 | NA | NA | 0.000 | 0.000 |
| y2 | ~~ | y2 | | 0.000 | 0.000 | NA | NA | 0.000 | 0.000 |
| y2 | ~1 | | | 0.000 | 0.000 | NA | NA | 0.000 | 0.000 |
| y1 | ~1 | | | 4.603 | 0.189 | 24.334 | 0.000 | 4.247 | 4.966 |
| m1 | ~~ | y1 | | -0.020 | 0.051 | -0.397 | 0.691 | -0.116 | 0.088 |
| m1 | ~~ | x | | -0.031 | 0.016 | -1.956 | 0.050 | -0.060 | 0.002 |
| y1 | ~~ | x | | 0.153 | 0.099 | 1.534 | 0.125 | -0.054 | 0.349 |
| m1 | ~~ | cov1 | | -0.019 | 0.016 | -1.215 | 0.224 | -0.052 | 0.008 |
| y1 | ~~ | cov1 | | -0.118 | 0.100 | -1.176 | 0.240 | -0.332 | 0.066 |
| m1 | ~~ | cov2 | | -0.001 | 0.031 | -0.028 | 0.978 | -0.061 | 0.059 |
| y1 | ~~ | cov2 | | -0.278 | 0.207 | -1.342 | 0.179 | -0.730 | 0.113 |
| cov2 | ~~ | cov3 | | 0.569 | 0.184 | 3.091 | 0.002 | 0.230 | 0.953 |
| m1 | ~~ | cov3 | | -0.010 | 0.035 | -0.283 | 0.777 | -0.085 | 0.055 |
| y1 | ~~ | cov3 | | -0.118 | 0.249 | -0.472 | 0.637 | -0.671 | 0.368 |
| m1 | ~~ | cov4 | | 0.062 | 0.115 | 0.541 | 0.588 | -0.163 | 0.289 |
| y1 | ~~ | cov4 | | -0.258 | 0.758 | -0.341 | 0.733 | -1.768 | 1.145 |
| m1 | ~~ | cov5 | | -0.018 | 0.059 | -0.306 | 0.760 | -0.145 | 0.091 |
| y1 | ~~ | cov5 | | -0.346 | 0.392 | -0.882 | 0.378 | -1.110 | 0.411 |

| lhs | op | rhs | label | est | se | z | pvalue | ci.lower | ci.upper |
|---|---|---|---|---|---|---|---|---|---|
| m1 | ~~ | cov6 | | -0.005 | 0.029 | -0.181 | 0.856 | -0.064 | 0.051 |
| y1 | ~~ | cov6 | | -0.180 | 0.138 | -1.300 | 0.194 | -0.472 | 0.084 |
| cov1 | ~~ | cov1 | | 0.249 | 0.007 | 38.105 | 0.000 | 0.226 | 0.250 |
| cov2 | ~~ | cov2 | | 0.937 | 0.266 | 3.528 | 0.000 | 0.427 | 1.481 |
| cov3 | ~~ | cov3 | | 1.126 | 0.439 | 2.565 | 0.010 | 0.476 | 2.094 |
| cov4 | ~~ | cov4 | | 14.487 | 2.379 | 6.090 | 0.000 | 10.035 | 19.225 |
| cov5 | ~~ | cov5 | | 3.078 | 0.641 | 4.799 | 0.000 | 1.886 | 4.312 |
| cov6 | ~~ | cov6 | | 0.637 | 0.085 | 7.508 | 0.000 | 0.465 | 0.801 |
| deltam | ~ | x | am2x | 0.039 | 0.069 | 0.561 | 0.575 | -0.102 | 0.171 |
| deltam | ~ | cov1 | | -0.007 | 0.066 | -0.112 | 0.911 | -0.123 | 0.139 |
| deltam | ~ | cov2 | | -0.060 | 0.042 | -1.454 | 0.146 | -0.153 | 0.009 |
| deltam | ~ | cov3 | | 0.015 | 0.037 | 0.403 | 0.687 | -0.070 | 0.085 |
| deltam | ~ | cov4 | | -0.003 | 0.008 | -0.346 | 0.729 | -0.019 | 0.013 |
| deltam | ~ | cov5 | | -0.011 | 0.015 | -0.743 | 0.458 | -0.039 | 0.023 |
| deltam | ~ | cov6 | | -0.030 | 0.047 | -0.643 | 0.520 | -0.117 | 0.069 |
| deltay | ~ | x | cy2x | -0.364 | 0.420 | -0.867 | 0.386 | -1.306 | 0.337 |
| deltay | ~ | deltam | by2m2 | -0.191 | 0.854 | -0.224 | 0.823 | -2.007 | 1.302 |
| deltay | ~ | cov1 | | 0.905 | 0.401 | 2.258 | 0.024 | 0.090 | 1.657 |
| deltay | ~ | cov2 | | 0.203 | 0.383 | 0.530 | 0.596 | -0.670 | 0.863 |
| deltay | ~ | cov3 | | -0.240 | 0.406 | -0.593 | 0.553 | -1.199 | 0.326 |
| deltay | ~ | cov4 | | -0.058 | 0.063 | -0.918 | 0.359 | -0.177 | 0.073 |
| deltay | ~ | cov5 | | 0.126 | 0.142 | 0.889 | 0.374 | -0.151 | 0.424 |
| deltay | ~ | cov6 | | 0.055 | 0.230 | 0.238 | 0.812 | -0.394 | 0.523 |
| m1 | ~~ | m1 | | 0.064 | 0.014 | 4.604 | 0.000 | 0.039 | 0.093 |
| y1 | ~~ | y1 | | 2.295 | 0.419 | 5.472 | 0.000 | 1.616 | 3.264 |
| x | ~~ | x | | 0.249 | 0.007 | 35.515 | 0.000 | 0.226 | 0.250 |
| x | ~1 | | | 0.475 | 0.066 | 7.171 | 0.000 | 0.344 | 0.607 |
| cov1 | ~1 | | | 0.475 | 0.066 | 7.202 | 0.000 | 0.344 | 0.607 |
| cov2 | ~1 | | | 0.541 | 0.122 | 4.440 | 0.000 | 0.311 | 0.803 |
| cov3 | ~1 | | | 0.705 | 0.132 | 5.342 | 0.000 | 0.459 | 0.984 |
| cov4 | ~1 | | | 9.852 | 0.488 | 20.206 | 0.000 | 8.885 | 10.852 |
| cov5 | ~1 | | | 6.934 | 0.221 | 31.408 | 0.000 | 6.492 | 7.361 |
| cov6 | ~1 | | | 0.951 | 0.101 | 9.394 | 0.000 | 0.738 | 1.148 |
| indirect | := | am2x*by2m2 | indirect | -0.007 | 0.063 | -0.118 | 0.906 | -0.122 | 0.145 |
| direct | := | cy2x | direct | -0.364 | 0.420 | -0.867 | 0.386 | -1.306 | 0.337 |
| total | := | cy2x+(am2x*by2m2) | total | -0.371 | 0.403 | -0.921 | 0.357 | -1.245 | 0.318 |

### 4 Effect of nicotine on ongoing alpha oscillations

#### 4.1 Table of PAF and power values across four resting states

Table 22: Mean, standard deviation (SD), and median of peak alpha frequency (PAF), and mean alpha power ($\mu V^2$) for each resting state time point by group (i.e. nicotine and placebo gum). Data displayed for each variation in PAF calculation parameters, including centre of gravity (CoG) or max peak picking (peak), and narrow (9–11 Hz) or wide (8–12 Hz) alpha frequency bands. Resting state time points 1 and 2 were pre-gum chewing and time points 3 and 4 were post-gum chewing.

| PAF variable | Gum | Rest | n | Mean | SD | Median | Mean power | SD power |
|---|---|---|---|---|---|---|---|---|
| CoG sensorimotor (9–11 Hz) | Nicotine | 1 | 29 | 9.94 | 0.17 | 9.96 | 17.33 | 7.00 |
| | | 2 | 29 | 9.93 | 0.18 | 9.92 | 20.05 | 7.31 |
| | | 3 | 29 | 9.96 | 0.20 | 9.97 | 19.80 | 7.94 |
| | | 4 | 29 | 9.93 | 0.19 | 9.93 | 20.81 | 8.13 |
| | Placebo | 1 | 33 | 10.00 | 0.20 | 9.98 | 17.66 | 6.44 |
| | | 2 | 32 | 9.98 | 0.21 | 9.98 | 17.73 | 8.77 |
| | | 3 | 33 | 9.99 | 0.24 | 9.97 | 20.28 | 7.11 |
| | | 4 | 32 | 10.01 | 0.21 | 10.00 | 19.68 | 6.29 |
| CoG global (9–11 Hz) | Nicotine | 1 | 29 | 9.95 | 0.16 | 9.95 | 17.59 | 6.80 |
| | | 2 | 29 | 9.95 | 0.18 | 9.94 | 19.83 | 7.16 |
| | | 3 | 29 | 9.97 | 0.20 | 10.01 | 19.46 | 7.94 |
| | | 4 | 29 | 9.95 | 0.20 | 9.97 | 20.33 | 8.14 |
| | Placebo | 1 | 33 | 10.04 | 0.22 | 10.02 | 17.64 | 5.99 |
| | | 2 | 32 | 10.01 | 0.21 | 9.97 | 17.65 | 8.58 |
| | | 3 | 33 | 10.03 | 0.25 | 9.99 | 20.07 | 6.93 |
| | | 4 | 32 | 10.04 | 0.22 | 10.04 | 19.60 | 6.10 |
| CoG sensorimotor (8–12 Hz) | Nicotine | 1 | 29 | 9.82 | 0.36 | 9.80 | 25.83 | 8.12 |
| | | 2 | 29 | 9.82 | 0.37 | 9.83 | 28.79 | 7.28 |
| | | 3 | 29 | 9.88 | 0.42 | 9.88 | 28.73 | 7.30 |
| | | 4 | 29 | 9.81 | 0.38 | 9.84 | 29.55 | 8.36 |
| | Placebo | 1 | 33 | 9.97 | 0.38 | 9.97 | 26.50 | 7.00 |
| | | 2 | 32 | 9.94 | 0.40 | 9.92 | 26.63 | 9.51 |
| | | 3 | 33 | 9.96 | 0.41 | 9.94 | 29.47 | 7.05 |
| | | 4 | 32 | 9.98 | 0.37 | 10.00 | 28.80 | 6.20 |
| CoG global (8–12 Hz) | Nicotine | 1 | 29 | 9.86 | 0.37 | 9.85 | 25.89 | 7.64 |

| PAF variable | Gum | Rest | n | Mean | SD | Median | Mean power | SD power |
|---|---|---|---|---|---|---|---|---|
| | | 2 | 29 | 9.87 | 0.39 | 9.89 | 28.37 | 6.69 |
| | | 3 | 29 | 9.93 | 0.44 | 9.94 | 28.12 | 6.90 |
| | | 4 | 29 | 9.85 | 0.41 | 9.92 | 28.87 | 8.12 |
| | Placebo | 1 | 33 | 10.04 | 0.42 | 10.06 | 26.54 | 6.54 |
| | | 2 | 32 | 10.01 | 0.43 | 10.00 | 26.47 | 9.49 |
| | | 3 | 33 | 10.02 | 0.43 | 10.00 | 29.28 | 6.85 |
| | | 4 | 32 | 10.05 | 0.41 | 10.03 | 28.89 | 6.22 |
| Peak sensorimotor (9–11 Hz) | Nicotine | 1 | 29 | 9.91 | 0.28 | 9.95 | | |
| | | 2 | 29 | 9.92 | 0.30 | 9.88 | | |
| | | 3 | 29 | 9.95 | 0.33 | 9.96 | | |
| | | 4 | 29 | 9.91 | 0.32 | 9.91 | | |
| | Placebo | 1 | 33 | 10.02 | 0.34 | 9.99 | | |
| | | 2 | 32 | 9.99 | 0.35 | 9.93 | | |
| | | 3 | 33 | 10.01 | 0.40 | 9.96 | | |
| | | 4 | 32 | 10.04 | 0.36 | 10.03 | | |
| Peak global (9–11 Hz) | Nicotine | 1 | 29 | 9.93 | 0.28 | 9.94 | | |
| | | 2 | 29 | 9.94 | 0.30 | 9.92 | | |
| | | 3 | 29 | 9.98 | 0.33 | 10.03 | | |
| | | 4 | 29 | 9.94 | 0.33 | 9.98 | | |
| | Placebo | 1 | 33 | 10.08 | 0.37 | 10.05 | | |
| | | 2 | 32 | 10.04 | 0.37 | 9.93 | | |
| | | 3 | 33 | 10.06 | 0.40 | 10.01 | | |
| | | 4 | 32 | 10.09 | 0.36 | 10.07 | | |
| Peak sensorimotor (8–12 Hz) | Nicotine | 1 | 29 | 9.75 | 0.58 | 9.75 | | |
| | | 2 | 29 | 9.78 | 0.57 | 9.81 | | |
| | | 3 | 29 | 9.87 | 0.66 | 9.93 | | |
| | | 4 | 29 | 9.78 | 0.59 | 9.87 | | |
| | Placebo | 1 | 33 | 9.97 | 0.63 | 9.97 | | |
| | | 2 | 32 | 9.95 | 0.66 | 9.88 | | |
| | | 3 | 33 | 10.01 | 0.65 | 9.91 | | |
| | | 4 | 32 | 10.04 | 0.59 | 10.07 | | |
| Peak global (8–12 Hz) | Nicotine | 1 | 29 | 9.83 | 0.59 | 9.79 | | |
| | | 2 | 29 | 9.87 | 0.58 | 9.85 | | |
| | | 3 | 29 | 9.95 | 0.67 | 10.04 | | |

| PAF variable | Gum | Rest | n | Mean | SD | Median | Mean power | SD power |
|---|---|---|---|---|---|---|---|---|
| | | 4 | 29 | 9.85 | 0.61 | 9.93 | | |
| | Placebo | 1 | 33 | 10.06 | 0.66 | 10.15 | | |
| | | 2 | 32 | 10.02 | 0.68 | 10.00 | | |
| | | 3 | 33 | 10.07 | 0.64 | 10.05 | | |
| | | 4 | 32 | 10.10 | 0.61 | 10.02 | | |

#### 4.2 Effect of nicotine on PAF calculated with a wide frequency window (8–12 Hz): RM-ANOVAs 2x2

Nicotine gum increases the speed but not the power of alpha activity. A complete table of PAF and power values for all four time points is displayed in Supplementary material 3.1.

We found that participants' PAF became faster after chewing nicotine gum compared to after chewing placebo gum. This was seen as a statistically significant interaction (Table 23) between the gum group (i.e. nicotine/placebo) and time point (i.e. rest 1 pre-gum/rest 3 post-gum) on global PAF (8–12 Hz; $F(1, 60) = 6.02, p = .017, \eta^2_G = 0.003$), such that the change in PAF was significantly larger for the nicotine group ($t(28) = -2.16, p = .04$, 95% CI: [-0.14, -0.0037]) compared to the placebo group ($t(32) = 1.02, p = .31$, 95% CI: [-0.019, 0.058]). There were no significant differences in global PAF between the two groups at baseline ($t(60) = -1.84, p = 0.07$, 95% CI: [-0.39, 0.016]) or post-gum ($t(58.8) = -0.86, p = 0.39$, 95% CI: [-0.32, 0.13]). No interactions or main effects were seen for sensorimotor PAF (Table 23), suggesting that the change in PAF is not occurring at the sensorimotor ROI but at another topographical location.

Table 23: RM-ANOVA testing effects of gum group (i.e. nicotine/placebo) and resting state Time point (i.e. pre/post-gum) on sensorimotor and global peak alpha frequency (PAF), calculated with wide 8–12 Hz window and the centre of gravity method.

| | Effect | F(df) | *p*-value | $\eta^2_G$ |
|---|---|---|---|---|
| Sensorimotor | | | | |
| | Gum group | 1.33(1,60) | .25 | 0.02 |
| | Time point | 1.81(1,60) | .18 | $< 0.01$ |
| | Gum group:Time point | 3.51(1,60) | .066 | $< 0.01$ |
| Global | | | | |
| | Gum group | 1.81(1,60) | .18 | 0.03 |
| | Time point | 1.96(1,60) | .17 | $< 0.01$ |
| | Gum group:Time point | 6.02(1,60) | .017 | $< 0.01$ |

##### ***Effects of nicotine on alpha power (8–12 Hz): RM-ANOVAs 2x2***

No main effects of nicotine were seen on alpha power, but there was a main effect of time point on alpha power, where power increased irrespective of group (Table 24, Figure 6).

Table 24: RM-ANOVA testing effects of gum group (i.e. gum: nicotine/placebo) on global and sensorimotor power ($\mu V^2$) at each resting state (i.e. time point: pre/post-gum).

| | Effect | F(df) | *p*-value | $\eta^2_G$ |
|---|---|---|---|---|
| Sensorimotor | | | | |
| | Gum group | 0.17 (1,60) | .68 | < 0.01 |
| | Time point | 14.57 (1,60) | <.001 | 0.04 |
| | Gum group:Time point | 0.00 (1,60) | .97 | < 0.01 |
| Global | | | | |
| | Gum group | 0.32 (1,60) | .58 | < 0.01 |
| | Time point | 11.67 (1,60) | <.001 | 0.03 |
| | Gum group:Time point | 0.12 (1,60) | .73 | < 0.01 |

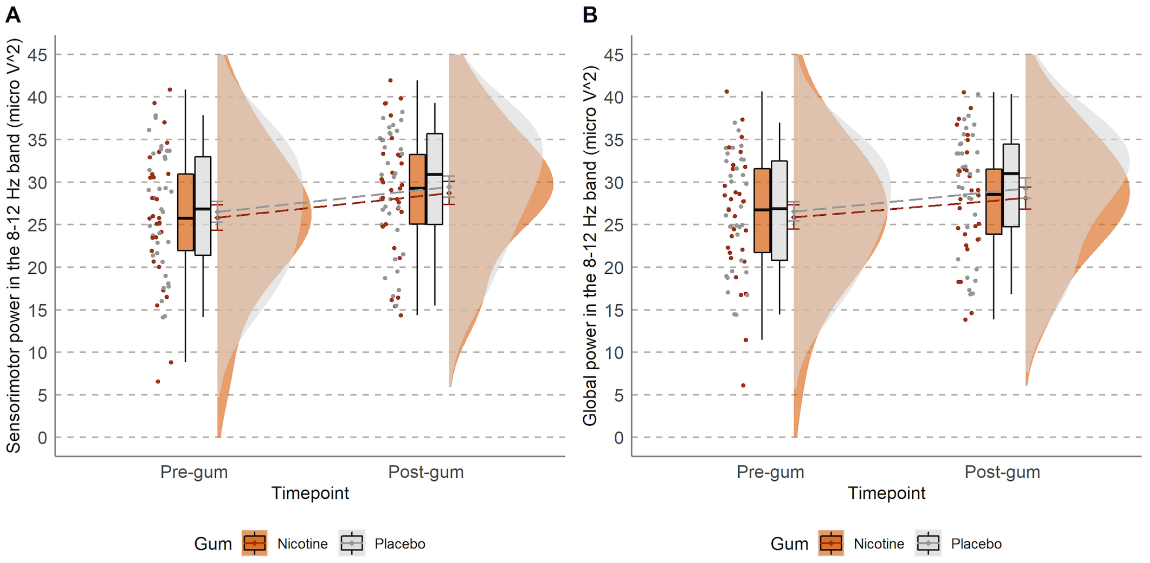

Figure 6: Average A) sensorimotor and B) global power ($\mu V^2$) in the 8–12 Hz band for all participants (N = 62) pre- and post-gum chewing, split by nicotine (in orange, n = 29) and placebo (in grey, n = 33) gum groups.

##### ***Effects of nicotine on PAF calculated with a narrow frequency window (9–11 Hz): RM-ANOVAs 2x2***

No effects of nicotine were seen on PAF calculated with a narrow window (Table 25, Figure 7).

Table 25: RM-ANOVA testing effects of gum group (i.e. gum: nicotine/placebo) on sensorimotor and global peak alpha frequency (PAF) calculated using a narrow frequency window (9–11 Hz) and the centre of gravity method at each resting state (i.e. rest: pre/post-gum).

| | Effect | F(df) | *p*-value | $\eta^2_G$ |
|---|---|---|---|---|
| Sensorimotor | | | | |
| | Gum group | 0.96(1,60) | .33 | 0.01 |
| | Time point | 0.20(1,60) | .66 | $< 0.01$ |
| | Gum group:Time point | 1.56(1,60) | .22 | $< 0.01$ |
| Global | | | | |
| | Gum group | 1.78(1,60) | .19 | 0.03 |
| | Time point | 0.15(1,60) | .70 | $< 0.01$ |
| | Gum group:Time point | 2.32(1,60) | .13 | $< 0.01$ |

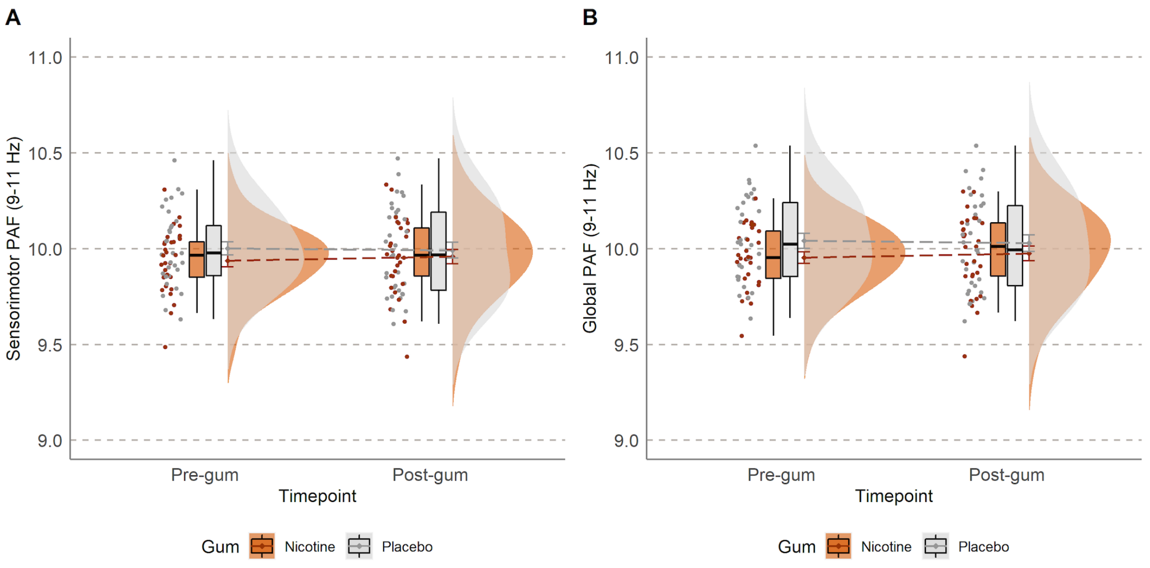

Figure 7: Mean A) sensorimotor and B) global peak alpha frequency (PAF) calculated with a narrow window (9–11 Hz) and the centre of gravity method for all participants (N = 62) pre- and post-gum chewing, split by nicotine (in orange, n = 29) and placebo (in grey, n = 33) gum groups.

***Effects of nicotine on PAF across four time points: ANOVAs PAF 4x2***

Examination of changes in PAF across all four resting state time points (Figure 8, Figure 9, and Figure 10) suggests an interaction effect between gum group and time point for global and sensorimotor PAF using a wide window (8–12 Hz; Table 26) but not for a narrow window (9–11 Hz; Table 27).

Table 26: RM-ANOVA testing effects of gum group (i.e. gum: nicotine/placebo) on global and sensorimotor peak alpha frequency (PAF) calculated using a wide frequency window (8–12 Hz) and the centre of gravity method at each of the four resting states (i.e. time point: four levels).

| | Effect | F(df) | $p$-value | $\eta^2_G$ |
|---|---|---|---|---|
| Global | | | | |
| | Gum group | 2.641(1,60) | .110 | 0.04 |
| | Time point | 1.398(3,174) | .245 | $< 0.01$ |
| | Gum group:Time point | 3.678(3,174) | .013 | $< 0.01$ |
| Sensorimotor | | | | |
| | Gum group | 2.180(1,60) | .145 | 0.03 |
| | Time point | 1.420(3,174) | .239 | $< 0.01$ |
| | Gum group:Time point | 2.878(3,174) | .038 | $< 0.01$ |

Table 27: RM-ANOVA testing effects of gum group (i.e. gum: nicotine/placebo) on global and sensorimotor peak alpha frequency (PAF) calculated using a narrow frequency window (9–11 Hz) and the centre of gravity method at each of the four resting states (i.e. time point: four levels).

| | Effect | F(df) | $p$-value | $\eta^2_G$ |
|---|---|---|---|---|
| Global | | | | |
| | Gum group | 2.463(1,60) | .12 | 0.04 |
| | Time point | 1.280(3,174) | .283 | $< 0.01$ |
| | Gum group:Time point | 1.735(3,174) | .162 | $< 0.01$ |
| Sensorimotor | | | | |
| | Gum group | 1.594(1,60) | .212 | 0.02 |
| | Time point | 0.756(3,174) | .510 | $< 0.01$ |
| | Gum group:Time point | 1.478(3,174) | .222 | $< 0.01$ |

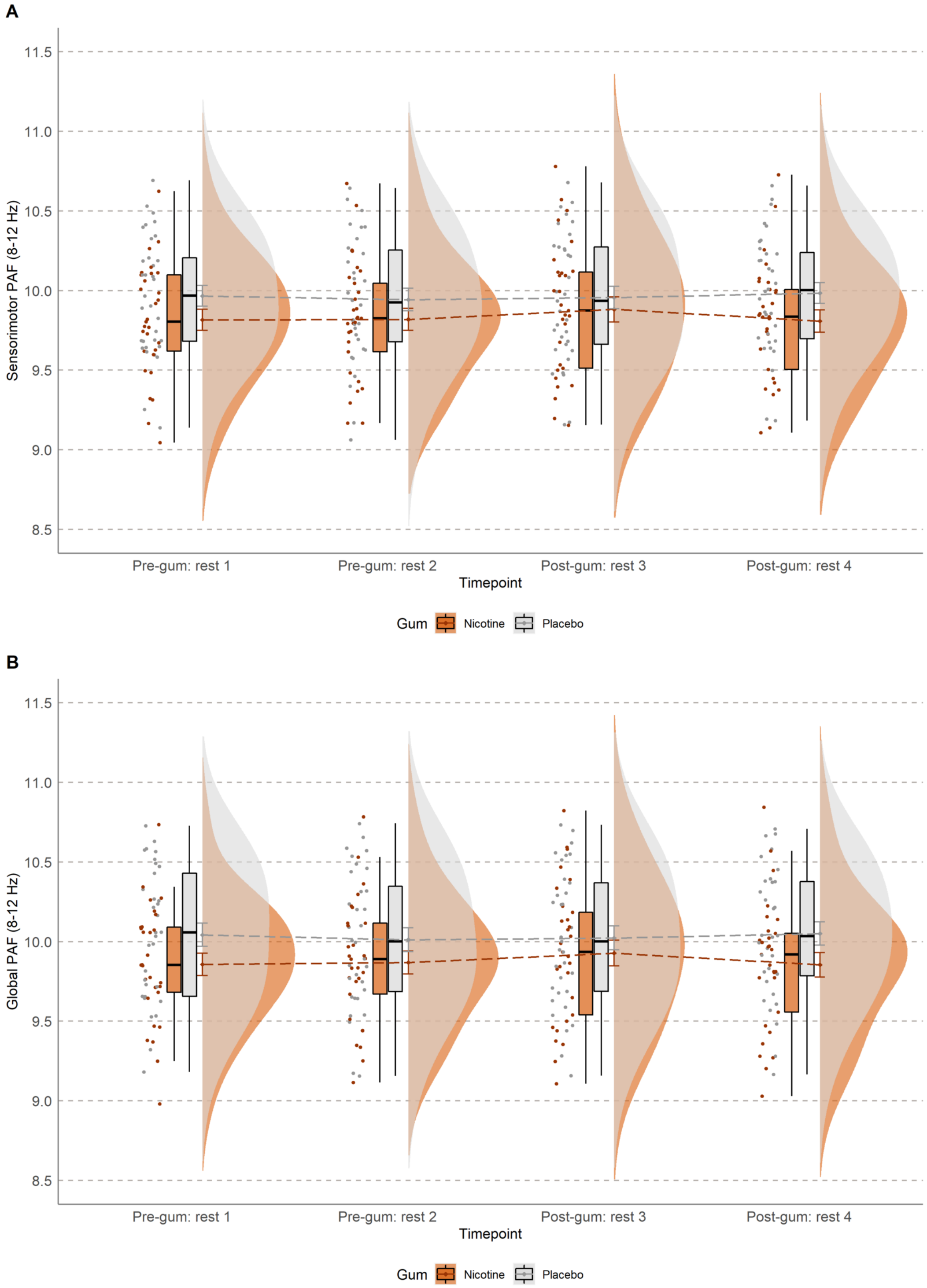

Figure 8: A) Sensorimotor and B) global peak alpha frequency (PAF), calculated with a wide frequency window (8–12 Hz) and the centre of gravity method, across four resting states for nicotine (orange, n = 29) and placebo (grey, n = 33) gum groups. Resting states 1 and 2 were pre-gum, while 3 and 4 were post-gum.

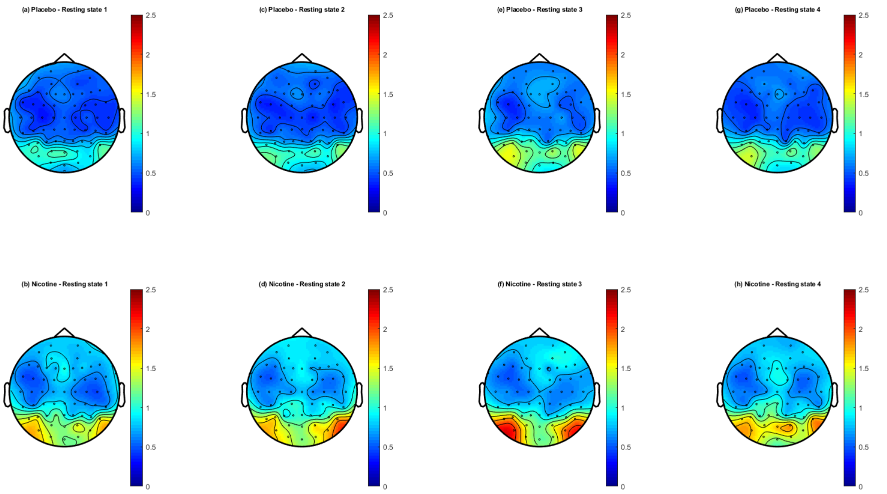

Figure 9: Topoplots of average power ($\mu V^2$) in the alpha range (8–12 Hz) for each electrode, separated by group, for four resting states across the experiment. Resting states 1 and 2 were pre-gum, while 3 and 4 were post-gum.

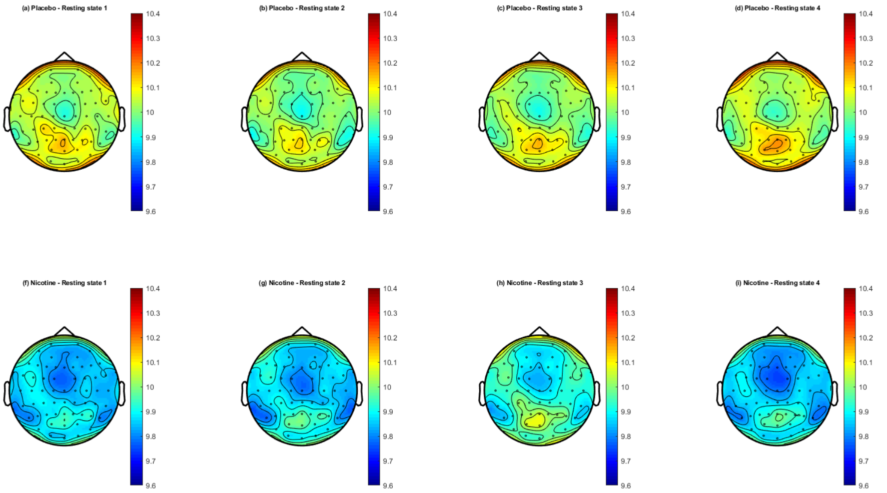

Figure 10: Topoplots of average peak alpha frequency (PAF) for each electrode, calculated with a wide frequency window (8–12 Hz) and the centre of gravity method, separated by group, for four resting states across the experiment. Resting states 1 and 2 were pre-gum, while 3 and 4 were post-gum.

### 5 Effects of nicotine on prolonged pain

##### *Effects of nicotine on PHP: RM-ANOVAs 2x2*

Without controlling for confounding variables, nicotine did not alter PHP ratings, indicating the importance of controlling for confounders. Specifically, there was no interaction between gum group (i.e. nicotine or placebo) and time point (i.e. rest 1 pre-gum/rest 3 post-gum) on PHP ratings ($F(1, 58) = 2.69, p = .11, \eta^2_G = 0.002$). Additionally, there were no significant main effects of gum group ($F(1, 58) = 0.045, p = .83, \eta^2_G = 0.00075$) or time point ($F(1, 58) = 1.53, p = .22, \eta^2_G = 0.00089$).

##### *Effects of nicotine on CPA: RM-ANOVAs 2x2*

There were no effects of nicotine on CPA ratings, as ratings decreased over time irrespective of which gum was chewed. There was no interaction between the effects of nicotine and time point on CPA ratings, $F(1, 60) = 0.20, p = .66, \eta^2_G = 0.00073$. Simple main effects showed that the gum group did not have an significant effect on CPA ratings, $F(1, 58) = 2.38, p = .13, \eta^2_G = 0.03$. However there was a significant main effect of time point on CPA ratings, $F(1, 60) = 18.27, p < .001, \eta^2_G = 0.063$.

### 6 Extras for relation between PAF and pain at baseline

#### 6.1 Median split on baseline PAF

As conducted in previous research [1,2], a median split on baseline sensorimotor PAF (median = 9.90) was executed to separate participants into faster and slower PAF groups.

An independent samples t-test showed that mean pain ratings during PHP were significantly higher for those with slower PAF at baseline (M = 6.28 ±2.09) compared with those with faster PAF at baseline (M = 5.04 ±2.45), with anecdotal evidence for this difference, $t(60) = -2.15$, $p = .036$, 95% CI: [-2.40, -0.085], $\text{BF}_{10} = 1.75$ (Figure 11). There was no significant difference in max pain ratings during PHP between those with slower PAF (M = 7.95 ±1.90) and those with faster PAF (M = 6.76 ±2.83), $W = 358.5$, $p = .087$, 95% CI: [-2.24, 0.11] (Figure 11).

Displayed in Figure 11 (plot C), an ANOVA showed main effects of sex ($F(1, 58) = 5.39, p = 0.024, \eta^2_G = 0.085$, $\text{BF}_{10} = 2.11$) and PAF speed group ($F(1, 58) = 5.04, p = 0.029, \eta^2_G = 0.08$, $\text{BF}_{10} = 1.75$) on mean PHP ratings, but without a significant interaction ($F(1, 58) = 0.71, p = 0.40, \eta^2_G = 0.012$, $\text{BF}_{10} = 4.27$). However, Bayesian analysis suggested that there was anecdotal evidence for these main effects, while there was moderate evidence for there being an interaction between sex and PAF group.

Post-hoc t-tests suggest that, within male participants, there was a significant difference in mean PHP ratings between PAF speed groups. Mean PHP ratings for males were higher for those with slower PAF at baseline (M = 5.85 ±2.08) than those with faster PAF (M = 4.13 ±2.33), with anecdotal evidence for this difference, $t(28) = -2.14$, $p = .041$, 95% CI: [-3.38, -0.075], $\text{BF}_{10} = 1.82$. Mean pain ratings during PHP for females were not significantly higher for those with slower PAF at baseline (M = 6.68 ±2.08) than those with faster PAF (M = 5.90 ±2.30), with anecdotal evidence for the lack of difference, $t(30) = -1.01$, $p = .32$, 95% CI: [-2.36, 0.80], $\text{BF}_{10} = 0.50$.

In contrast with PHP, mean pain ratings during CPA were not significantly higher for those with slower PAF at baseline (M = 4.74 ±1.46) compared with those with faster PAF at baseline (M = 4.42 ±1.62), with anecdotal evidence for a lack of difference, $t(60) = -0.83$, $p = .41$, 95% CI: [-1.11, 0.46], $\text{BF}_{10} = 0.35$ (Figure 11, plot B). Max pain ratings during CPA were also not significantly higher for those with slower PAF at baseline (M = 5.90 ±1.78) compared with those with faster PAF at baseline (M = 5.38 ±1.89), with anecdotal evidence for this lack of difference, $t(60) = -1.11$, $p = .27$, 95% CI: [-1.45, 0.41], $\text{BF}_{10} = 0.44$.

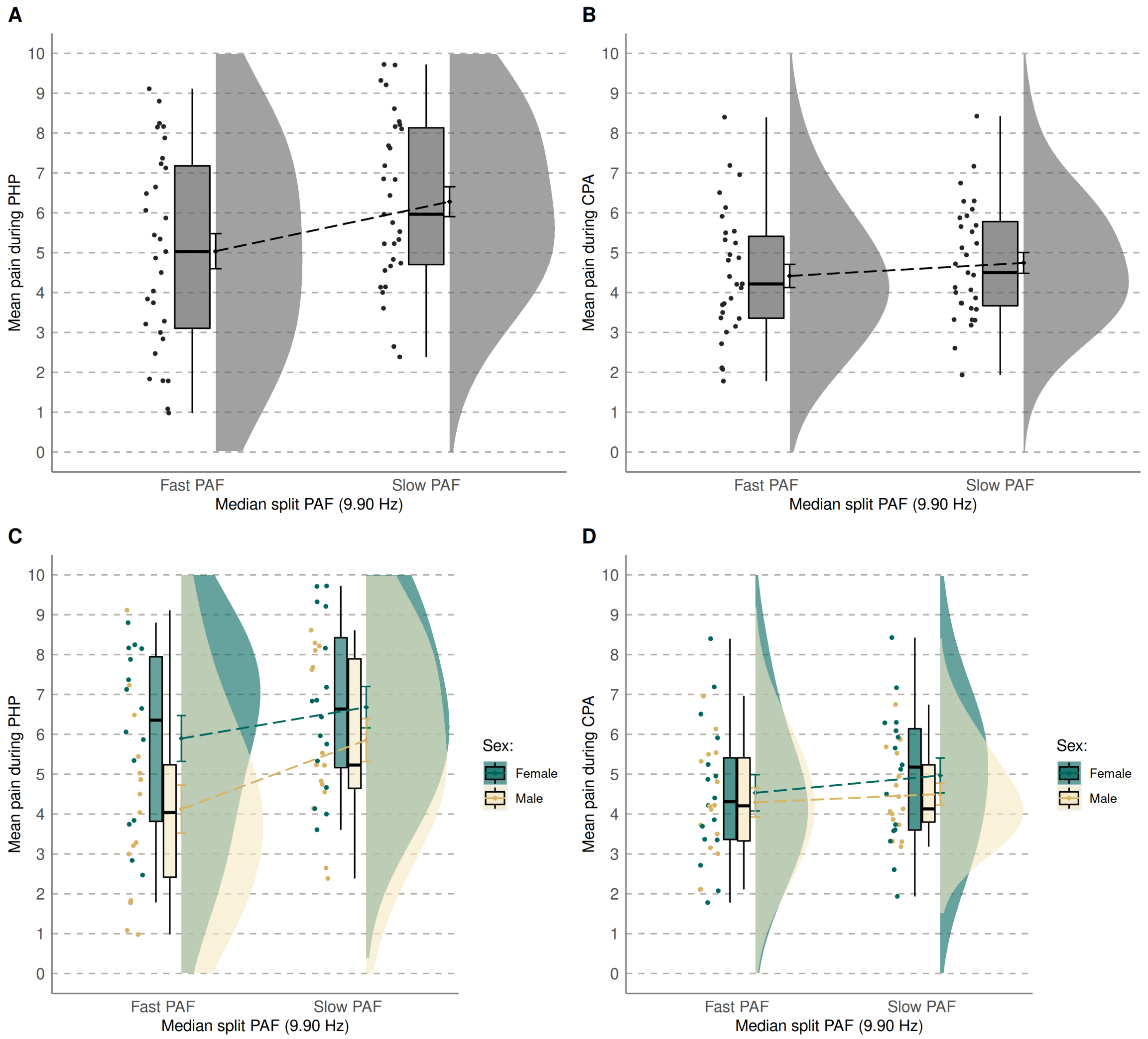

Figure 11: Median split of sensorimotor PAF (median = 9.90 Hz) looking at mean pain during A) phasic heat pain (PHP) and B) cuff pressure algometry (CPA) for the whole sample (N=62), as well as split by sex for C) PHP and D) CPA.

#### 6.2 Correlations between PAF and pain at baseline

Table 28: Spearman correlation coefficients (*p*-values) between PAF calculated using the CoG method and mean pain ratings during prolonged pain tests for all participants (N = 62), and separately in male (n = 30) and female (n = 32) participants.

| | | PHP | | CPA | |
|---|---|---|---|---|---|
| | | | $BF_{10}$ | | $BF_{10}$ |
| Sensorimotor PAF | Narrow (9–11 Hz) | -0.17 (.20) | 0.96 | -0.054 (.67) | 0.32 |
| | *male* | *-0.23 (.22)* | *1.15* | *-0.13 (.50)* | *0.50* |
| | *female* | *-0.18 (.32)* | *0.78* | *-0.061 (.74)* | *0.40* |
| | Wide (8–12 Hz) | -0.15 (.25) | 0.62 | -0.071 (.58) | 0.36 |
| | *male* | *-0.32 (.088)* | *1.96* | *-0.16 (.40)* | *0.53* |
| | *female* | *-0.066 (.72)* | *0.42* | *-0.056 (.76)* | *0.42* |
| Global PAF | Narrow (9–11 Hz) | -0.15 (.25) | 0.58 | -0.062 (.63) | 0.30 |
| | *male* | *-0.32 (.09)* | *1.66* | *-0.21 (.28)* | *0.61* |
| | *female* | *-0.098 (.59)* | *0.49* | *-0.032 (.86)* | *0.39* |
| | Wide (8–12 Hz) | -0.14 (.28) | 0.52 | -0.086 (.50) | 0.33 |
| | *male* | *-0.39 (.032)* | **3.22** | *-0.23 (.23)* | *0.61* |
| | *female* | *-0.032 (.86)* | *0.40* | *-0.086 (.64)* | *0.40* |

*Note. CoG = centre of gravity; PAF = peak alpha frequency; PHP = phasic heat pain; CPA = cuff pressure algometry.

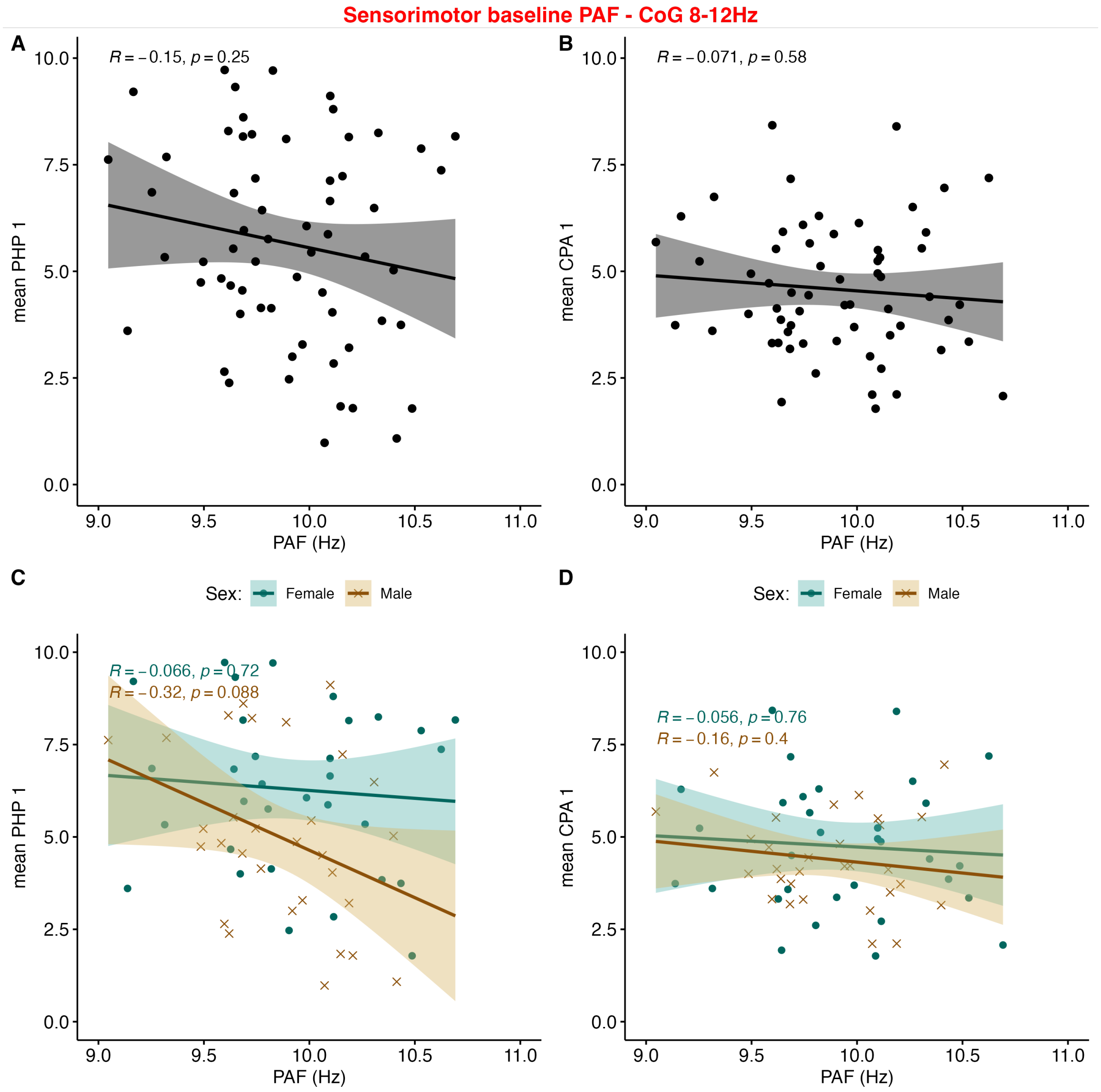

Figure 12: Wide band (8–12 Hz) Sensorimotor peak alpha frequency (PAF) does not correlate with A) mean pain ratings during phasic heat pain (PHP) or B) cuff pressure algometry (CPA). Regression lines and shaded 95% confidence intervals. Correlations separated by sex for females in teal (n = 32) and males in brown (n = 30) for C) PHP and D) CPA, suggest strengthening of a negative relationship between sensorimotor PAF and PHP ratings for males.

***The method used to calculate PAF alters the PAF–pain relationship***

PAF values calculated by the centre of gravity (CoG) or peak picking method were highly correlated for the wide window (8–12 Hz; $rs = 0.99, p < .001$) and the narrow window (9–11 Hz; $rs = 0.99, p < .001$).

Using the peak picking method did not change outcomes for correlations between PAF and mean pain ratings.

Table 29: Spearman correlation coefficients ($p$-values) between baseline PAF calculated with the peak picking method and mean pain ratings during phasic heat pain (PHP) and cuff pressure algometry (CPA) for all participants (N = 62), and separately in male (n = 30) and female (n = 32) groups.

| | | PHP | | CPA | |
|---|---|---|---|---|---|
| | | | $BF_{10}$ | | $BF_{10}$ |
| Sensorimotor PAF | Narrow (9–11 Hz) | -0.12 (.35) | 0.48 | -0.038 (.77) | 0.29 |
| | *male* | *-0.27 (.15)* | *1.41* | *-0.11 (.55)* | *0.43* |
| | *female* | *-0.17 (.36)* | *0.65* | *-0.072 (.70)* | *0.39* |
| | Wide (8–12 Hz) | -0.15 (.26) | 0.56 | -0.063 (.62) | 0.30 |
| | *male* | *-0.38 (.038)* | **2.37** | *-0.14 (.46)* | *0.42* |
| | *female* | *-0.051 (.78)* | *0.42* | *-0.040 (.83)* | *0.41* |
| Global PAF | Narrow (9–11 Hz) | -0.12 (.35) | 0.48 | -0.038 (.77) | 0.29 |
| | *male* | *-0.31 (.099)* | *1.84* | *-0.20 (.30)* | *0.52* |
| | *female* | *-0.069 (.71)* | *0.43* | *-0.016 (.93)* | *0.40* |
| | Wide (8–12 Hz) | -0.15 (.26) | 0.56 | -0.063 (.63) | 0.30 |
| | *male* | -0.42 (.022)* | **3.92** | *-0.18 (.35)* | *0.46* |
| | *female* | *-0.011 (.95)* | *0.39* | *-0.054 (.77)* | *0.40* |

***Using max rather than mean PAF does not alter the PAF–pain relationship***

A similar pattern was seen for correlations between global PAF (8–12 Hz) and max pain during PHP (Table 30), with no relationship for the whole sample ($rs = -0.052, p = .69$, $BF_{10} = 0.40$), a stronger and significant negative relationship for male participants alone ($rs = -0.39, p = .036$, $BF_{10}$= 2.71), and no relationship for females only ($rs = -0.073, p = .69$, $BF_{10} = 0.39$).

Table 30: Spearman correlation coefficients ($p$-values) between baseline PAF and max pain ratings during phasic heat pain (PHP) and cuff pressure algometry (CPA) for all participants (N = 62), and separately in male (n = 30) and female (n = 32) groups.

| | | PHP | | CPA | |
|---|---|---|---|---|---|
| | | | $BF_{10}$ | | $BF_{10}$ |
| Sensorimotor PAF | Narrow (9–11 Hz) | -0.096 (.46) | 0.66 | -0.065 (.62) | 0.33 |
| | *male* | *-0.24 (.0.20)* | *1.21* | *-0.13 (.48)* | *0.61* |
| | *female* | *-0.11 (.57)* | *0.56* | *-0.043 (.82)* | *0.39* |
| | Wide (8–12 Hz) | -0.051 (.69) | 0.39 | -0.094 (.47) | 0.41 |
| | *male* | *-0.29 (.13)* | *1.53* | *-0.15 (.44)* | *0.57* |
| | *female* | *-0.054 (.77)* | *0.39* | *-0.078 (.67)* | *0.44* |
| Global PAF | Narrow (9–11 Hz) | -0.086 (.51) | 0.54 | -0.082 (.53) | 0.32 |
| | *male* | *-0.33 (.071)* | *1.95* | *-0.23 (.22)* | *0.85* |
| | *female* | *-0.020 (.91)* | *0.46* | *-0.015 (.93)* | *0.39* |
| | Wide (8–12 Hz) | -0.052 (.69) | 0.40 | -0.12 (.36) | 0.40 |
| | *male* | *-0.39 (.036)* | **2.71** | *-0.24 (.20)* | *0.73* |
| | *female* | *-0.073 (.69)* | *0.39* | *-0.11 (.55)* | *0.42* |

[Median split of sensorimotor PAF (median = 9.90 Hz) looking at mean pain during A) phasic heat pain (PHP) and B) cuff pressure algometry (CPA) for the whole sample (N=62), as well as split by sex for C) PHP and D) CPA.

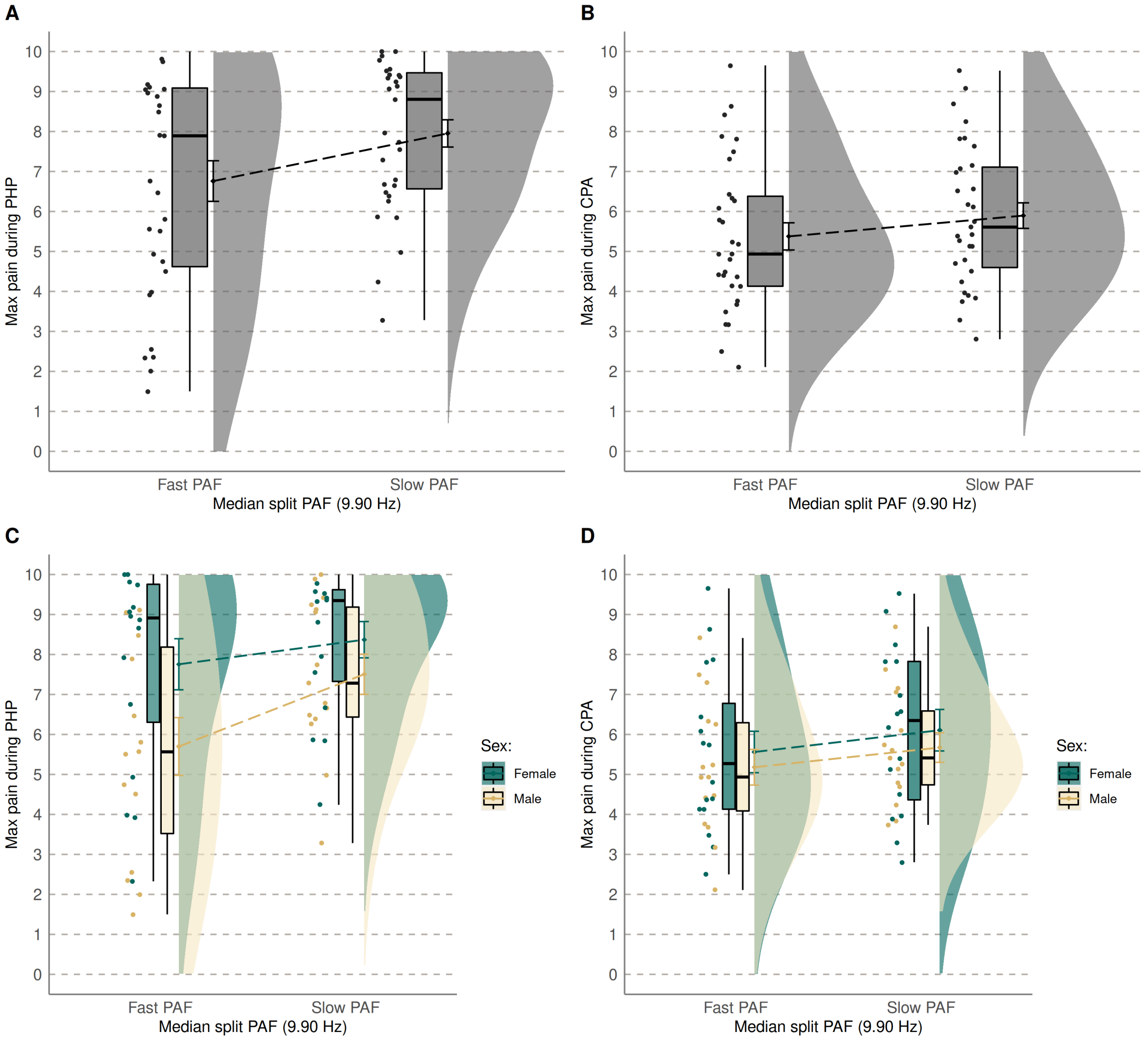

Figure 13: Median split of sensorimotor PAF (median = 9.90 Hz) looking at max pain during A) phasic heat pain (PHP) and B) cuff pressure algometry (CPA) for the whole sample (N=62), as well as split by sex for C) PHP and D) CPA.

***Using AUC rather than mean PAF does not alter the PAF–pain relationship***

Table 31: Spearman correlation coefficients ($p$-values) between PAF and AUC values calculated for the whole duration of the prolonged pain tests for all participants (N = 62), and separately in male (n = 30) and female (n = 32) groups. Phasic heat pain (PHP) and cuff pressure algometry (CPA).

| | | PHP | | CPA | |
|---|---|---|---|---|---|
| | | | $BF_{10}$ | | $BF_{10}$ |
| Sensorimotor PAF | Narrow (9–11 Hz) | -0.16 (.22) | 0.90 | -0.053 (.68) | 0.32 |
| | *male* | *-0.19 (.31)* | *1.27* | *-0.13 (.50)* | *0.50* |
| | *female* | *-0.24 (.18)* | *0.65* | *-0.061 (.74)* | *0.40* |
| | Wide (8–12 Hz) | -0.15 (.24) | 0.63 | -0.070 (.59) | 0.36 |
| | *male* | *-0.29 (.12)* | *1.47* | *-0.16 (.40)* | *0.53* |
| | *female* | *-0.12 (.50)* | *0.44* | *-0.056 (.76)* | *0.42* |
| Global PAF | Narrow (9–11 Hz) | -0.14 (.28) | 0.54 | -0.061 (.64) | 0.30 |
| | *male* | *-0.27 (.15)* | *1.07* | *-0.21 (.28)* | *0.61* |
| | *female* | *-0.16 (.38)* | *0.52* | *-0.032 (.86)* | *0.39* |
| | Wide (8–12 Hz) | -0.14 (.28) | 0.51 | -0.086 (.51) | 0.33 |
| | *male* | *-0.36 (.053)* | **2.14** | *-0.23 (.23)* | *0.61* |
| | *female* | *-0.091 (.62)* | *0.41* | *-0.086 (.64)* | *0.40* |

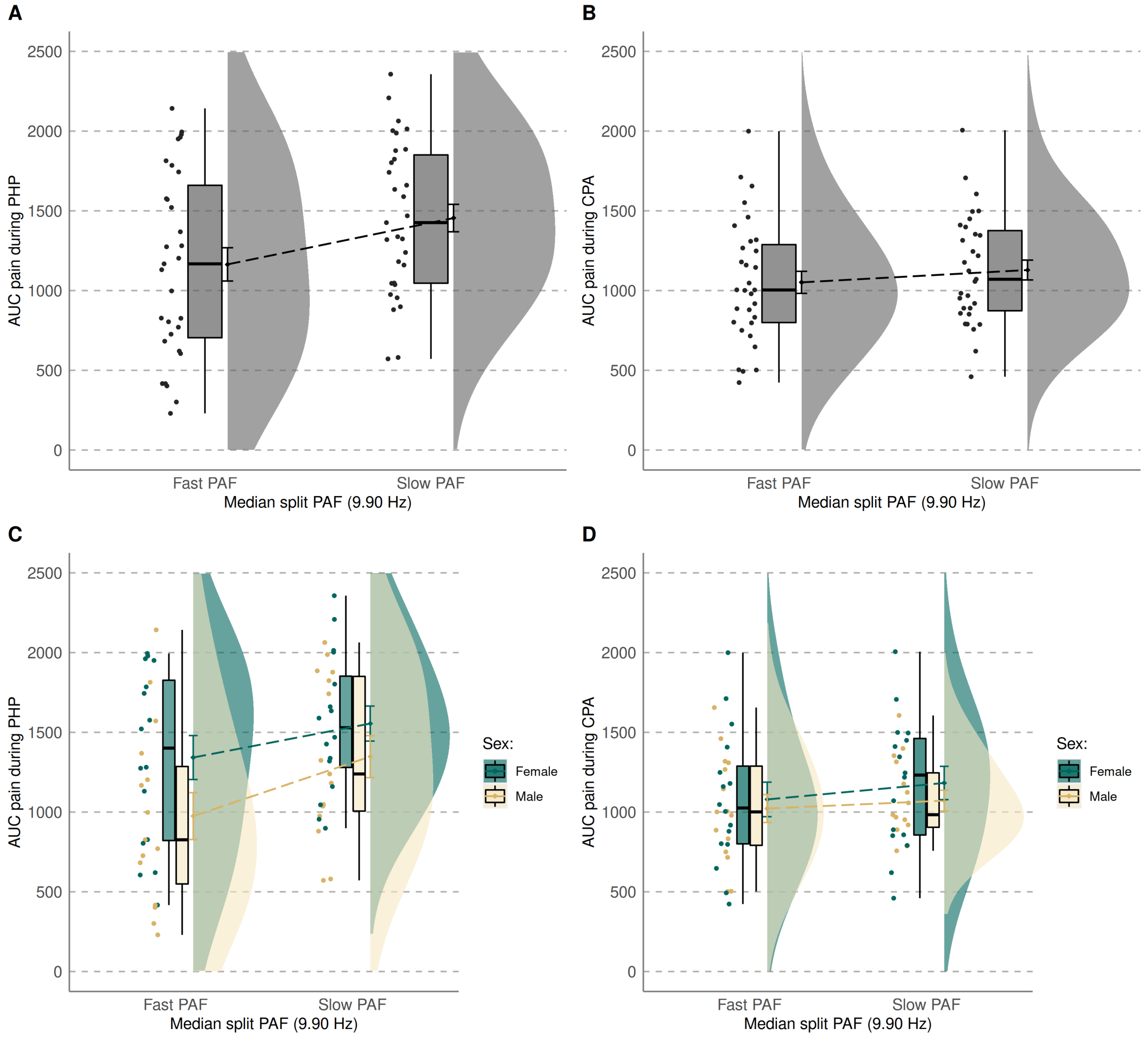

Figure 14: Median split of sensorimotor PAF (median = 9.90 Hz) looking at the AUC for full duration - unstandardised during A) phasic heat pain (PHP) and B) cuff pressure algometry (CPA) for the whole sample (N=62), as well as split by sex for C) PHP and D) CPA.

#### 6.3 Removing five participants who received phasic heat pain at 45 °C does not alter the PAF–pain relationship

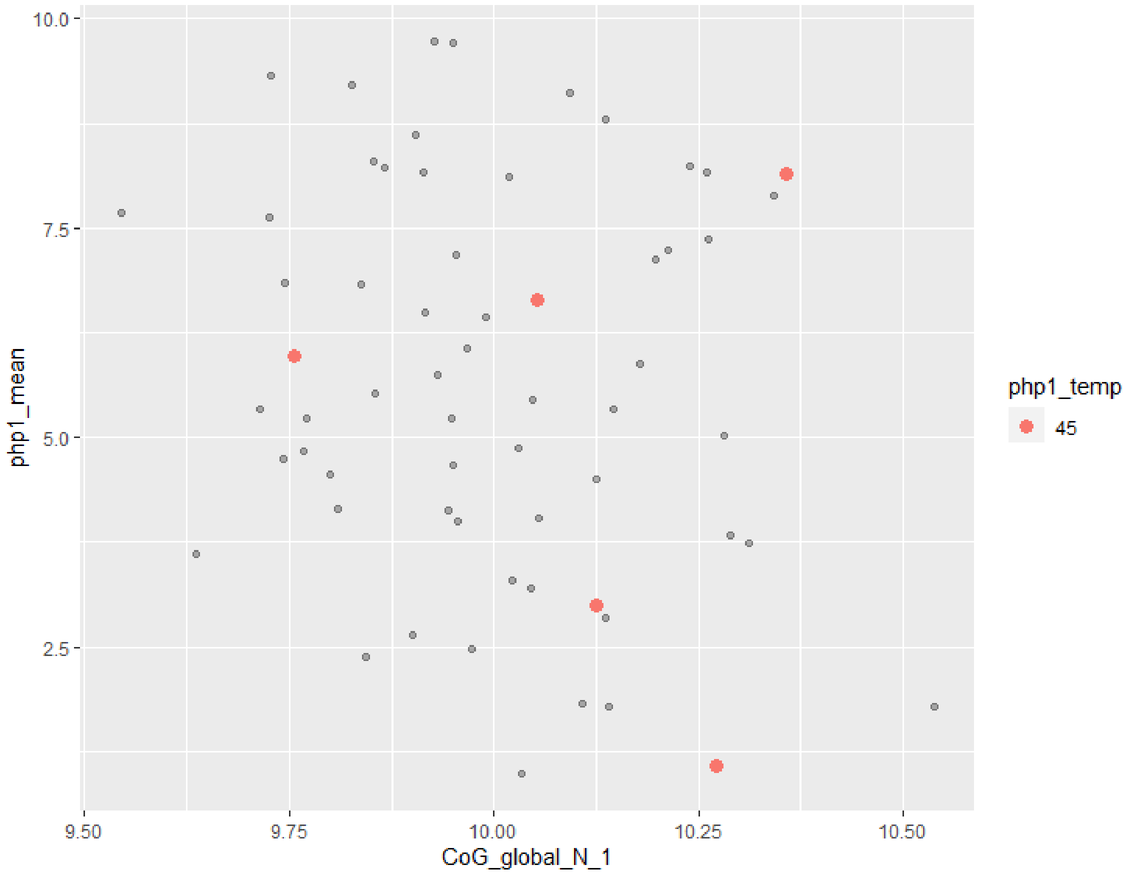

Figure 15: Display of five participant who requested a lower temperature (i.e. 45 °C) as they could not tolerate phasic heat pain delivered at 46 °C. Removing these five participants did not influence the relationship between PAF and pain.

### 7 Methods

#### 7.1 Pre-determined confounding selection (DAGS)

Directed acyclic graphs (DAG) were constructed using dagitty.net [3]; the paths represent known causal effects that were assumed based on prior knowledge [4]. Confounding variables were chosen based on these DAGs (Figure 16 and Figure 17), and were defined as any measured variables that are involved in alternative paths between the mediator and outcome [4].

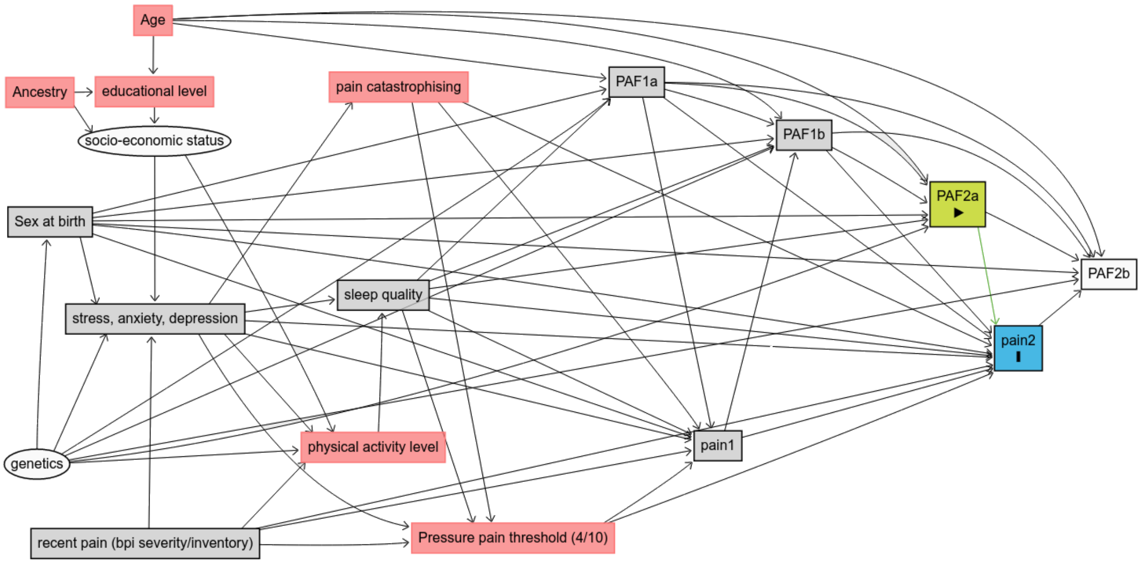

Figure 16: Directed acyclic graphic depicting the assumed mechanisms in the mediator–outcome relationship for cuff pressure algometry. $PAF_{1a}$, $pain_1$, and $PAF_{1b}$ were measured prior to nicotine intervention, while $PAF_{2a}$, $pain_2$, and $PAF_{2b}$ were measured post-intervention. Grey squares were those assessed as confounding variables. *Note. PAF = peak alpha frequency.*

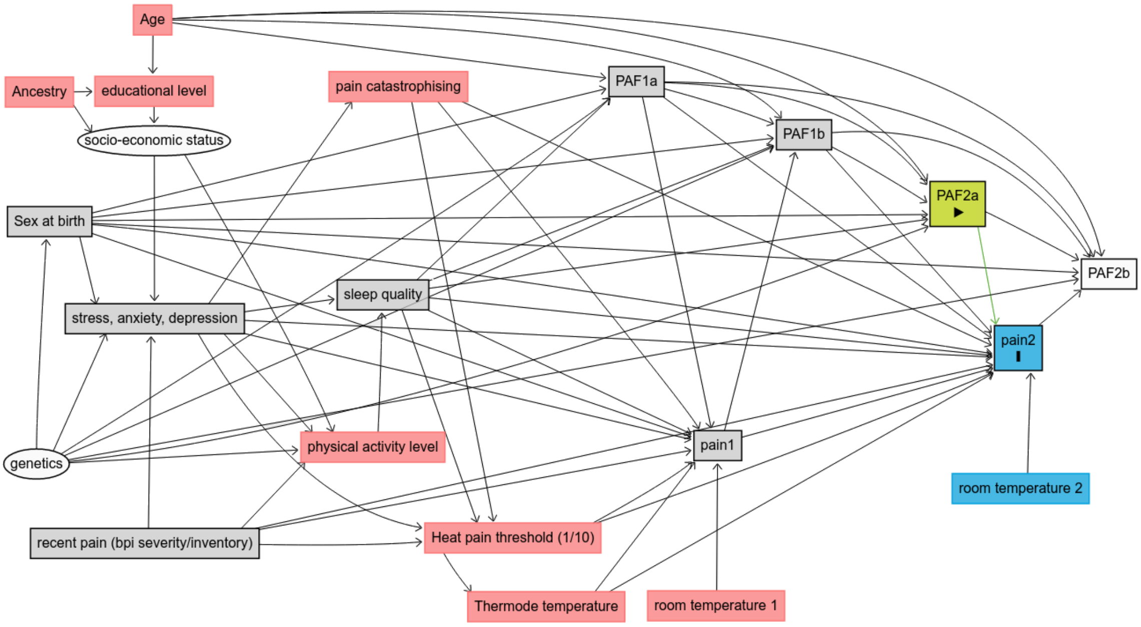

Figure 17: Directed acyclic graphic depicting the assumed mechanisms in the mediator–outcome relationship for phasic heat pain. $\mathrm{PAF}_{1a}$, $\mathrm{pain}_1$, and $\mathrm{PAF}_{1b}$ were measured prior to nicotine intervention, while $\mathrm{PAF}_{2a}$, $\mathrm{pain}_2$, and $\mathrm{PAF}_{2b}$ were measured post-intervention. Grey squares were those assessed as confounding variables. *Note. PAF = peak alpha frequency.*

#### 7.2 `PROCESS` results similar to difference score 2W-LCS model

***Phasic heat pain***

Conducting mediation analysis using `PROCESS` indicates total and direct effects of gum group on post-gum PHP ratings, with no indirect effect via post-gum PAF values. There were effects of gum group, sex, and pre-gum PAF on post-gum PAF, similar to the 2W-LCS model reported in the manuscript. For PHP there are also effects of gum and pain ratings pre-gum on pain ratings post-gum, similar to the 2W-LCS model, however, there is no additional effect of stress. Therefore, the analysis conducted in `PROCESS` supports the 2W-LCS mediation analysis conducted in `lavaan`.

******************** PROCESS for R Version 4.2 beta ****************

`Written by Andrew F. Hayes, Ph.D.  www.afhayes.com`

Documentation available in Hayes (2022). www.guilford.com/p/hayes3

---

Model : 4 Y : y2 X : x M : m2

Covariates: m1 y1 cov1 cov2 cov3 cov4 cov5 cov6

Sample size: 60

Custom seed: 2022

---

Outcome Variable: m2

Model Summary:

| R | R-sq | MSE | F | df1 | df2 | p |
|---|---|---|---|---|---|---|
| 0.96 | 0.91 | 0.02 | 57.67 | 9 | 50 | 0.00 |

Model:

| | coeff | se | t | p | LLCI | ULCI |
|---|---|---|---|---|---|---|
| constant | 0.0423 | 0.5083 | 0.0832 | 0.9340 | -0.9786 | 1.0632 |
| x | 0.0918 | 0.0397 | 2.3118 | 0.0249 | 0.0120 | 0.1715 |
| m1 | 1.0167 | 0.0479 | 21.2229 | 0.0000 | 0.9204 | 1.1129 |
| y1 | -0.0102 | 0.0086 | -1.1752 | 0.2455 | -0.0275 | 0.0072 |
| cov1 | -0.0889 | 0.0405 | -2.1964 | 0.0327 | -0.1703 | -0.0076 |
| cov2 | -0.0348 | 0.0241 | -1.4432 | 0.1552 | -0.0832 | 0.0136 |

| | coeff | se | t | p | LLCI | ULCI |
|---|---|---|---|---|---|---|
| cov3 | 0.0340 | 0.0217 | 1.5681 | 0.1232 | -0.0095 | 0.0774 |
| cov4 | -0.0052 | 0.0052 | -0.9868 | 0.3285 | -0.0157 | 0.0053 |
| cov5 | -0.0144 | 0.0107 | -1.3473 | 0.1840 | -0.0358 | 0.0071 |
| cov6 | 0.0177 | 0.0239 | 0.7438 | 0.4605 | -0.0302 | 0.0657 |

---

Outcome Variable: y2

Model Summary:

| R | R-sq | MSE | F | df1 | df2 | p |
|---|---|---|---|---|---|---|
| 0.94 | 0.90 | 0.74 | 44.60 | 10 | 49 | 0.00 |

Model:

| | coeff | se | t | p | LLCI | ULCI |
|---|---|---|---|---|---|---|
| constant | 1.6495 | 3.1271 | 0.5275 | 0.6002 | -4.6346 | 7.9336 |
| x | -0.6845 | 0.2569 | -2.6645 | 0.0104 | -1.2008 | -0.1683 |
| m2 | 0.2878 | 0.8700 | 0.3308 | 0.7422 | -1.4606 | 2.0362 |
| m1 | -0.3215 | 0.9323 | -0.3448 | 0.7317 | -2.1950 | 1.5521 |
| y1 | 1.0189 | 0.0539 | 18.8984 | 0.0000 | 0.9106 | 1.1273 |
| cov1 | 0.1653 | 0.2609 | 0.6336 | 0.5293 | -0.3590 | 0.6895 |
| cov2 | 0.3025 | 0.1513 | 1.9998 | 0.0511 | -0.0015 | 0.6066 |
| cov3 | -0.0713 | 0.1364 | -0.5226 | 0.6036 | -0.3455 | 0.2029 |
| cov4 | -0.0940 | 0.0325 | -2.8945 | 0.0057 | -0.1593 | -0.0287 |
| cov5 | -0.0068 | 0.0668 | -0.1019 | 0.9193 | -0.1410 | 0.1274 |
| cov6 | -0.1742 | 0.1476 | -1.1806 | 0.2434 | -0.4707 | 0.1223 |

************************ TOTAL EFFECT MODEL ************************** Outcome Variable: y2

Model Summary:

| R | R-sq | MSE | F | df1 | df2 | p |
|---|---|---|---|---|---|---|
| 0.95 | 0.90 | 0.73 | 50.44 | 9 | 50 | 0.00 |

Model:

| | coeff | se | t | p | LLCI | ULCI |
|---|---|---|---|---|---|---|
| constant | 1.6617 | 3.0989 | 0.5362 | 0.5942 | -4.5626 | 7.8860 |
| x | -0.6581 | 0.2420 | -2.7195 | 0.0090 | -1.1442 | -0.1720 |
| m1 | -0.0289 | 0.2921 | -0.0988 | 0.9217 | -0.6155 | 0.5578 |
| y1 | 1.0160 | 0.0527 | 19.2751 | 0.0000 | 0.9101 | 1.1219 |
| cov1 | 0.1397 | 0.2469 | 0.5658 | 0.5741 | -0.3562 | 0.6356 |
| cov2 | 0.2925 | 0.1469 | 1.9913 | 0.0519 | -0.0025 | 0.5876 |
| cov3 | -0.0615 | 0.1320 | -0.4661 | 0.6432 | -0.3267 | 0.2036 |
| cov4 | -0.0955 | 0.0319 | -2.9955 | 0.0043 | -0.1596 | -0.0315 |
| cov5 | -0.0109 | 0.0650 | -0.1683 | 0.8671 | -0.1416 | 0.1197 |
| cov6 | -0.1691 | 0.1454 | -1.1628 | 0.2504 | -0.4612 | 0.1230 |

---

Bootstrapping progress: |»»»»»»»»»»»»»»»»»»»»»»»»»»»»»»»»»| 100%

************ TOTAL, DIRECT, AND INDIRECT EFFECTS OF X ON Y ************

| | effect | se | t | p | LLCI | ULCI | c_ps |
|---|---|---|---|---|---|---|---|
| Total effect of X on Y: | -0.6581 | 0.2420 | -2.7195 | 0.0090 | -1.1442 | -0.1720 | -0.2634 |
| | effect | se | t | p | LLCI | ULCI | c'_ps |
| Direct effect of X on Y: | -0.6845 | 0.2569 | -2.6645 | 0.0104 | -1.2008 | -0.1683 | -0.2740 |
| | Effect | BootSE | BootLLCI | BootULCI | | | |
| Indirect effect(s) of X on Y: | | | | | | | |
| m2 | 0.0264 | 0.0897 | -0.1075 | 0.2641 | | | |
| Partially standardized indirect effect(s) of X on Y: | | | | | | | |
| m2 | 0.0106 | 0.0366 | -0.0438 | 0.1108 | | | |

********** BOOTSTRAP RESULTS FOR REGRESSION MODEL PARAMETERS **********

Outcome variable: m2

| | Coeff | BootMean | BootSE | BootLLCI | BootULCI |
|---|---|---|---|---|---|
| constant | 0.0423 | 0.0549 | 0.4596 | -0.8006 | 1.0544 |
| x | 0.0918 | 0.0907 | 0.0392 | 0.0139 | 0.1622 |
| m1 | 1.0167 | 1.0166 | 0.0447 | 0.9269 | 1.1023 |
| y1 | -0.0102 | -0.0110 | 0.0100 | -0.0321 | 0.0075 |
| cov1 | -0.0889 | -0.0964 | 0.0419 | -0.1801 | -0.0178 |
| cov2 | -0.0348 | -0.0349 | 0.0289 | -0.0825 | 0.0329 |
| cov3 | 0.0340 | 0.0338 | 0.0234 | -0.0129 | 0.0874 |
| cov4 | -0.0052 | -0.0053 | 0.0056 | -0.0164 | 0.0051 |
| cov5 | -0.0144 | -0.0144 | 0.0135 | -0.0384 | 0.0151 |
| cov6 | 0.0177 | 0.0156 | 0.0235 | -0.0288 | 0.0625 |

Outcome variable: y2

| | Coeff | BootMean | BootSE | BootLLCI | BootULCI |
|---|---|---|---|---|---|
| constant | 1.6495 | 1.4241 | 2.9174 | -4.7156 | 7.5110 |
| x | -0.6845 | -0.6922 | 0.2979 | -1.3008 | -0.1306 |
| m2 | 0.2878 | 0.4203 | 0.8565 | -1.3355 | 2.0972 |
| m1 | -0.3215 | -0.4425 | 0.8887 | -2.0953 | 1.3635 |
| y1 | 1.0189 | 1.0223 | 0.0513 | 0.9266 | 1.1326 |
| cov1 | 0.1653 | 0.1970 | 0.3173 | -0.4142 | 0.8529 |
| cov2 | 0.3025 | 0.3223 | 0.1836 | -0.0705 | 0.6530 |
| cov3 | -0.0713 | -0.1148 | 0.1665 | -0.4878 | 0.1277 |
| cov4 | -0.0940 | -0.0883 | 0.0500 | -0.1818 | 0.0055 |
| cov5 | -0.0068 | -0.0018 | 0.0710 | -0.1536 | 0.1330 |
| cov6 | -0.1742 | -0.1660 | 0.1271 | -0.4183 | 0.0950 |

******************** ANALYSIS NOTES AND ERRORS ************************

Level of confidence for all confidence intervals in output: 95

Number of bootstraps for percentile bootstrap confidence intervals: 1000

NOTE: Some cases with missing data were deleted. The number of deleted cases was: 2 >

##### ***Cuff pressure algometry***

Conducting mediation analysis using `PROCESS` did not indicate total, direct, or indirect effects of gum group on CPA ratings. There were effects of gum group, sex, and pre-gum PAF on post-gum PAF, similar to the 2W-LCS model reported in the manuscript . There were no effects of gum group on CPA ratings, only a significant effect of pain ratings pre-gum on the pain ratings post-gum. Therefore, the analysis conducted in `PROCESS` supports the 2W-LCS mediation analysis conducted in `lavaan`.

******************** PROCESS for R Version 4.2 beta ****************

`Written by Andrew F. Hayes, Ph.D.  www.afhayes.com`

Documentation available in Hayes (2022). www.guilford.com/p/hayes3

---

Model : 4 Y : y2 X : x M : m2

Covariates: m1 y1 cov1 cov2 cov3 cov4 cov5 cov6

Sample size: 62

Custom seed: 2022

---

Outcome Variable: m2

Model Summary:

| R | R-sq | MSE | F | df1 | df2 | p |
|---|---|---|---|---|---|---|
| 0.96 | 0.91 | 0.02 | 59.29 | 9 | 52 | 0.00 |

Model:

| | coeff | se | t | p | LLCI | ULCI |
|---|---|---|---|---|---|---|
| constant | -0.1394 | 0.4936 | -0.2823 | 0.7788 | -1.1298 | 0.8511 |
| x | 0.0939 | 0.0391 | 2.3978 | 0.0201 | 0.0153 | 0.1724 |
| m1 | 1.0274 | 0.0468 | 21.9582 | 0.0000 | 0.9336 | 1.1213 |
| y1 | -0.0002 | 0.0127 | -0.0185 | 0.9853 | -0.0258 | 0.0253 |
| cov1 | -0.0722 | 0.0379 | -1.9024 | 0.0627 | -0.1483 | 0.0040 |
| cov2 | -0.0311 | 0.0238 | -1.3077 | 0.1967 | -0.0789 | 0.0166 |
| cov3 | 0.0292 | 0.0212 | 1.3755 | 0.1749 | -0.0134 | 0.0717 |
| cov4 | -0.0044 | 0.0052 | -0.8470 | 0.4009 | -0.0147 | 0.0060 |

| | coeff | se | t | p | LLCI | ULCI |
|---|---|---|---|---|---|---|
| cov5 | -0.0138 | 0.0107 | -1.2982 | 0.2000 | -0.0352 | 0.0075 |
| cov6 | 0.0175 | 0.0233 | 0.7493 | 0.4571 | -0.0293 | 0.0643 |

Outcome Variable: y2

Model Summary:

| R | R-sq | MSE | F | df1 | df2 | p |
|---|---|---|---|---|---|---|
| 0.67 | 0.46 | 2.42 | 4.3 | 10 | 51 | 0.0002 |

Model:

| | coeff | se | t | p | LLCI | ULCI |
|---|---|---|---|---|---|---|
| constant | -9.4138 | 5.4985 | -1.7121 | 0.0930 | -20.4527 | 1.6251 |
| x | 0.1508 | 0.4592 | 0.3283 | 0.7440 | -0.7712 | 1.0727 |
| m2 | -1.5230 | 1.5436 | -0.9866 | 0.3285 | -4.6220 | 1.5760 |
| m1 | 2.4601 | 1.6693 | 1.4737 | 0.1467 | -0.8912 | 5.8114 |
| y1 | 0.7949 | 0.1419 | 5.6020 | 0.0000 | 0.5100 | 1.0798 |
| cov1 | 0.7441 | 0.4366 | 1.7042 | 0.0944 | -0.1325 | 1.6206 |
| cov2 | 0.1110 | 0.2694 | 0.4119 | 0.6821 | -0.4299 | 0.6518 |
| cov3 | -0.1441 | 0.2402 | -0.6001 | 0.5511 | -0.6264 | 0.3381 |
| cov4 | -0.0620 | 0.0578 | -1.0733 | 0.2882 | -0.1781 | 0.0540 |
| cov5 | 0.0462 | 0.1205 | 0.3831 | 0.7032 | -0.1957 | 0.2881 |
| cov6 | 0.0783 | 0.2612 | 0.2999 | 0.7654 | -0.4461 | 0.6028 |

************************ TOTAL EFFECT MODEL ************************** Outcome Variable: y2

Model Summary:

| R | R-sq | MSE | F | df1 | df2 | p |
|---|---|---|---|---|---|---|
| 0.67 | 0.45 | 2.42 | 4.67 | 9 | 52 | 0.0001 |

Model:

| | coeff | se | t | p | LLCI | ULCI |
|---|---|---|---|---|---|---|
| constant | -9.2015 | 5.4929 | -1.6752 | 0.0999 | -20.2240 | 1.8209 |
| x | 0.0078 | 0.4357 | 0.0179 | 0.9858 | -0.8665 | 0.8820 |
| m1 | 0.8953 | 0.5207 | 1.7194 | 0.0915 | -0.1496 | 1.9402 |
| y1 | 0.7953 | 0.1419 | 5.6060 | 0.0000 | 0.5106 | 1.0800 |

| | coeff | se | t | p | LLCI | ULCI |
|---|---|---|---|---|---|---|
| cov1 | 0.8540 | 0.4221 | 2.0233 | 0.0482 | 0.0070 | 1.7009 |
| cov2 | 0.1584 | 0.2650 | 0.5977 | 0.5526 | -0.3734 | 0.6902 |
| cov3 | -0.1885 | 0.2359 | -0.7993 | 0.4278 | -0.6619 | 0.2848 |
| cov4 | -0.0554 | 0.0574 | -0.9650 | 0.3390 | -0.1706 | 0.0598 |
| cov5 | 0.0672 | 0.1186 | 0.5671 | 0.5731 | -0.1707 | 0.3051 |
| cov6 | 0.0517 | 0.2598 | 0.1991 | 0.8430 | -0.4695 | 0.5730 |

---

Bootstrapping progress: |»»»»»»»»»»»»»»»»»»»»»»»»»»»»»»»»»»»| 100%

************ TOTAL, DIRECT, AND INDIRECT EFFECTS OF X ON Y ************

| | effect | se | t | p | LLCI | ULCI | c_ps |
|---|---|---|---|---|---|---|---|
| Total effect of X on Y: | 0.0078 | 0.4357 | 0.0179 | 0.9858 | -0.8665 | 0.8820 | 0.0040 |
| | effect | se | t | p | LLCI | ULCI | c'_ps |
| Direct effect of X on Y: | 0.1508 | 0.4592 | 0.3283 | 0.7440 | -0.7712 | 1.0727 | 0.0781 |
| | Effect | BootSE | BootLLCI | BootULCI | | | |
| Indirect effect(s) of X on Y: | | | | | | | |
| m2 | -0.1430 | 0.1932 | -0.6166 | 0.1472 | | | |
| Partially standardized indirect effect(s) of X on Y: | | | | | | | |
| m2 | -0.0741 | 0.1008 | -0.3182 | 0.0783 | | | |

********** BOOTSTRAP RESULTS FOR REGRESSION MODEL PARAMETERS **********

Outcome variable: m2

| | Coeff | BootMean | BootSE | BootLLCI | BootULCI |
|---|---|---|---|---|---|
| constant | -0.1394 | -0.1638 | 0.4179 | -0.9571 | 0.6746 |
| x | 0.0939 | 0.0896 | 0.0363 | 0.0174 | 0.1631 |
| m1 | 1.0274 | 1.0298 | 0.0441 | 0.9422 | 1.1167 |
| y1 | -0.0002 | 0.0003 | 0.0125 | -0.0240 | 0.0265 |

| | Coeff | BootMean | BootSE | BootLLCI | BootULCI |
|---|---|---|---|---|---|
| cov1 | -0.0722 | -0.0762 | 0.0378 | -0.1569 | -0.0057 |
| cov2 | -0.0311 | -0.0303 | 0.0260 | -0.0737 | 0.0314 |
| cov3 | 0.0292 | 0.0273 | 0.0226 | -0.0251 | 0.0698 |
| cov4 | -0.0044 | -0.0045 | 0.0057 | -0.0164 | 0.0057 |
| cov5 | -0.0138 | -0.0130 | 0.0133 | -0.0373 | 0.0155 |
| cov6 | 0.0175 | 0.0150 | 0.0228 | -0.0314 | 0.0569 |

Outcome variable: y2

| | Coeff | BootMean | BootSE | BootLLCI | BootULCI |
|---|---|---|---|---|---|
| constant | -9.4138 | -8.9254 | 5.9510 | -20.4027 | 2.8342 |
| x | 0.1508 | 0.1129 | 0.4571 | -0.7853 | 1.0171 |
| m2 | -1.5230 | -1.8158 | 1.8246 | -5.5058 | 1.6244 |
| m1 | 2.4601 | 2.7083 | 1.9373 | -0.9951 | 6.6085 |
| y1 | 0.7949 | 0.8225 | 0.1684 | 0.4745 | 1.1366 |
| cov1 | 0.7441 | 0.6827 | 0.4343 | -0.1467 | 1.5617 |
| cov2 | 0.1110 | 0.1190 | 0.3290 | -0.6042 | 0.6979 |
| cov3 | -0.1441 | -0.2239 | 0.3761 | -1.0676 | 0.3521 |
| cov4 | -0.0620 | -0.0628 | 0.0612 | -0.1900 | 0.0566 |
| cov5 | 0.0462 | 0.0353 | 0.1298 | -0.1944 | 0.3072 |
| cov6 | 0.0783 | 0.0857 | 0.2569 | -0.3816 | 0.6146 |

******************** ANALYSIS NOTES AND ERRORS ***********************

Level of confidence for all confidence intervals in output: 95

Number of bootstraps for percentile bootstrap confidence intervals: 1000
